# Correlated evolution of large DNA fragments in the 3D genome of *Arabidopsis thaliana*

**DOI:** 10.1101/759589

**Authors:** Yubin Yan, Zhaohong Li, Ye Li, Zefeng Wu, Ruolin Yang

**Author notes:** These authors contributed equally to this work (Yubin Yan, Zhaohong Li).

## Abstract

In eukaryotes, the three-dimensional (3D) conformation of the genome is far from random, and this nonrandom chromatin organization is strongly correlated with gene expression and protein function, which are two critical determinants of the selective constraints and evolutionary rates of genes. However, whether genes and other elements that are located close to each other in the 3D genome evolve in a coordinated way has not been investigated in any organism. To address this question, we constructed chromatin interaction networks (CINs) in *Arabidopsis thaliana* based on high-throughput chromosome conformation capture (Hi-C) data and demonstrated that adjacent large DNA fragments in the CIN indeed exhibit more similar levels of polymorphism and evolutionary rates than random fragment pairs. Using simulations that account for the linear distance between fragments, we proved that the 3D chromosomal organization plays a role in the observed correlated evolution. Spatially interacting fragments also exhibit more similar mutation rates and functional constraints in both coding and noncoding regions than the random expectations, indicating that the correlated evolution between 3D neighbors is a result of combined evolutionary forces. A collection of 39 genomic and epigenomic features can explain much of the variance in genetic diversity and evolutionary rates across the genome. Moreover, features that have a greater effect on the evolution of regional sequences tend to show higher similarity between neighboring fragments in the CIN, suggesting a pivotal role of epigenetic modifications and chromatin organization in determining the correlated evolution of large DNA fragments in the 3D genome.

## Introduction

At all levels of the hierarchical organization of living systems, biological units rarely evolve in isolation. For instance, species interact with each other in various ways, such as parasitism, competition, and mutualism. Within organisms, proteins encoded by different genes of the same genome or different genomes (i.e., the nuclear and cytoplasmic genomes) frequently act together to accomplish specific functions. When two or more evolving units interact and produce reciprocal evolutionary changes in each other, we say there is coevolution between them (Carmona et al. 2015). At the species level, a change in one species may impose evolutionary pressure on its interacting species, giving rise to reciprocal modifications in these organisms, as observed in host-pathogen and pollinator/herbivore-plant coevolution (Ehrlich and Raven 1964; Burdon and Thrall 2009). At the molecular level, a mutation that disrupts the structure and function of a protein or a tRNA may be compensated by a second mutation in the same molecule, resulting in no deleterious effect of the first mutation (Kondrashov et al. 2002; Kern and Kondrashov 2004). Such compensatory mutations can occur between interacting molecules as well, leading to molecular coevolution between different molecules (Pazos and Valencia 2008).

Compensatory coevolution at the molecular level may result in correlated evolutionary rates and similar phylogenetic profiles between interacting proteins. Many methods have been developed for detecting protein coevolution based on principles of molecular phylogenetics, and the coevolutionary pattern between residues has been used to predict protein structure and novel interactions [see de Juan et al. (2013) for a review]. In addition to the similarity in phylogenetic profiles, correlated evolutionary rates have been regarded as one of the consequences of compensatory coevolution and have been widely observed between interacting proteins. For example, accelerated rates of evolution in nuclear genes that cooperate with mitochondrial genes have been reported in many animal species (Osada and Akashi 2012; Barreto and Burton 2013; Parmakelis et al. 2013; Barreto et al. 2018), possibly as a result of the high mutation rate in the mitochondrial genome and the accumulation of compensatory mutations in the nuclear genes (Rand et al. 2004; Levin et al. 2014; Sloan et al. 2018). Such accelerated compensatory coevolution has been observed in some plants with elevated mutation rates in the plastid genome as well (Zhang et al. 2015; Weng et al. 2016). The correlation of evolutionary rates between interacting genes is evident even at the whole-interactome level in an organism (Fraser et al. 2002; Alvarez-Ponce and Fares 2012).

It should be noted, however, that not all of the correlation in evolutionary rates between interacting genes can be attributed to the coevolutionary process. Interacting proteins are usually involved in related biological functions and exhibit comparable expression patterns, which will lead to similarity in the selective pressure exerted on the proteins and give rise to the observed correlated evolutionary rate (Juan et al. 2008). In any case, an analysis of correlated evolution between interacting genes or other evolving units will provide insights into the organization and functionality of living organisms, whether the covariation derives from similar selective pressure or reciprocal evolutionary changes.

As mentioned above, correlated evolution and coevolution at the molecular level have been extensively studied, focusing mostly on functional genes in the genome. With the advance of chromosome conformation capture (3C) and its high-throughput derivate (i.e., Hi-C) technique (Lieberman-Aiden et al. 2009) during the last decade, another type of interaction, the physical contacts of genomic DNA in the nucleus, has been revealed at the whole-genome scale with an unprecedented resolution in many organisms. Similar to gene arrangement at the linear level, the 3D organization of the genome in the nucleus is far from random. In eukaryotes, chromosomes are arranged into discrete territories and other finer structures (e.g., A/B compartments, topologically associated domains, chromatin loops), and this architecture influences many biological activities such as gene transcription and splicing (Bonev and Cavalli 2016; Ruiz-Velasco and Zaugg 2017). Recent studies based on Hi-C data have shown that genes that are in direct contact in the 3D space of the nucleus tend to be coexpressed (Ben-Elazar et al. 2013; Homouz and Kudlicki 2013; Ay, Bunnik, et al. 2014; Babaei et al. 2015) and to present comparable biological functions based on different measurements of functional similarity (Homouz and Kudlicki 2013; Diament et al. 2014). In addition, fragments in physical contact in the 3D genome exhibit more correlated mutations (Perlaza-Jimenez and Walther 2018), suggesting functional interaction between 3D neighbors. Because expression level/breadth and functional constraints are two important predictors of the evolutionary rate of protein-coding genes (Pal et al. 2006; Zhang and Yang 2015), we expect that interacting genes in the 3D chromatin network will evolve in a coordinated fashion. However, to our knowledge, such a hypothesis has not been explicitly tested in any species. Furthermore, considering that interacting DNA fragments in the nucleus contain a substantial proportion of noncoding sequences, it is worth investigating whether these sequences subjected to lower level of selective pressure also evolve in a coordinated way.

Here, we explicitly tested the hypothesis that genes and other noncoding sequences located nearby in the 3D genome evolve in a coordinated manner using *Arabidopsis thaliana* as a case study. Specifically, we constructed chromatin interaction networks (CINs) with Hi-C datasets in *A. thaliana* and compared the similarity of genetic diversity and evolutionary rates between 3D neighboring fragments with the random expectation. We aimed to answer the following questions: 1). Do genetic fragments that interact at the 3D scale evolve in a coordinated manner in terms of genetic diversity and evolutionary rate? 2). If so, what evolutionary forces (e.g., mutation, natural selection) have driven the observed pattern?

In addition, it is known that chromatin organization and other genomic features (e.g., GC content, recombination rates, and replication timing) influence the regional mutation rate in both germline and somatic cells (Stamatoyannopoulos et al. 2009; Schuster-Bockler and Lehner 2012; Woo and Li 2012; Segurel et al. 2014; Makova and Hardison 2015) and, hence, affect local genetic diversity and evolutionary rates. With the availability of a plethora of (epi)genomic data in the model species Arabidopsis, we are able to comprehensively explore the relationships between local (epi)genomic features and regional evolutionary rates, along with their implications for the correlated evolution of 3D neighboring fragments.

## Results

### Neighboring fragments in the CIN show similar levels of nucleotide diversity and evolutionary rates

We split the Arabidopsis genome into 1,193 100 kb nonoverlapping windows (i.e., fragments), and constructed the CIN with both significant intra- and inter-chromosomal contacts using Hi-C data from Wang et al. (2015) and Liu et al. (2016). Hi-C is a high-throughput method that can reveal the spatial proximity of genomic DNA fragments (Lieberman-Aiden et al. 2009). The chromosome conformation captured by Hi-C can be represented as a network, in which a node denotes a genomic fragment while an edge stands for the presence of spatial proximity between two DNA fragments (see supplementary fig. S1, Supplementary Material online for a graphical representation of Hi-C and CIN). Because some of our analyses require a large number of sites from particular categories (e.g., 0-fold and 4-fold degenerate sites) within a single fragment to obtain reliable results, we retained only individual windows with more than 15 protein-coding genes where more than 20% of the sequences could be reliably aligned to *A. lyrata* (see Materials and Methods). The results obtained for the 50 kb resolution and other filtration criteria are generally similar (see Discussion and supplementary text, Supplementary Material online).

Both nucleotide diversity and evolutionary rates vary greatly across the genome at 100 kb scale (supplementary fig. S2, Supplementary Material online). Putatively selected sites present low levels of polymorphism and divergence while putatively neutral sites have higher diversity and evolve faster (supplementary fig. S3 and table S1, Supplementary Material online). Fragments with more connections in the CIN might be subjected to a higher level of selective constraint, as reflected by a strong negative correlation between the node degree (i.e., the number of interactions for a fragment) and mean diversity (fig. 1A; r = −0.46; P < 2.2 × 10^−16^) and a significant positive correlation between the node degree and the proportion of SNPs with a derived allele frequency of less than 0.5% [%(*DAF* < 0.005); fig. 1B; r = 0.43; P < 2.2 × 10^−16^]. Highly connected fragments also tend to evolve more slowly than those exhibiting fewer connections (fig. 1C; r = −0.53; P < 2.2 × 10^−16^). There is a significant positive correlation between the node degree and the number of genes within fragments (fig. 1D; r = 0.52; P < 2.2 × 10^−16^), which may explain the correlation between the degree centrality and selective constraints. However, the results of partial correlation analyses show that the number of genes within fragments cannot fully explain the observed pattern. After accounting for the variation of gene numbers, the correlation coefficients between the node degree and diversity, %(*DAF* < 0.005), and divergence are −0.36, 0.36, and −0.41, respectively (P = 5.0 × 10^−30^, 5.6 × 10^−31^, and 6.4 × 10^−40^, respectively). Overall, our results recapitulate the findings made in protein-protein interaction networks (Fraser et al. 2002; Alvarez-Ponce and Fares 2012), for which proteins with higher degree of connectivity evolve more slowly.

**Figure 1.**
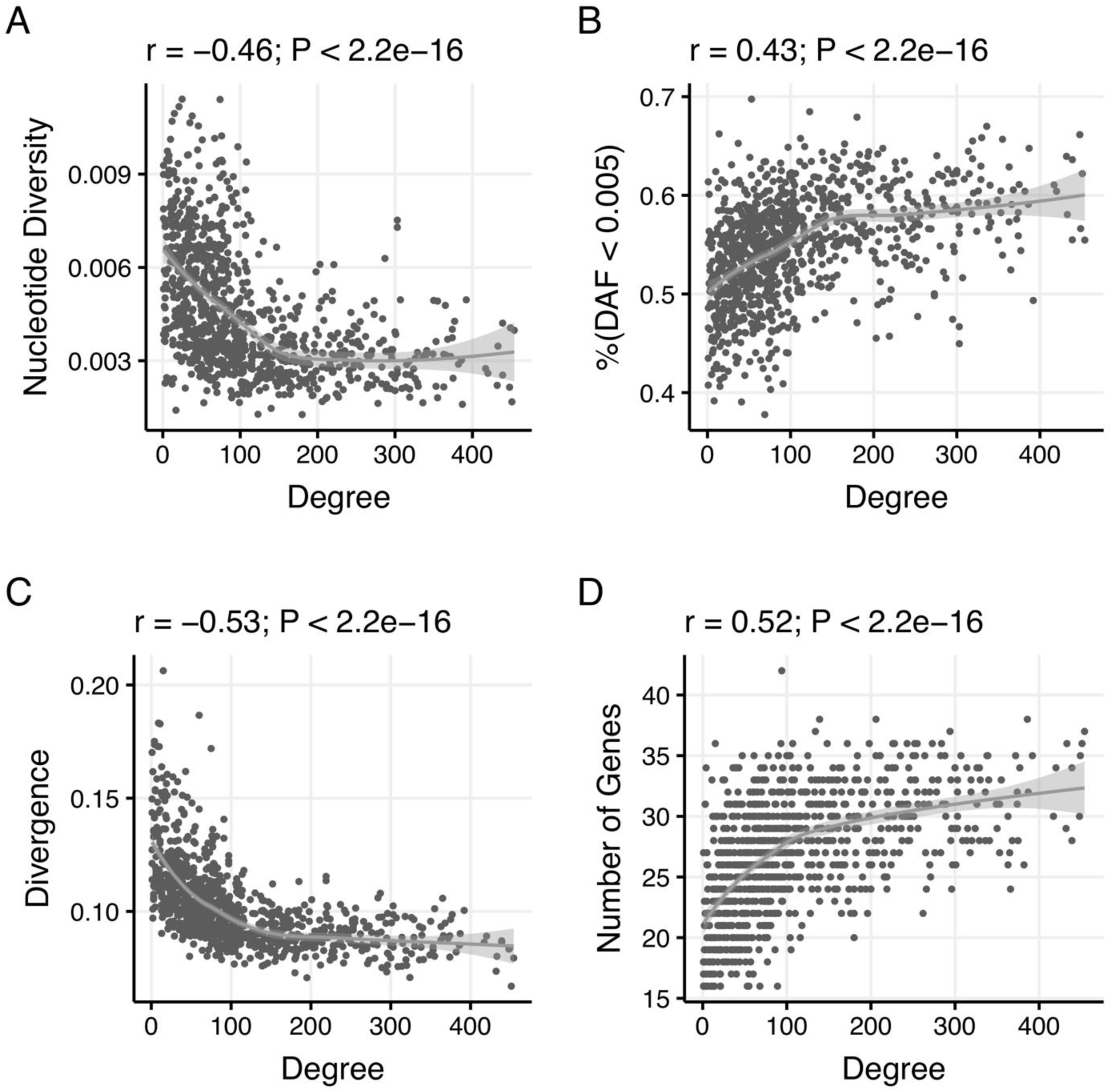
Fragments with more connections in the CIN are subjected to higher levels of selective constraints and evolve more slowly. The four panels show the correlation between the degree in the CIN and nucleotide diversity (A), %(*DAF* < 0.005) (B), evolutionary rates (C), and the number of genes (D). The gray line and shaded area are the loess fitting line and its 95% confidence interval, respectively.

To test whether large DNA fragments that are in contact in the 3D space evolve in a coordinated fashion, we compared the observed average differences in nucleotide diversity and evolutionary rates between immediate neighbors in the CIN to those in different null models. In the first model, we rewired the graph while the degree distribution was preserved. For both polymorphism and divergence, the observed difference between fragments in contact is significantly smaller than that of the random networks (P < 0.001 in both cases; supplementary fig. S4, Supplementary Material online), suggesting that neighboring fragments in the 3D genome indeed evolve in a coordinated way. Since DNA fragments that are in proximity at the linear scale tend to be neighbors in the 3D space as well and physically linked genes tend to be coexpressed (Boutanaev et al. 2002; Williams and Bowles 2004) and evolve at similar rates (Williams and Hurst 2000), the detected similarity may be a product of one-dimensional organization of the genome. To test whether the 3D arrangement of DNA in the nucleus contributes to the observed correlated evolution, we performed a more conservative simulation, the cyclic chromosome shift (CCS) model (Diament et al. 2014), in which the linear adjacency among fragments is preserved along their “circular” chromosomes, while their 3D interacting relationship is perturbed (supplementary fig. S1B, Supplementary Material online). The differences in genetic diversity and evolutionary rates in the real data are still significantly smaller than those in the simulated networks (P < 0.001 in both cases; fig. 2A and B). This conclusion also holds under the cyclic genome shift (CGS) model, which is similar to the CCS model but allows fragments to rotate and move between different chromosomes (supplementary figs. S1B and S5, Supplementary Material online).

**Figure 2.**
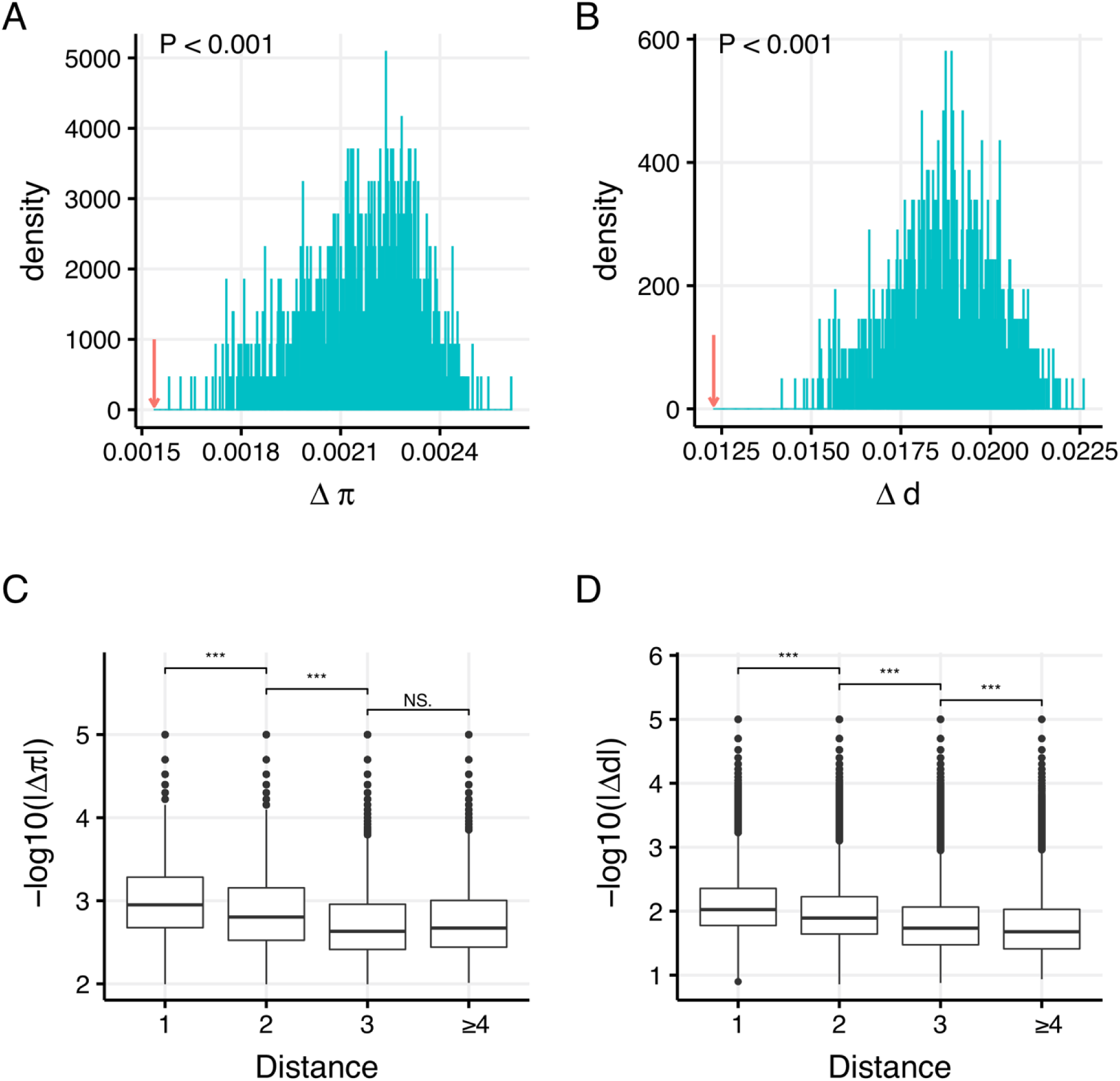
Large DNA fragments that are in direct contact in the CIN display a similar genetic diversity and evolutionary rate, and this similarity cannot be explained by their linear adjacency. The upper two panels show the distribution of the mean difference in genetic diversity (A) and evolutionary rates (B) between adjacent fragments in 1,000 simulated networks using the CCS null model. The blue lines show the simulated distribution of a parameter, while the red arrow represents the observed mean difference of a parameter. The P-value calculated as the number of mean differences in simulated networks less than the observed difference is shown at the top left. The lower two panels show that the similarity in genetic diversity (C) and evolutionary rates (D) between fragment pairs decreases when the 3D distance between them increases. *** P < 0.001, NS: nonsignificant.

The CCS model perturbs the CIN while keeping the following two properties intact: 1) the spatial organization of chromosomes (i.e., network topology); 2) the linear proximity between fragments along their respective chromosomes. However, for the degree-preserving rewiring of the network, only the first property is maintained, while for the CGS model, fragments are allowed to move to other chromosomes, therefore the second property is not as well preserved as in the CCS model. If the observed difference between connected nodes in the true network is significantly smaller than that expected by the CCS model, then the 3D chromosomal conformation other than the linear arrangement of chromosomes would be a major factor explaining the observed correlated evolution. The CCS model is the most logic null model for testing the effects of spatial proximity on the correlated evolution between connected fragments in the CIN, therefore, we only present the results of the CCS simulations in the following sections.

Furthermore, the similarities in nucleotide diversity and evolutionary rates between fragment pairs both decrease with an increase in topological distance in the CIN (fig. 2C and D). These results indicate that the large fragments containing genes and other DNA elements are arranged in a nonrandom manner in Arabidopsis and that 3D chromatin organization contributes to the correlated evolution of genetic elements that are close to each other in higher dimensional space.

### Evolutionary forces that contribute to the correlated evolution between 3D chromatin neighbors: mutation rate

The levels of genetic diversity and evolutionary rate of a region are shaped by a combination of evolutionary forces at different timescales (Hartl and Clark 2007). To understand why neighboring fragments in the 3D space show similar levels of diversity and divergence, we investigated the roles of different evolutionary forces that shape the polymorphism and evolutionary rates of large DNA fragments. We first assessed the contribution of the local mutation rate, which varies in the eukaryotic genome at different scales (Hodgkinson and Eyre-Walker 2011; Terekhanova et al. 2017; Smith et al. 2018). We first used the average substitution rate at putatively neutral sites [4-fold degenerate sites and putatively neutral noncoding sites (NNSs; see definition in Materials and Methods)] between *A. thaliana* and *A. lyrata* as a surrogate of the mutation rate and used the CCS simulation to assess the significance of the similarity between 3D neighboring fragments. The result shows that the observed difference is significantly smaller than that in the simulated networks (P < 0.001; fig. 3A), suggesting that the local mutation rate covaries between 3D adjacent fragments and plays a role in the correlated evolution between 3D neighbors. The conclusion is unaffected when we assess the similarity in mutation rate between 3D adjacent fragments using either the divergence at 4-fold degenerate sites or NNSs alone as a proxy of the local mutation rate (P < 0.001 for both cases; supplementary fig. S6A and B, Supplementary Material online).

**Figure 3.**
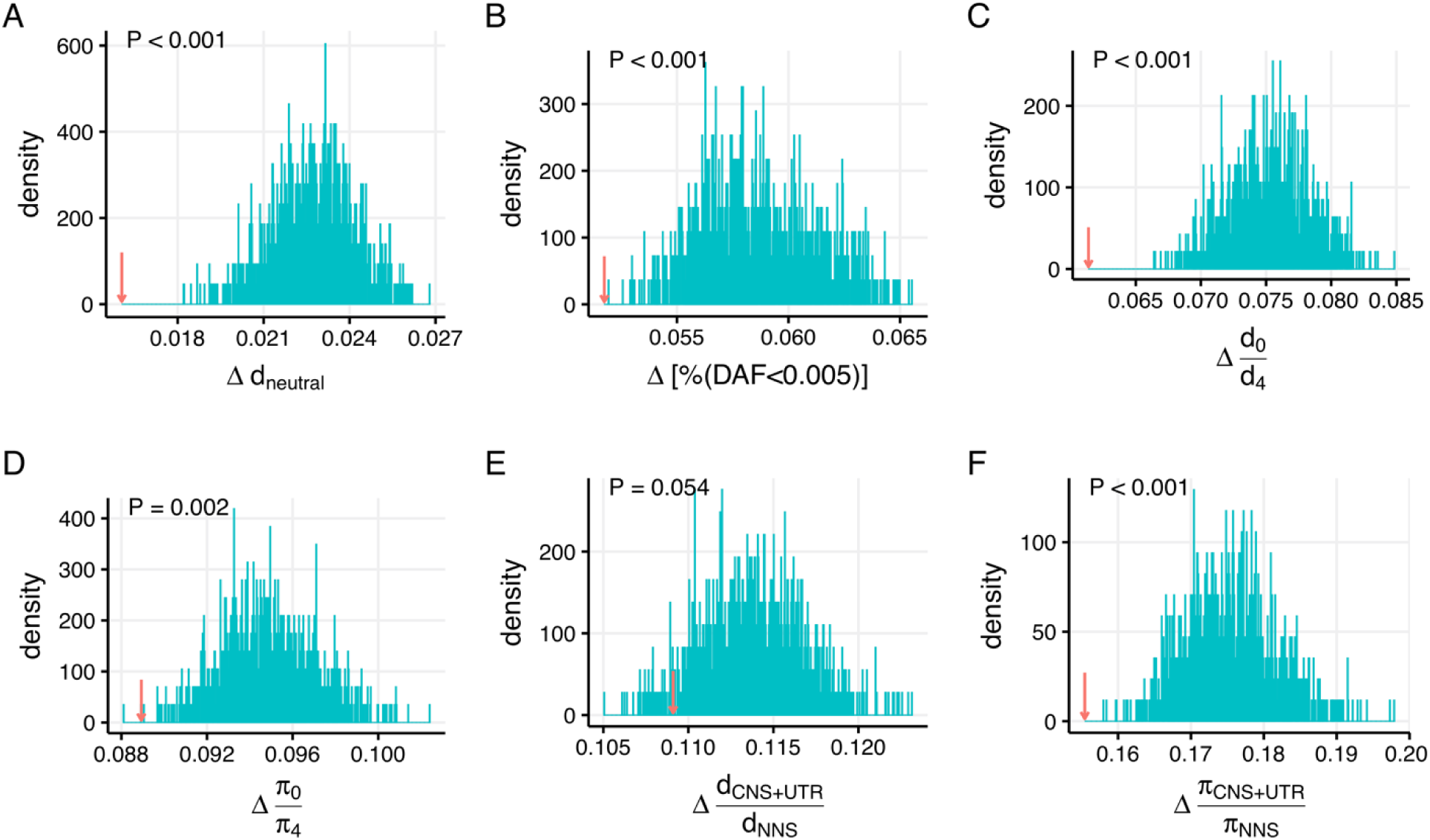
Neighboring fragments in the CIN exhibit similar regional mutation rates and evolutionary constraints. The observed average difference and the distribution of the average difference between adjacent nodes in simulated networks are shown for divergence at putatively neutral sites (*d_neutral_*; A), %(*DAF* < 0.005) (B), *d*_0_/*d*_4_ (C), *π*_0_/*π*_4_ (D), *d_CNS+UTR_*/*d_NNS_* (E), and *d_CNS+UTR_*/*d_NNS_* (F).

Although the putatively neutral sites as defined above exhibited high levels of genetic diversity and evolutionary rates (supplementary fig. S3, Supplementary Material online), they may be under some level of selection and their evolution is not affected by mutation alone. Thus, we further used 1,694 *de novo* mutations identified in *A. thaliana* mutation accumulation lines (Weng et al., 2019) to assess the similarity in mutation rate between spatially adjacent fragments at 100 kb scale. The data of *de novo* mutations are sparse (supplementary fig. S6C, Supplementary Material online), but the average difference in the number of mutations between 3D neighboring fragments is significantly smaller than that in the simulated networks (P = 0.001; supplementary fig. S6D, Supplementary Material online).

### Evolutionary forces that contribute to the correlated evolution between 3D chromatin neighbors: purifying selection

To assess the contribution of selective constraints to the correlated evolution of neighboring fragments, we estimated various population genetic parameters to measure the strength and efficacy of purifying selection acting on a whole fragment or functional elements in the fragment. First, the level of excess rare variants can be interpreted as the relative strength of purifying selection across different classes of genomic regions (Altshuler et al. 2012; Khurana et al. 2013). Therefore, we calculated %(*DAF* < 0.005) in each fragment using polymorphism data from the 1001 Genomes Project (Alonso-Blanco et al. 2016). The result shows that the simulated networks present a higher degree of difference between connected nodes than that in the true network (P < 0.001; fig. 3B). Next, we used the ratios of divergence and diversity of 0-fold versus 4-fold degenerate sites (denoted as *d*_0_/*d*_4_ and *π*_0_/*π*_4_, respectively) to quantify the strength or the efficacy of the negative selection acting on protein-coding genes in each fragment. For both estimates, the average difference between connected nodes in the real network is significantly smaller than that in the simulated networks (P < 0.001 and P = 0.002 for *d*_0_/*d*_4_ and *π*_0_/*π*_4_, respectively; fig. 3C and D). Our results suggest that adjacent fragments and the genes they contain in the CIN are subjected to more similar evolutionary constraints than random pairs, which agrees with previous reports that neighboring genes in the 3D space present more similar functions and expression patterns than random gene pairs. We also compared the similarity of the strength and efficacy of the negative selection of potentially functional noncoding sequences (CNS+UTR: see definition in Materials and Methods) between chromatin neighbors to that of the null model. In these cases, we used putatively neutral noncoding sites (i.e., NNSs) as neutral comparators. The results show that the difference in *d_CNS+UTR_*/*d_NNS_* between connected fragments in the real network is marginally smaller than that in the simulated networks (P = 0.054; fig. 3E), while the disparity between the true data and the null model is significant for *π_CNS+UTR_*/*π_NNS_* (P = 0.004; fig. 3F).

We further quantified the strength of purifying selection on 0-fold nonsynonymous sites for each fragment using the DFE-alpha software (Keightley and Eyre-Walker, 2007). The DFE-alpha method simultaneously estimates the distribution of fitness effects (DFE) of new deleterious mutations and the rate of adaptive evolution, in which it accounts for the effects of demographic changes and slightly deleterious mutations on the estimation. To this end, we utilized biallelic SNPs from two separate populations, one containing 22 relict individuals from the Iberian Peninsula (denoted as the relict population) and the other comprising 50 accessions from Spain. For both populations, the difference of the proportion of effectively neutral mutations (0 < *N_e_s* < 1) between 3D neighboring fragments is significantly smaller than that expected by the CCS model. However, for each category of mutations with different strength of deleterious effects, there are no statistically significant differences between the true network and simulated networks (supplementary figs. S7 and S8, Supplementary Material online). Our results suggest that the proportion of neutral mutations is more similar between connected fragments in the CIN than the random expectation, but the full DFE may differ.

### Evolutionary forces that contribute to the correlated evolution between 3D chromatin neighbors: positive selection

Although several studies have shown that the *A. thaliana* genome exhibits little evidence of genome-wide adaptive selection (Bustamante et al. 2002; Foxe et al. 2008; Wright and Andolfatto 2008; Gossmann et al. 2010; Slotte et al. 2011), substantial heterogeneity in the rate of positive selection across different categories of genes is also evident (Slotte et al. 2011). Previous studies have demonstrated that protein function is partly responsible for this heterogeneity in Arabidopsis and other organisms (Obbard et al. 2009; Slotte et al. 2011; Enard et al. 2016). Notably, genes associated with immune defence mechanisms are subjected to more adaptive evolution. It is also known that spatially adjacent genes have similar functions (Homouz and Kudlicki 2013; Diament et al. 2014). Thus, do clusters of genes that are close to each other in the 3D space possess similar rates of adaptive evolution? To answer this question, we calculated the proportion of adaptively driven substitutions (*α*) and the rate of adaptive substitution relative to the rate of neutral evolution (*ω_a_*) for 0-fold sites in each 100 kb fragment using the software DFE-alpha (Eyre-Walker and Keightley 2009). We quantified these two measurements using divergence information from the *A. thaliana*-*A. lyrata* alignment and SNPs from either the relict or the Spanish population. As for the relict population, the average *α* is 20.27% (fig. 4A), which is substantially higher than the proportion previously reported, whereas the average *α* estimated using SNPs in the Spanish population is 6.24% (fig. 4D), a number that is very close to the results of previous studies (Gossmann et al. 2010; Slotte et al. 2011). The discrepancy in adaptive rates estimated using polymorphisms from the two populations could exist because the relict and Spanish populations have different historical effective population sizes. In the Spanish population, natural selection is less effective in removing slightly deleterious mutations and fixing slightly advantageous mutations because of the stronger effect of genetic drift in small populations, resulting in a reduction in *α*. When we compare the observed difference in *α* between neighboring nodes to that in the null model, no significant difference is found between the actual and simulated values for results derived from both the relict and Spanish populations (P = 0.923 and 0.639, respectively; fig. 4B and E). On the other hand, the difference in *ω_a_* between neighboring fragments is significantly smaller than that in the simulated networks for estimates derived from the relict and Spanish populations (P = 0.010 and 0.002, respectively; fig. 4C and F).

**Figure 4.**
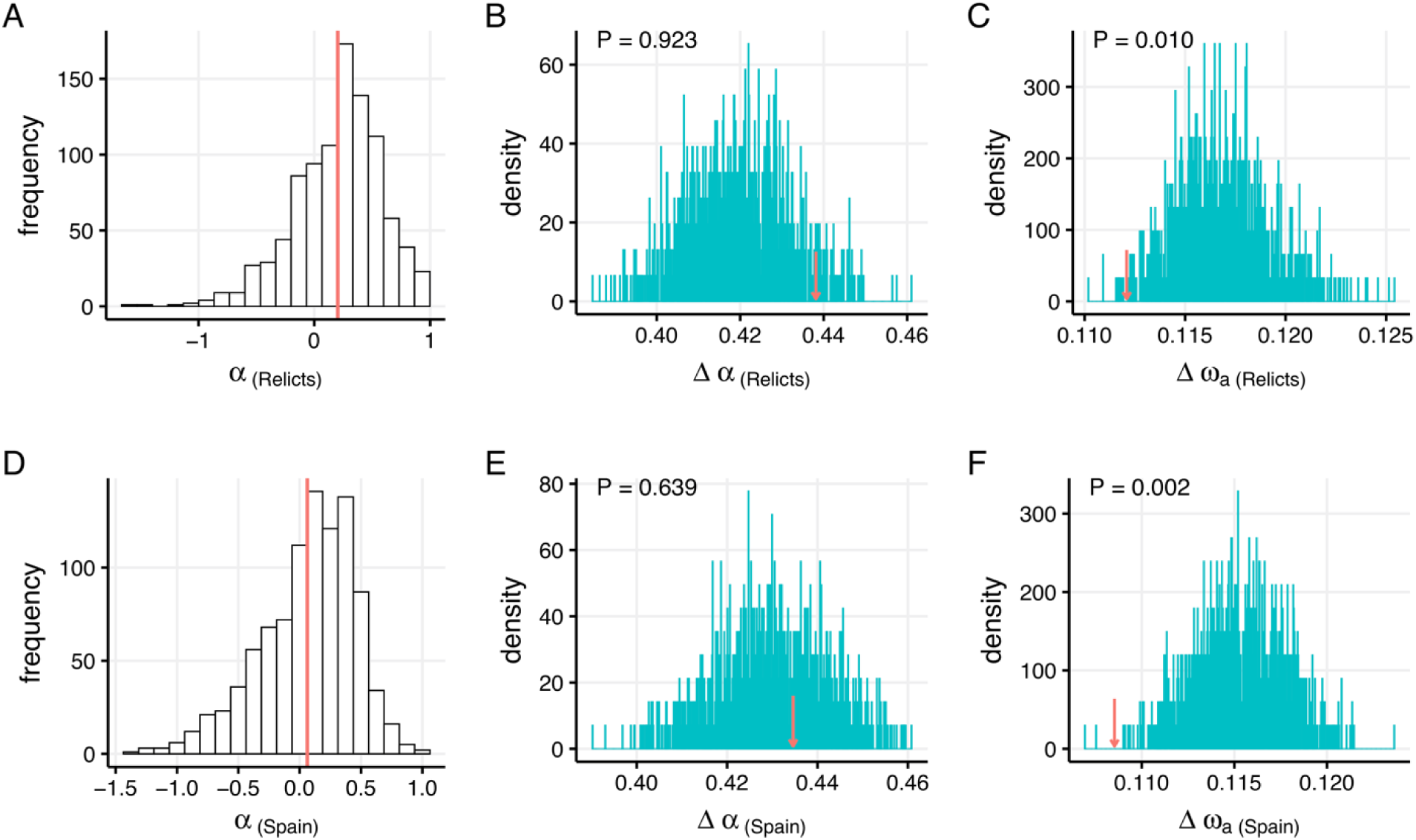
The rate of adaptive evolution and similarity of positive selection between 3D neighboring fragments. The upper three panels show the distribution of *α* (A) in 100 kb fragments and the results of CCS simulations for *α* (B) and *ω_a_* (C) using polymorphism data from the relict population. The lower three panels show the distribution of *α* (A) in 100 kb fragments and the results of CCS simulations for *α* (B) and *ω_a_* (C) using polymorphism data from the Spanish population.

The rate of nonadaptive substitution relative to the rate of neutral evolution (*ω_na_* = *ω* − *ω_a_*, where *ω* = *d*_0_/*d*_4_) covaries between 3D neighboring fragments as well (supplementary fig. S9, Supplementary Material online). *ω_na_* is proportional to the average fixation probability of a selected mutation, 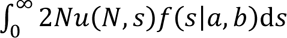, where *N* is the population size, *u*(*N*, *s*) is the fixation probability of a new mutation of selective effect *N_e_s*, and *f*(*s*|*a*, *b*) is the DFE of deleterious mutations estimated from polymorphism data (Eyre-Walker and Keightley 2009). The covariation in *ω_na_* indicates that, in the absence of advantageous mutations, the predicted divergence at selected sites is more similar between connected fragments in the CIN than that expected by the null model.

### (Epi)genomic determinants of nucleotide diversity and divergence across the genome

Mutation, natural selection, and random genetic drift together shape the variation of genetic diversity within populations and divergence between species. Factors that affect these evolutionary parameters thus may contribute to the observed correlated evolution between 3D neighbors. Such factors include recombination rates, GC content, gene density, replication timing, and chromatin organization (Makova and Hardison 2015). To elucidate the influence of (epi)genomic features on the regional variation in genetic diversity and divergence and relate them to the large-scale correlated evolution in the 3D genome, we compiled a comprehensive set of genetic and epigenetic features that have been mapped genome wide in Arabidopsis (supplementary table S2, Supplementary Material online). We first clustered these features using their pairwise correlation coefficients as a distance metric and found that they are largely clustered into two groups. The first group includes features associated with open chromatin, active transcription, and early replication timing, and the second group contains features associated with heterochromatin, inactive transcription, and late replication timing (fig. 5A). Remarkably, most epigenomic features associated with open chromatin present a strong anti-correlation with polymorphism and evolutionary rates (fig. 5B), with H3K36me3 shows the highest negative correlation with the substitution rate (r = −0.74, P < 2.2 × 10^−16^), while the density of DNase I hypersensitive sites (DHSs) in flowers exhibits the highest negative correlation with nucleotide diversity (r = −0.60, P < 2.2 × 10^−16^). On the other hand, histone variants and epigenetic marks that define heterochromatin and inactive transcription show an opposite trend (fig. 5B), among which the histone variant H2A.W presents the highest positive correlation with the divergence between *A. thaliana* and *A. lyrata* (r = 0.69, P < 2.2 × 10^−16^), while H3.1 presents the highest positive correlation with nucleotide diversity (r = 0.64, P < 2.2 × 10^−16^).

**Figure 5.**
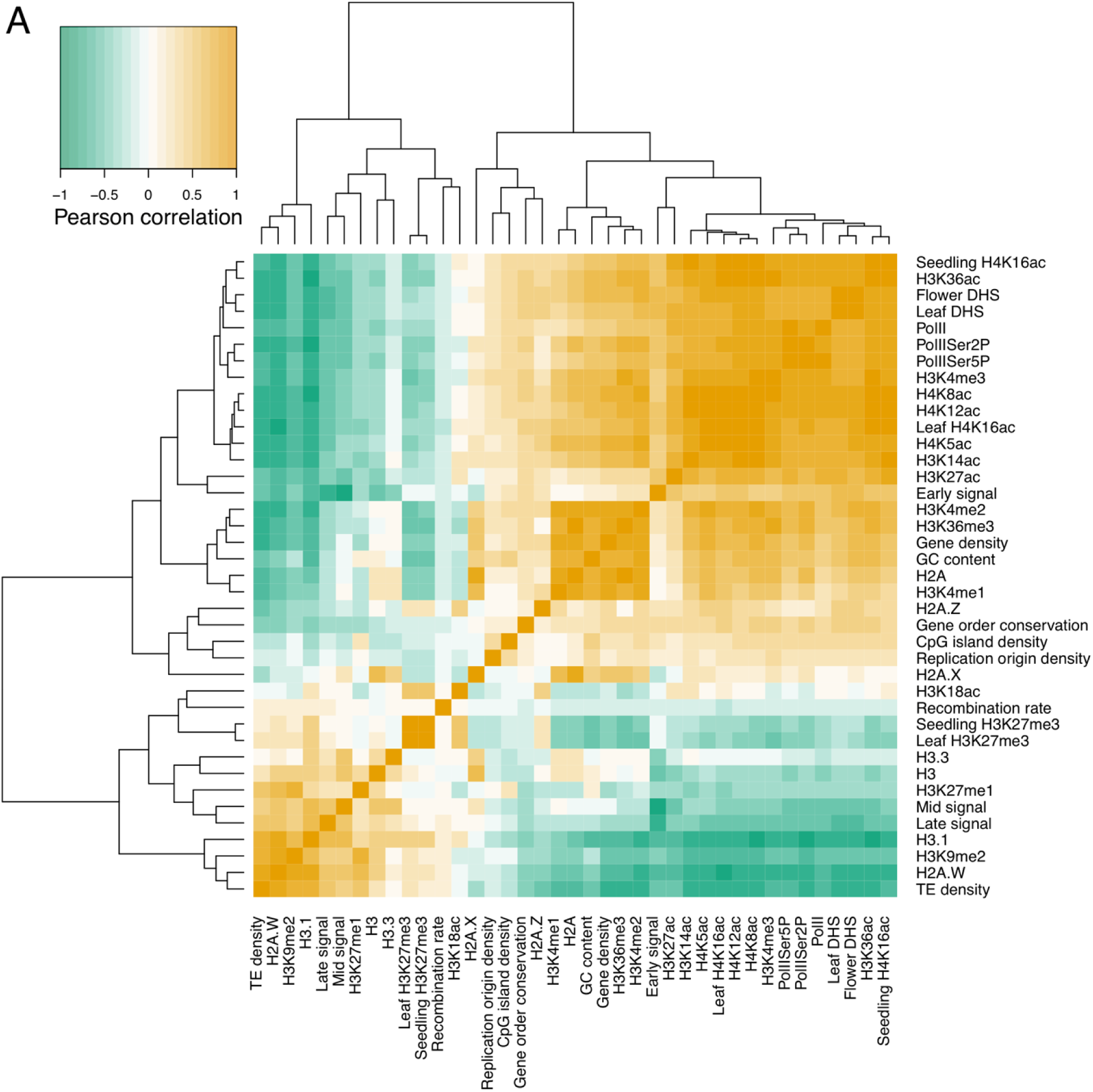
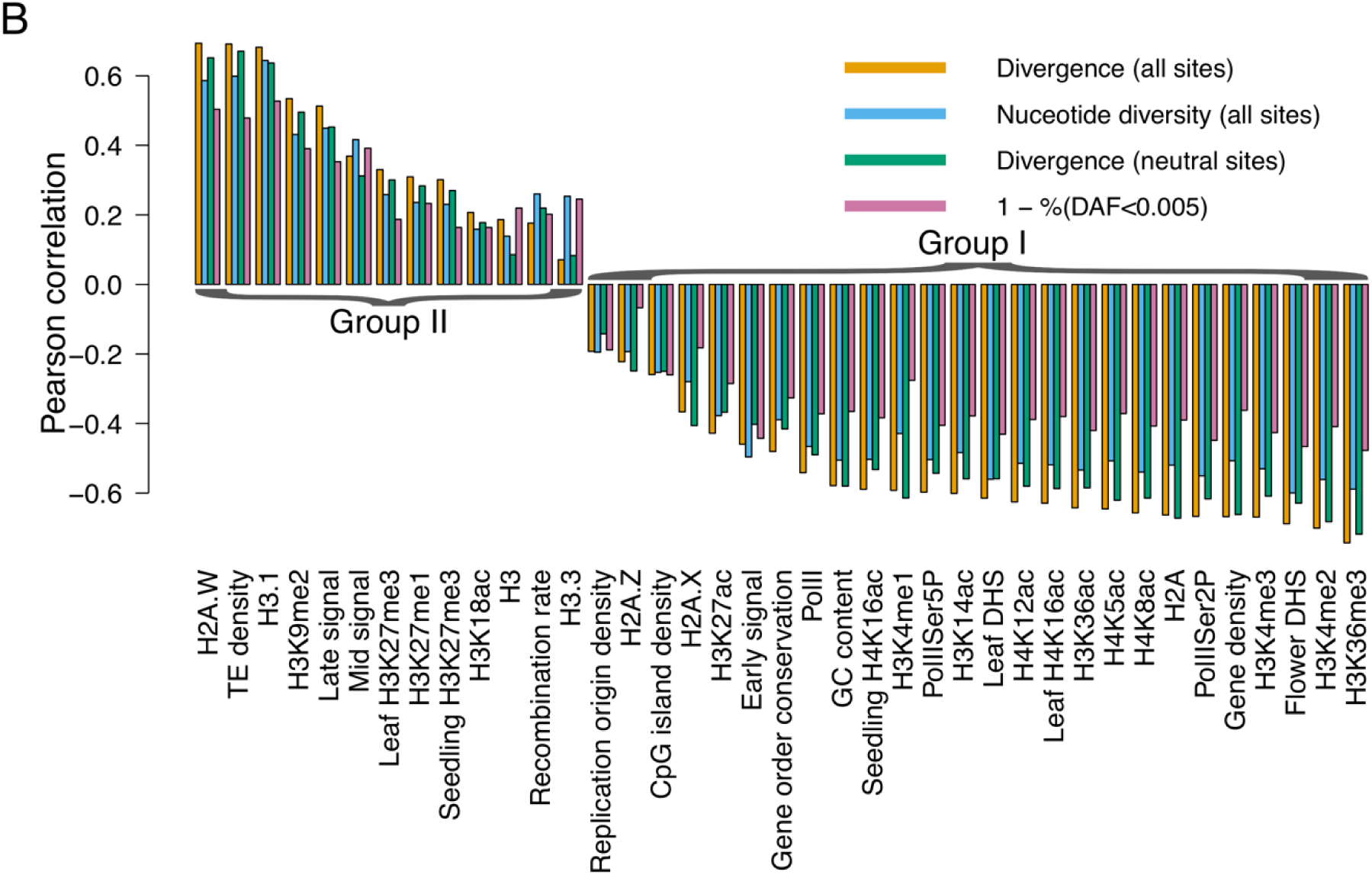
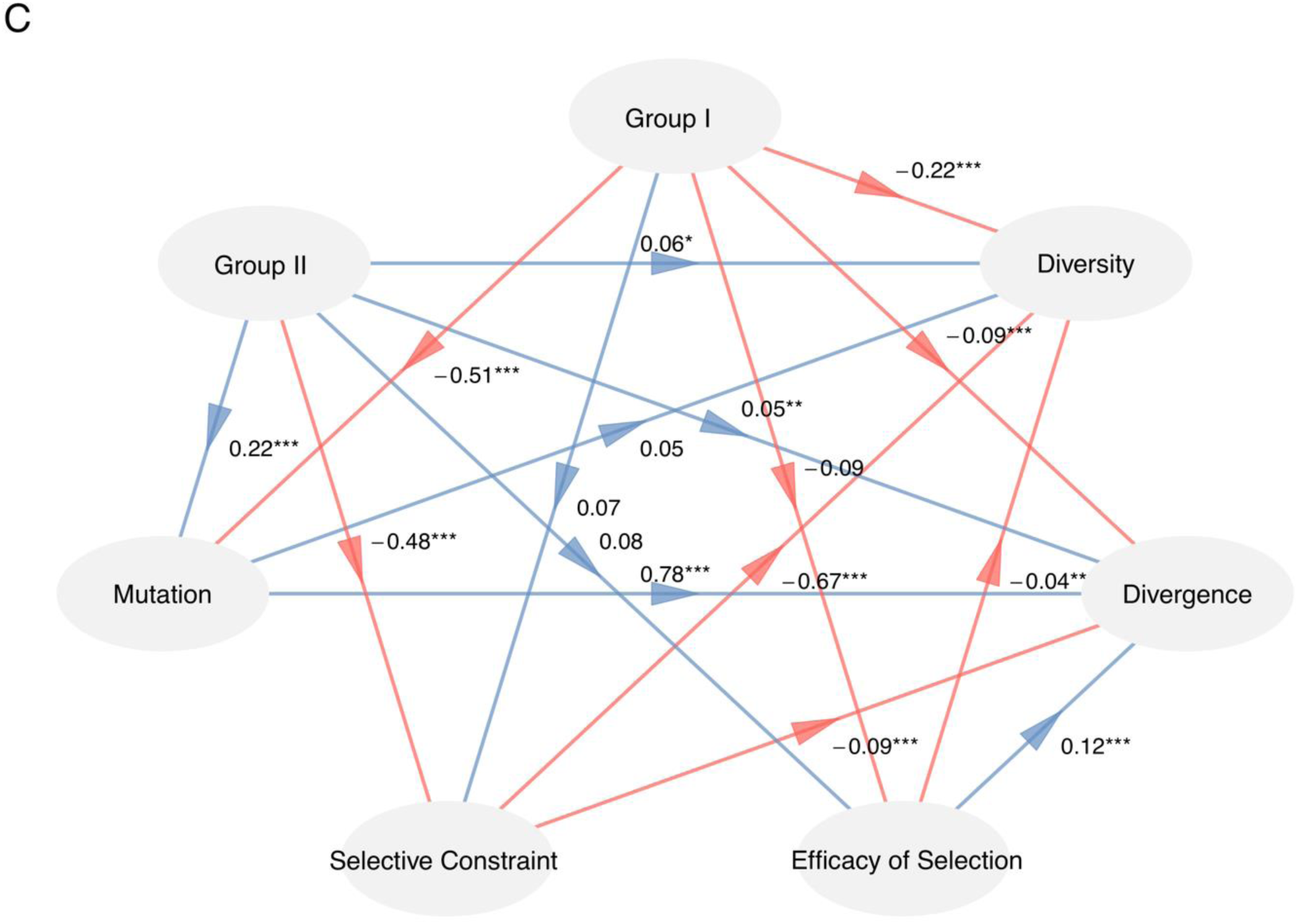
A set of diverse (epi)genomic features affect local nucleotide diversity and evolutionary rates along the genome. (A) Correlation heatmap of 39 (epi)genomic features. (B) Pearson’s correlation coefficients of divergence, genetic diversity, divergence at putatively neutral sites and 1 − %(*DAF* < 0.005) with diverse (epi)genomic features in 100 kb nonoverlapping windows. (C) The result of PLS-PM showing the causal relationship between the two groups of features and evolutionary parameters. Divergence at putatively neutral sites, %(*DAF* < 0.005), and *π*_0_/*π*_4_ are used to represent mutations, selective constraint, and the efficacy of selection, respectively. Arrows represent the predefined causal directions between variables; red indicates a negative impact and blue a positive one. The path coefficients and their significance are labeled beside the arrows. *** P < 0.001, ** P < 0.01, and * P < 0.05.

Chromatin organization has a great impact on regional mutation-rate variation in the genome at various scales (Schuster-Bockler and Lehner 2012; Makova and Hardison 2015; Terekhanova et al. 2017). Thus, the observed pattern of correlation between epigenetic marks and evolutionary parameters may be mediated by the influence of chromatin architecture on the local mutation rate. Indeed, we found that epigenetic features show the same trend of correlations with substitution rates at putatively neutral sites between *A. thaliana* and *A. lyrata* (which serves as a proxy of the mutation rate) as observed between epigenetic marks and nucleotide diversity and divergence, but to a lesser extent (fig. 5B). Using %(*DAF* < 0.005) as a measure of selective constraint, we found that regions with higher levels of heterochromatin signals are subjected to lower selective constraints, and an opposite trend was observed for regions marked by higher levels of open chromatin signals (fig. 5B). Taken together, these results support the notion that chromatin organization has a great influence on the evolution of regional DNA fragments, which is consistent with previous studies in humans (Schuster-Bockler and Lehner 2012; Makova and Hardison 2015).

Next, we sought to determine which genetic or epigenetic features have the most significant impact on regional nucleotide diversity and evolutionary rates and how much of the local variation in diversity and divergence can be explained by the combination of features. We found that 57%, 44%, 70%, and 64% of the variance in nucleotide diversity, selective constraint, evolutionary rates, and mutation rates can be explained by combining all features (fig. 6). For polymorphisms within species, the histone variant H3.1 alone can account for 41% of the variance in regional genetic diversity and 28% of the variance in selective constraints (fig. 6A and B). The expression of H3.1 is highly coupled with the S phase of cell cycle, and the deposition of H3.1 is associated with chromatin restoration after DNA replication and possible DNA repair (Tagami et al. 2004). H3.1 is enriched in silent regions of the genome, including areas that are marked by repressive chromatin modifications H3K9me2 and H3K27me3, and DNA methylation (Stroud et al. 2012). A considerable proportion of the variance in the divergence between species along the genome can be explained by the histone modification H3K36me3, where the explained proportion reaches 55% for overall divergence and 52% for divergence at putatively neutral sites (fig. 6C and D). In animals, H3K36me3 is enriched at the 3’ end of actively transcribed genes, and is associated with efficient transcription elongation (Xiao et al. 2016). In Arabidopsis, H3K36me3 is also coupled with active gene expression and transcription elongation (Sequeira-Mendes et al. 2014), but it’s signal peaks at the 5’-half of actively transcribed genes (Mahrez et al. 2016; Xiao et al. 2016). In contrast, the variance of the efficacy of natural selection is poorly explained by these features. Only 18% and 7% of the variance in *π*_0_/*π*_4_ and *ω_a_* can be explained by the combination of features, with no single feature explaining more than 5% of the variance in both cases (fig. 6E and F).

**Figure 6.**
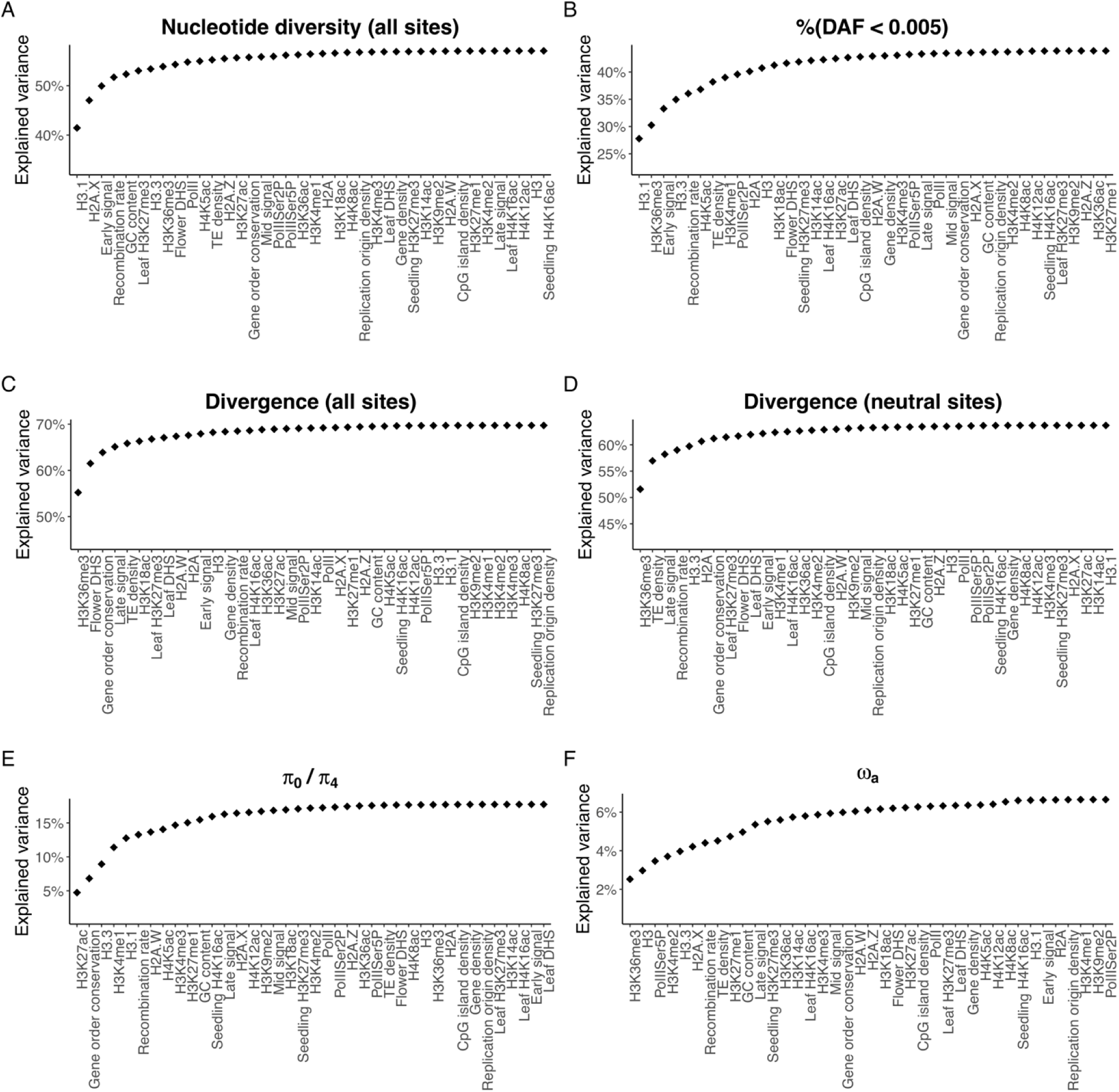
The combination of multiple (epi)genomic features can explain a great proportion of variation of genetic diversity and evolutionary rates across the Arabidopsis genome. The cumulative *R*^2^ of linear models (adding the feature on the *x* axis as a predictor at each step) for nucleotide diversity (A), %(*DAF* < 0.005) (B), divergence at all sites (C), divergence at neutral sites (D), *π*_0_/*π*_4_ (E), and *ω_a_* (F) are shown.

To discern the evolutionary force through which the investigated (epi)genomic characteristics affect the observed correlated evolution, we carried out a partial least squares path modeling (PLS-PM) analysis. PLS-PM is a statistical method for analyzing complex multivariate relationships among variables, in which several observed variables are summarized into many fewer latent variables, and the path coefficients and their significances between latent variables are calculated using a predefined causal network (Sanchez 2013). We used two latent variables to represent the two groups of features: group I contained features that are negatively correlated with genetic diversity and evolutionary rate, and group II included features that show positive correlations. The result of PLS-PM analysis shows that both groups have significant impacts on local mutation rates, but in opposite directions (path coefficient = −0.51 and 0.22 for group I and group II, respectively; fig. 5C). The mutation rate in turn significantly affects regional evolutionary rates (path coefficient = 0.78) but has limited influence on regional genetic diversity (path coefficient = 0.05). Under the neutral theory, evolutionary changes between species and polymorphisms within species are mainly due to neutral or nearly neutral mutations (Kimura 1983). However, it should be noted that nucleotide diversity at neutral sites is also affected by linked selection (background selection and genetic hitchhiking), whose variation within genome is shaped by the strength of selection, the rate of recombination, and the density of selected sites. This result suggests that linked selection may play a more important role in determining regional nucleotide variation than local mutation rate. *A. thaliana* is a selfing species with a low effective recombination rate, therefore, linked selection may be strong even in 100 kb fragments. On the other hand, substitution rate is less affected by linked selection. Only group II has a significant influence on the selective constraint, which in turn significantly affects diversity to a great extent and has a limited yet significant influence on divergence. Neither group I nor group II has a significant impact on the efficacy of selection (measured as *π*_0_/*π*_4_). In addition, the two groups have direct influences on diversity and divergence, indicating that some of the variance in polymorphisms and evolutionary rates cannot be explained by our measures of mutation, selective constraint, and the efficacy of selection. The loadings and cross-loadings of the observed indicators with the two latent variables are shown in supplementary fig. S10, Supplementary Material online.

### (Epi)genomic features that have a greater influence on the genetic diversity and evolutionary rate of individual fragments also exhibit higher levels of assortativity in the CIN

(Epi)genomic features that have a significant influence on the variation in regional nucleotide diversity and evolutionary rates may not necessarily contribute to the correlated evolution of neighboring nodes in the CIN. To further explore which features make greater contributions to the correlated evolution between 3D neighboring fragments, we calculated the assortativity of each feature in the network. Assortativity measures the extent to which interacting nodes in a network present similar values for a feature, which is an approach that was first introduced in the social sciences (Newman 2002) and has been applied in CINs recently (Pancaldi et al. 2016). A higher positive assortativity of a feature means a higher propensity for neighboring nodes to present similar values. If a feature exhibits a high level of assortativity in the network and has a significant influence on the regional variation of evolutionary rates, then this feature would make an important contribution to the correlated evolution between neighboring fragments in the 3D genome. Among the 39 (epi)genomic features, the occupancy of Pol II (assortativity = 0.211) and its variants (PolIISer2P, 0.269; PolIISer5P, 0.236), H3.1 (0.362), H2A.W (0.257) and H3K36ac (0.253) show relatively high levels of assortativity, whereas the assortativity is very low for the density of replication origins (0.003) and recombination rates (0.010).

We further correlated the levels of the assortativity of features with their Pearson correlations with evolutionary parameters. Interestingly, those features exerting a greater influence on regional genetic diversity and evolutionary rates also present higher levels of assortativity (fig. 7A and B). For those features in group I, the correlations between the levels of assortativity and their correlations with diversity and divergence are 0.84 (P = 2.9 × 10^−4^) and 0.68 (P = 1.0 × 10^−2^), respectively, while the correlations for the features in group II are −0.86 (P = 1.3 × 10^−8^) and −0.79 (P = 1.7 × 10^−6^), respectively. Features with higher levels of assortativity also have a greater influence on local mutation rates and selective constraints (fig. 7C and D). The correlation between the levels of the assortativity of features and their effects on the evolution of local fragments is strong and significant, which may have profound implications for the organization of the eukaryotic genome. If a genetic or epigenetic feature that has a significant impact on the evolution of regional fragments but presents a very low level of assortativity, then the rapid evolution of one fragment may impose pressure on the evolution of its interacting counterpart, leading to functional incompatibility between the products of spatially contacting DNA fragments.

**Figure 7.**
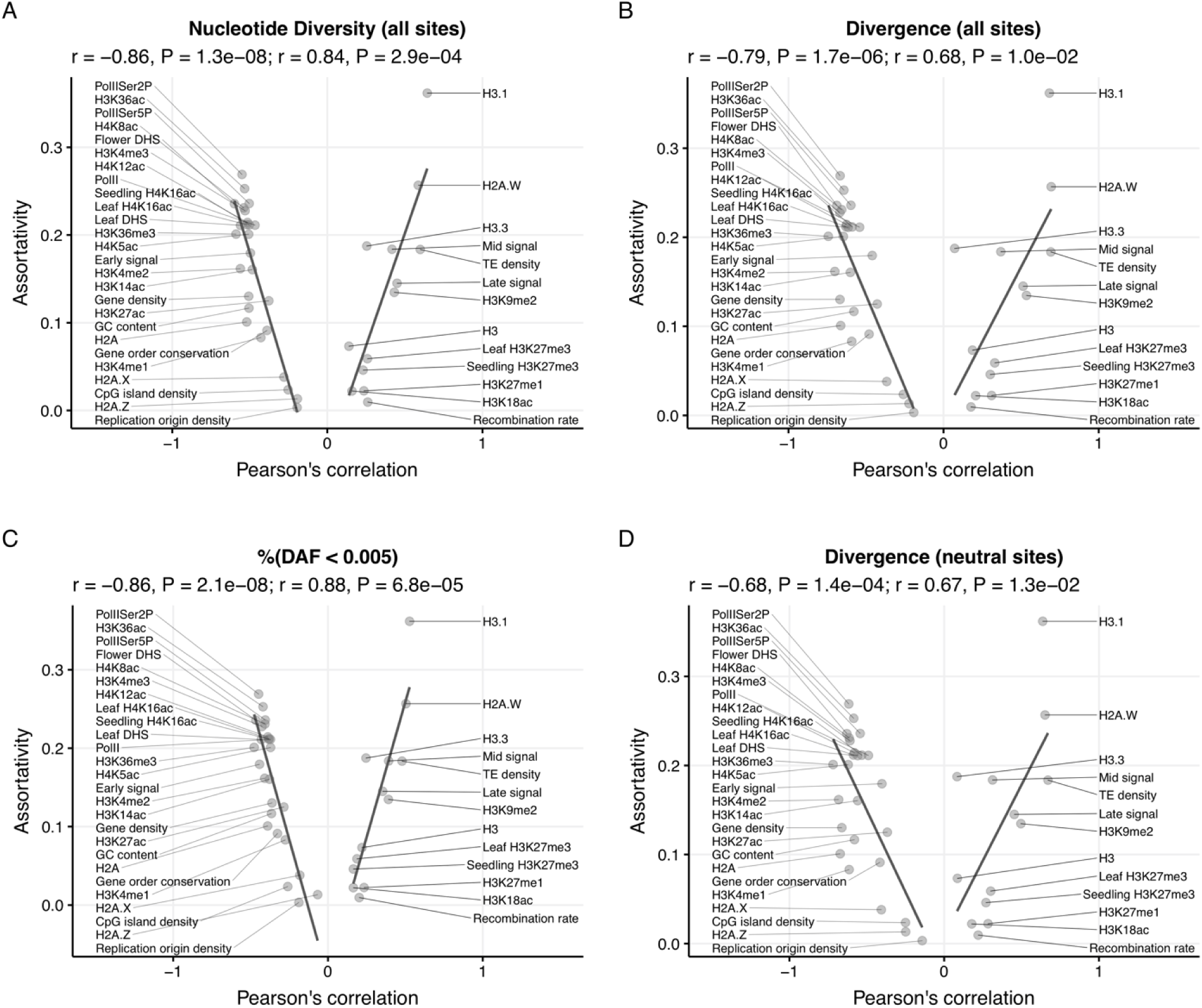
(Epi)genomic features that have a higher impact on the evolution of regional sequences exhibit higher levels of assortativity in the CIN. The *x* axis gives the Pearson’s correlation coefficient of each genomic feature with genetic diversity (A), evolutionary rate (B), selective constraint (C), and mutation rate (D). The *y* axis gives the assortativity of each feature in the CIN, which is the same in all panels. The solid lines represent the linear best fits between assortativity and the Pearson’s correlation coefficients, and are obtained for r > 0 and r < 0 separately.

## Discussion

Correlated evolution between biological units at various scales, whether derived from shared selective constraints or reciprocal evolutionary changes, has provoked the interests of biologists for a long time (Carmona et al. 2015). Here, we investigated the correlated evolution of large DNA fragments based on the three-dimensional chromosomal conformation in *A. thaliana* and examined the evolutionary forces and (epi)genomic features that potentially affect the large-scale correlated evolution in the 3D genome. We found that both the genetic diversity and evolutionary rate are more similar between 3D chromatin neighbors than between random pairs, and that both mutation and selection play essential roles in the observed correlated evolution. Chromatin organization has a major impact on regional genetic diversity and evolutionary rates in the genome. In addition, those (epi)genomic features that have a more significant influence on the evolution of local fragments also present higher levels of assortativity in the CIN, hence contributing more to the observed correlated evolution. The correlated evolution of large-scale fragments in the 3D genome inferred from our results will provide us new insights into the evolution and chromatin organization of the Arabidopsis genome and the organization of the eukaryotic genome in general.

It is well known that the organization of genes within eukaryotic genomes is not random at the one-dimensional scale (Hurst et al. 2004; Kosak and Groudine 2004) and linked genes often show comparable expression (Boutanaev et al. 2002; Williams and Bowles 2004) and evolve at similar rates (Williams and Hurst 2000). At the three-dimensional level, genes that are located close to each other present similar expression levels and protein functions (Ben-Elazar et al. 2013; Homouz and Kudlicki 2013; Diament et al. 2014). The present study extends our understanding of the organization of the eukaryotic genome by showing that adjacent large fragments carrying clusters of genes and other elements in the 3D genome evolve in a coordinated way. In addition, we focused on not only protein-coding genes and functional noncoding sequences but also on putatively non-functional sites, whose correlated evolution in the CIN has been shaped by mutation and linked selection.

It should be noted that the correlated evolution of large DNA fragments in the 3D genome of *A. thaliana* is not restricted to the 100 kb scale, it holds under the 50 kb resolution as well (supplementary text and supplementary figs. S11-S16, Supplementary Material online), indicating that the correlated evolution and its evolutionary and (epi)genomic determinants are general patterns at large scales in the Arabidopsis genome. Besides, using different Hi-C data, different filtration criteria, inter- or intra-chromosomal interactions only, or *A. thaliana* lineage specific divergence did not change the overall pattern (supplementary text and supplementary figs. S17-S26, Supplementary Material online).

The observed covariation of genetic diversity and evolutionary rates between 3D neighboring fragments may be explained by the evolutionary forces that act on them. First, shared evolutionary pressure exerted on interacting functional elements such as protein-coding sequences and noncoding regulatory elements may contribute to the observed correlated evolution. This similarity of selective constraints could in turn be derived from comparable expression levels and breadths due to similar regulatory mechanisms acting on them (Babaei et al. 2015). Second, the proteins encoded by DNA sequences residing in neighboring fragments may directly interact and adapt to changes in each other, resulting in a coordinated evolutionary rate. Using adjusted mutual information as a measure of the correlated mutation between SNPs in Arabidopsis, Perlaza-Jimenez and Walther (2018) demonstrated that sequences in physical contact at the 3D scale significantly overlap with pairs of genomic regions that are rich in correlated mutations, suggesting a potential role for compensatory mutations between 3D chromatin neighbors. Compensatory mutation does not exclusively occur between protein residues but can also occur between noncoding sequences (Landry et al. 2005; Takahasi et al. 2011; Barriere et al. 2012) and between coding and regulatory sequences (Hershberg and Margalit 2006). In our simulation analyses, the difference in *α* between the protein-coding genes residing in 3D neighboring fragments is not significantly smaller than that in the random networks, but the difference in *ω_a_* is significantly smaller than expected. Both *α* and *ω_a_* are measures of adaptive evolution; 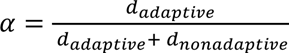, and 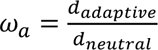, where *d_adaptive_* and *d_nonadaptive_* are the rates of adaptive and nonadaptive substitutions at putatively selected sites, respectively, while *d_neutral_* is the substitution rate at neutral sites. *α* measures the prevalence of adaptive evolution, while *ω_a_* reflects the strength or efficacy of positive selection. For the purpose of comparison between species or genomic regions, *ω_a_* is thought to be a more appropriate metric of adaptive evolution (Gossmann et al. 2012). Thus, in terms of the strength and efficacy of positive selection, our results suggest that genes residing in neighboring fragments at the 3D scale present more similar levels of adaptive evolution than expected.

For neutrally evolving sites, the genetic diversity is proportional to the effective population size and mutation rate (*θ* = 4*N_e_μ*), while the evolutionary rate is equal to the mutation rate (*k* = *μ*) (Kimura 1983). At both timescales, if a large fraction of nucleotides in a sequence are subjected to no or very weak selection, which may be the case for the Hi-C fragments because their long consecutive sequences contain a significant proportion of noncoding DNA, then the mutation rate may play a dominant role in determining the similarity of genetic diversity and substitution rates. Indeed, our PLS-PM analysis showed that mutation has a significant impact on the evolutionary rates of large-scale fragments, but its influence on genetic diversity is limited. On the other hand, the mutation rate itself varies both within and across genomes (Baer et al. 2007; Hodgkinson and Eyre-Walker 2011) and is affected by various (epi)genomic features, among which chromatin architecture is one of the decisive factors (Schuster-Bockler and Lehner 2012; Makova and Hardison 2015). Consistent with previous studies in humans, we found that mutation rate is elevated in genomic regions of closed chromatin and is suppressed in regions of open chromatin. The reduced mutation rate in regions with active gene expression may be attributed to transcription-coupled repair (Green et al. 2003), which is an additional repair mechanism that corrects errors in DNA before the next round of replication. Replication timing is another important determinant of mutation rate variation. Our results showed that mutation is reduced in regions with early replication and is enhanced in regions with late replication, corroborating the conclusion of previous studies (Makova and Hardison 2015). Furthermore, the combination of 39 (epi)genomic features can explain more than 60% of the variance of mutation rates across the genome at the 100 kb scale. In humans, more than 55% of the variance of somatic mutation rates can be explained by a combination of (epi)genomic features, whereas this proportion is reduced to 35% for germline mutation rate variation (Schuster-Bockler and Lehner 2012). The higher proportion of variance that can be explained by the (epi)genomic features in Arabidopsis compared to humans may stem from the fact that in plants, the soma and germen are not separated; therefore, the epigenomic features captured in somatic cells reflect the chromatin organization in germline cells to a greater extent in plant than in human cells.

Intriguingly, those (epi)genomic features that have a greater influence on regional mutation rates also present higher levels of assortativity in the network. Many features associated with active transcription (e.g., Pol II and its phosphorylated forms, H3K36me3, and H3K36ac) exhibit both highly negative correlations with mutation rates and high levels of assortativity in the network. The histone variants that are associated with heterochromatin (H3.1 and H2A.W) (Yelagandula et al. 2014) also exhibit very high levels of assortativity but highly positive correlations with mutation rate. However, the assortativity of replication timing is moderate (0.179 and 0.145 for early and late replication, respectively), suggesting that the role of replication in the observed correlated mutation rates is not as significant as that of transcription. Overall, our findings suggest that epigenetic modifications not only affect the structure of the 3D genome but also influence the correlated evolution between neighboring fragments in the CIN.

In addition to natural selection and mutation rates, random genetic drift is a pivotal determinant of genetic diversity (Ellegren and Galtier 2016). When the effective population size (*N_e_*) is small, genetic drift will play a principal role in determining the fate of a newly arisen mutations, thus reducing the efficiency of natural selection. Similar to other evolutionary parameters, *N_e_* is not constant across the genome (Charlesworth 2009). The within-genome variation of *N_e_* results from the differences in the strength of selection at linked sites (Charlesworth 2009), which in turn is associated with the local recombination rate and the density of selected sites (Cutter and Payseur 2013). Our simulation results showed that adjacent fragments do not present similar rates of recombination but exhibit comparable gene density (P = 0.911 and P < 0.001, respectively; supplementary fig. S27, Supplementary Material online). Using *π*_4_/*d*_4_ and *π_NNS_*/*d_NNS_* as proxies of *N_e_*, we found that spatially contacting fragments possess more similar *N_e_* than that expected by the CCS model (supplementary fig. S28A and B, Supplementary Material online). However, the difference in relative *N_e_* (denoted as *f*) estimated using varne (Zeng et al., 2019) between 3D interacting fragments is not significantly smaller than the random expectation (supplementary fig. S28C and D, Supplementary Material online). The rationale behind using *π*_4_/*d*_4_ and *π_NNS_*/*d_NNS_* as proxies of *N_e_* is that for neutral sites, nucleotide diversity is determined by 4*N_e_μ* while the substitution rate is determined by *μ*, thus 4*N_e_* = (4*N_e_μ*)/*μ* = *π_neutral_*/*d_neutral_*. The software varne incorporates a set of methods to jointly infer demographic history and detect variation in *N_e_* and *μ* between loci by utilizing polymorphism and divergence data (Zeng et al. 2019). Both approaches rely on within-species nucleotide diversity, which is shaped by the composite parameter *N_e_μ*. Thus, our results should be interpreted with caution, because *N_e_* and *μ* are inherently confounded. In any case, the analyses of the efficacy of negative and positive selection (measured as *π*_0_/*π*_4_ and *ω_a_*, respectively) indicated that the difference in *N_e_* is not large enough to generate considerable discrepancy in the efficacy of natural selection between spatially nearby fragments.

In the present study, we investigated the large-scale correlated evolution in only one species, the model plant *A. thaliana*. The genome of this species has some properties that differ from those of other eukaryotic organisms. First, the genome of *A. thaliana* is small and compact (∼125 Mb), and the density of coding sequences is relatively high compared to that in other species possessing a large genome. Second, as a predominantly selfing species, the effective recombination rate is rather low in *A. thaliana*. These two properties together affect linked selection in the genome, which in turn affects genetic diversity (Ellegren and Galtier 2016). Third, some characteristics of chromatin organization in the nucleus of *A. thaliana* differ from those in other species. For example, the topologically associated domain (TAD) is a conspicuous characteristic of the chromatin architecture in mammals and has also recently been found in rice (*Oryza sativa*) (Liu et al. 2017), whereas this structure is not prominent in *A. thaliana* (Dogan and Liu 2018). Preliminary analyses in *A. lyrata* and rice suggests that genome size and the density of coding sequences do not affect the correlated evolution between large DNA fragments in the CIN (supplementary text and supplementary figs. S29 and S30, Supplementary Material online). However, whether the observed pattern of correlated evolution can be extrapolated to other species requires further investigation.

## Materials and Methods

### Construction of the CIN

The raw reads of two Hi-C datasets in *A. thaliana* (Wang et al. 2015; Liu et al. 2016) were downloaded from the NCBI Sequence Read Archive. Since these two datasets represent the average chromatin architecture in the same tissue at the same developmental stage (i.e., the aerial portions of 10-day-old seedlings), we combined the short reads from these two studies to detect chromatin interactions, as did in Liu et al. (2016). The reads were mapped to the *A. thaliana* genome (TAIR10) using bowtie2 (Langmead and Salzberg 2012) with default parameters, and PCR duplicates and reads mapped to multiple locations in the genome were removed from downstream analysis. The contact matrix at the 100 kb resolution was obtained using HiC-Pro (Servant et al. 2015). We applied two different strategies to detect significant intra- and inter-chromosomal contacts. For intra-chromosomal interactions, we used Fit-Hi-C (Ay, Bailey, et al. 2014) to identify significant contacts. We used an FDR cutoff of 0.05 to remove low-confidence intra-chromosomal contacts. For contacts between chromosomes, it is difficult to model the null distribution of contacts since there is no trend of decreasing interaction probability with increasing linear distance. Thus, we applied a uniform probability model to calculate the P-values of interactions as described previously (Duan et al. 2010; Xie et al. 2016). We carried out multiple test corrections and applied a stringent FDR cutoff of 0.0001 to remove the noise of the Hi-C data. The CIN was constructed with both intra- and inter-chromosomal high-confidence contacts using the igraph package (version 1.2.4) (Csardi and Nepusz 2006) in R.

Because the DFE-alpha method is not suitable for estimating the rate of adaptive evolution with a small number of genes, we removed individual 100 kb fragments with no more than 15 protein-coding genes from the chromatin network construction procedure and all downstream analyses. Furthermore, we removed fragments for which more than 80% of nucleotides could not be aligned to *A. lyrata* or masked as repeat sequences in either the *A. thaliana* genome or their corresponding aligned sequences in *A. lyrata*.

### Calculation of genetic diversity and divergence

Polymorphism data for *A. thaliana* were downloaded from the 1001 Genomes website (http://1001genomes.org), which contains genome-wide SNPs and indels from 1,135 globally distributed Arabidopsis accessions (Alonso-Blanco et al. 2016). We filtered out SNPs with more than two alleles and insertion/deletion (indel) polymorphisms, resulting in 10,707,430 biallelic SNPs. The averaged nucleotide diversity (*π*) was calculated for each nonoverlapping 100 kb window to represent the genetic diversity of fragments. Additionally, the average *π* for 0-fold nonsynonymous sites, 4-fold synonymous sites, putatively functional noncoding sequences, and putatively neutral noncoding sequences were calculated for each fragment. The putatively functional noncoding sequences were defined as the conserved noncoding sequences (CNS) identified by Haudry et al. (2013) plus the untranslated regions (5’UTR and 3’UTR), abbreviated as CNS+UTR. The putatively neutral noncoding sequences (NNSs) were defined as the noncoding sequences excluding the CNS+UTR sequences and nucleotides that are identified as belonging to DHSs in PlantTFDB 4.0 (http://planttfdb.cbi.pku.edu.cn) (Jin et al. 2017).

Regarding the evolutionary rates of fragments, we calculated the divergence between *A. thaliana* and *A. lyrata* for each fragment, as well as the divergence at sites with different functional constraints in each window. Whole-genome alignment between *A. thaliana* and *A. lyrata* was carried out using LASTZ (Harris 2007), followed by chaining and netting (Kent et al. 2003). Blocks shorter than 100 bp were discarded. Any site that was masked as part of a repeat sequence in either *A. thaliana* or *A. lyrata* was excluded from the divergence analysis. We used the Nei-Gojobori method (Nei and Gojobori 1986) to calculate divergence between the two species.

### Inference of natural selection

Several parameters were calculated to quantify the strength and efficiency of negative selection and positive selection for each fragment in the CIN. First, we quantified the strength of selective constraints as %(*DAF* < 0.005) in the global collection of Arabidopsis in each window. Ancestral alleles were derived from whole-genome alignment with *A. lyrata*. Second, we calculated the ratio of 0-fold nonsynonymous versus 4-fold synonymous nucleotide diversity (*π*_0_/*π*_4_) and the ratio of nucleotide diversity at putatively functional noncoding sites to diversity at neutral noncoding positions (*π_CNS+UTR_*/*π_NNS_*) in each window as the strength or efficiency of purifying selection on protein-coding genes and functional noncoding sequences, respectively.

We also computed the fraction of adaptive nucleotide substitutions (*α*) and the rate of adaptive evolution relative to the rate of neutral evolution (*ω_a_*) using the DFE-alpha approach (Eyre-Walker and Keightley 2009) for 0-fold degenerate sites. Because DFE-alpha is inappropriate for estimating DFE and *α* using large samples (Kim et al. 2017), we restricted our analyses to either the ancestral population on the Iberian Peninsula (22 accessions) or the more recent Spanish population (50 accessions). DFE was estimated under a 2-epoch model in which a stepwise population size change was applied. The Jukes-Cantor model was applied to correct the substitution rate between species, and polymorphic sites were removed when calculating *α* and *ω_a_*.

### Inference of effective population size

We calculated *π*_4_/*d*_4_ and *π_NNS_*/*d_NNS_* for each fragment as its proxies of *N_e_*. In addition, we applied varne (Zeng et al. 2019) to detect variation in *N_e_* between loci. We used the simplified model with divergence data, which is appropriate for data with a large number of loci. The folded site frequency spectrum at neutral sites in each fragment is computed from polymorphism data in either the relict or Spanish population. Divergence information at neutral sites is inferred from *A. thaliana*-*A. lyrata* alignment. We ran the program with the LBFGS and TNEWTON algorithms separately, each with 100 independent searches. For the Spanish population, the correlation of relative *N_e_* between the two different algorithms is very low, and the estimates from different searches with top log likelihoods are drastically different, indicating problems in finding the best values. Such problems are not found in the relict population, therefore, we only present results obtained with polymorphism information in the relict population.

### Simulation (P-value computation)

We applied a conservative empirical null model, the cyclic chromosome shift model (Diament et al. 2014), to test whether fragments in direct contact in the CIN evolve in a coordinated way and whether chromatin organization plays a role in the correlated evolution of 3D neighbors. Specifically, we preserved the spatial configuration of the Hi-C network and randomly shifted the location of fragments along their respective chromosomes. By doing so, the linear adjacency between fragments along the “circular” chromosomes and topology of the network remain unchanged, but the contact relationships between fragments in the 3D structure are perturbed (supplementary fig. S1B, Supplementary Material online). We generated 1,000 simulated networks and calculated the P-value as the number of networks in which the mean value of the difference of a parameter between neighboring nodes is smaller than that in the actual spatial conformation. We also employed other empirical null models: degree-preserving rewiring of the network and cyclic genome shift, in which fragments are allowed to rotate and move between different chromosomes.

### Genome-wide feature data processing and regression analysis

Short reads for DHSs, histone acetylations, histone methylations, the occupancy of histones and Pol II and their variants were downloaded from NCBI (see supplementary table S2 for a complete list of source references and SRA accession numbers), and mapped to the Arabidopsis reference genome (TAIR10). PCR duplicates were filtered out. The relative density of a feature in a 100 kb window was calculated as the log2-transformed and normalized read count with a mean of 0 and a standard deviation of 1. Gene density was calculated as the total number of nucleotides covered by genes (including introns) in a window divided by the length of the window. Gene-level recombination rates were retrieved from Marais et al. (2004) and averaged for each 100 kb window. The GC content was computed as the fraction of G and C nucleotides in a window. The annotation of transposable elements (TEs) in *A. thaliana* was downloaded from The Arabidopsis Information Resource (TAIR) website (www.arabidopsis.org), and CpG islands in the Arabidopsis genome were identified using CpGIScan (Jian et al. 2017). The coverage of TEs and CpG islands was calculated as the fractions of nonoverlapping TEs and CpG islands in a window, respectively. Gene order conservation scores were retrieved from the Ensembl Plants database (http://plants.ensembl.org) and averaged for each window. The coverage of replication origins was calculated using data from Vergara et al. (2017). The replication timing of three categories (i.e., early, mid, and late) was computed using data from Concia et al. (2018).

To determine which genomic features that correlate with and contribute to the variation of genetic diversity and evolutionary rates in the Arabidopsis genome, we first calculated the Pearson’s correlation coefficients of each evolutionary parameter with the above 39 genomic features using the “cor.test” function in R. We then performed multiple linear regression analyses using the genomic features as independent variables and each focal evolutionary parameter as a dependent variable with forward stepwise selection using the “regsubsets” function of the leaps R package. A PLS-PM analysis was carried out to infer the underlying relationships among (epi)genomic features, evolutionary forces, genetic diversity, and evolutionary rates using the R package plspm (Sanchez 2013). We used two latent variables (group I and group II) to represent the 39 (epi)genomic features; group I summarizes the features that present negative correlations with genetic diversity and evolutionary rates, and group II represents the features that are positively correlated with diversity and divergence.

### Network analysis

We calculated the assortativity of each of the 39 genomic features in the CIN using the “assortativity” function of the igraph R package. For continuous variables, assortativity is defined as

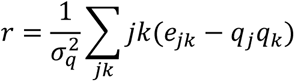

where *q_i_* = ∑_*j*_ *e_ij_*, *e_ij_* is the fraction of edges connecting vertices of type *i* and *j*, and *σ_q_* is the standard deviation of *q* (Newman 2002). If the assortativity of a feature is high, it means that connected nodes tend to exhibit similar values.

## Supplementary Material

Supplementary tables S1-S2 and figures S1-S30 are attached separately.

## Acknowledgments

We thank Bing Su, Kai Zeng, and Tian Tang for valuable discussions regarding this work. This work was supported by the Fundamental Research Funds for the Central Universities of China (Z109021804) to Y.Y. and the National Natural Science Foundation of China (31670321), the Hundred Talents Program of Shaanxi Province of China (A289021612), and the Fund of Northwest A&F University (Z111021404) to R.Y.

## Author Contributions

R.Y. conceived and designed the research; Y.Y. and Z.L. collected and analyzed the data with input from all authors; Y.Y. wrote the manuscript with input from R.Y. and Z.L.

