## Supplementary for "Correlated evolution of large DNA fragments in the 3D genome of *Arabidopsis thaliana*"

**Figure S19. Distribution of difference in evolutionary parameters between interacting fragments using the CGS null model when only inter- or intra-chromosomal interactions are considered. .34**

**Figure S30. Correlated evolution between 3D interacting fragments in rice at the 100 kb scale..47**

### Supplementary text

#### 1. Correlated evolution of spatially interacting fragments at the 50 kb scale

To confirm that our conclusions are not restricted to the scale (100 kb) that we presented in the main text, we repeated most of the analyses at a 50 kb resolution, except for the assessment of the adaptive rate because there are too few genes in a window to obtain reliable estimates for this parameter. The Arabidopsis genome was split into 2,385 nonoverlapping windows, and fragments with less than eight genes and more than 80% unmappable sites were removed. All the results are quantitatively similar to the results at a 100 kb resolution (Supplementary figs. S11-S16), suggesting that the correlated evolution and its evolutionary and (epi)genomic determinants are general patterns at large scales in the Arabidopsis genome.

We only investigated the correlated evolution of large fragments at the 100 kb and 50 kb scales in this study. Basically, we can explore the same questions at larger or smaller scales. *A. thaliana* has a small genome of ~125 Mb. Using a larger scale, saying at 1 Mb scale, then the number of nonoverlapping windows is small,  and each fragment is more likely to be connected by other fragments in the 3D genome. Therefore, the resolution might be too coarse to get useful insights. In an extreme case, if we study chromosome-wise interactions, all chromosomes would be spatially close to each other. On the other hand, at a lower scale than 50 kb, the genes embedded in a fragment would be decreased on average, making a comprehensive assessment of the many relationships we investigated in the main text unreliable. Overall, we chose the two scales because we are mainly interested in large scale correlated evolution in the 3D genome and these resolutions are also appropriate scales for such a study in Arabidopsis.

It should be pointed out that there might be variation in evolutionary parameters (e.g., mutation rates) within large fragments. To assess this variation, we divided each 100 kb fragment into two 50 kb nonoverlapping small fragments, and randomly sample one small fragment in each 100kb fragment. We carried out 1,000 random sampling, and each time we computed the correlation coefficient between the large and small resolutions. For mutation rates, the average Pearson’s correlation coefficient between 100 kb and 50 kb fragments is 0.916, suggesting that the variation in mutation rate within fragments at broad scale is limited. Besides, studies on variation in evolutionary parameters in different organisms showed that the general pattern is usually qualitatively similar for analyses at different genomic scales (Slotte et al. 2011; Terekhanova et al. 2017; Smith et al. 2018 ).

### 2. Assessing the effect of using different Hi-C data and different filtration criteria

To assess the effect of Hi-C dataset, we constructed a CIN at 100 kb using another independent experiment (Feng et al. 2014). This dataset was obtained from four-week-old rossete leaves, which is differ from the data used in the main text in term of developmental stage and tissue. The pattern of correlated evolution is the same as that in the main text (Supplementary fig. S17).

By removing fragments with few genes and poor mappability, we excluded approximately 20% (236 out of 1,193) of 100 kb windows from our analyses. Most excluded fragments are located at the centromeric regions. To evaluate the influence of this filtration process, we repeated parts of the analyses without omitting any fragments. Our results showed that the differences in nucleotide diversity, overall divergence between species, divergence at neutral sites, and $\%(DAF < 0.005)$ between 3D neighboring fragments are significantly smaller than that in the CCS simulations (Supplementary fig. S18), which is consistent with the results in the main text. The pattern of correlated evolution is retained when we keep inter- or intra-chromosomal interactions only (Supplementary fig. S19).

### 3. Assessing the effect of using *A. thaliana* lineage specific divergence

We used the divergence between *A. thaliana* and *A. lyrata* as measurements of evolutionary rate or mutation rate in *A. thaliana*. By doing this, we assume that the evolutionary rate or mutation rate is symmetric between orthologous regions of *A. thaliana* and *A. lyrata*. However, this assumption may be violated. To investigate this bias, we performed three-way alignment with an additional outgroup, *Capsella rubella,* using multiz (Blanchette et al. 2004). Divergent sites that are specific to the *A. thaliana* lineage were identified as where *A. lyrata* and *C. rubella* had the same nucleotide and this nucleotide differed in the *A. thaliana* sequence. Ancestral alleles were identified as the alleles where *A. lyrata* and *C. rubella* had the same nucleotide, and this nucleotide was one of the two alleles in *A. thaliana*. We repeated all the analyses presented in the main text at 100 kb scale. The average length of analyzable sites in 100 kb windows drops from 76.6 kb to 68.5 kb, but the results are generally the same for all analyses (Supplementary figs. S20-S26).

#### 4. Correlated evolution of spatially interacting fragments in *A. lyrata*

To evaluate the generality of the correlated evolution between 3D neighboring fragments, we constructed CINs in another two plant species. The CIN at the 100 kb scale in *A. lyrata* was constructed with the Hi-C data generated by Zhu et al. (2017) using the same methods as in *A. thaliana*. Because *A. lyrata* has a relatively sparse genome, in order to not filter out too many windows, instead of removing fragments with less than 16 genes, we removed fragments with less than eight genes and more than 80% of unmappable sites. The average length of mappable sites in 100 kb windows is 38.4 kb. We calculated *A. lyrata* lineage specific divergence using *A. lyrata-A. thaliana-C. rubella* three-way alignment for each fragment, and compared the overall divergence between connected fragments in the actual CIN and that in the simulated networks. The result shows that the difference between spatially contacting fragments is significantly smaller than that expected in the CCS model (Supplementary fig. S29A), and the difference increases with an increase in topological distance in the network (Supplementary fig. S29B). No comparable polymorphism database was found in *A. lyrata*, thus only between species divergence was assessed.

#### 5. Correlated evolution of spatially interacting fragments in rice

The Hi-C data generated by Dong et al. (2018) was used to construct the CIN at 100 kb in rice (*Oryza sativa*). Fragments with less than eight genes and more than 80% unmappable sites were discarded from network construction. Nucleotide diversity for each 100 kb nonoverlapping window was computed using biallelic SNPs in the Geng/Japonica population from the 3,000 Rice Genome Project (Wang et al. 2018). Divergence in each 100 kb window was calculated with *O. sativa-O. meridionalis* whole genome alignment. Our CCS simulations show that for both nucleotide diversity and divergence, the differences between spatially contacting fragments are significantly smaller than that expected in the CCS model (Supplementary fig. S30A and B), and the difference increases with an increase in topological distance in the network (Supplementary fig. S29C and D).

### Supplementary tables

**Table S1.** Summary statistics for the overall and four classes of sites at the100 kb scale.

| | $\pi$ (× $10^{-3}$) | Divergence | Alignable Sites (kb) |
|---|---|---|---|
| **All** | 4.54 ± 1.98 | 0.102 ± 0.018 | 76.64 ± 11.78 |
| **0-fold** | 2.11 ± 1.29 | 0.038 ± 0.016 | 19.82 ± 4.41 |
| **4-fold** | 7.09 ± 4.00 | 0.139 ± 0.026 | 4.53 ± 1.03 |
| **CNS+UTR** | 4.31 ± 2.00 | 0.095 ± 0.016 | 13.17 ± 3.63 |
| **NNS** | 5.50 ± 2.06 | 0.148 ± 0.022 | 23.29 ± 3.84 |

Note: All – all analyzable sites in a window; 0-fold – 0-fold degenerate sites; 4-fold – 4-fold degenerate sites; CNS+UTR – conserved noncoding sequences and untranslated sequences; NNS – putatively neutral noncoding sites.

**Table S2.** Data sources.

| Data type | Sample | SRA accession number | Reference |
|---|---|---|---|
| ***Arabidopsis thaliana*** | | | |
| Hi-C (DpnII) | 10-day-old seedlings | SRR2626163<br>SRR2626429 | Liu et al. 2016 |
| Hi-C (DpnII) | 10-day-old seedlings | SRR1029603<br>SRR1029605 | Wang et al. 2015 |
| Hi-C (HindIII) | Four-week-old rosette leaves | SRR1504819 | Feng et al. 2014 |
| DHSs | Two-week-old leaves | SRR388657<br>SRR388658<br>SRR388659 | Zhang et al. 2012 |
| DHSs | Two-week-old flowers | SRR388660<br>SRR388661 | Zhang et al. 2012 |
| H3K27ac | leaves | SRR1509474 | Zhu et al. 2015 |
| H3K4me1 | leaves | SRR1509477 | |

| | | | |
|---|---|---|---|
| H3 | 12-day-old leaves | SRR1848390<br>SRR1848384<br>SRR1848387 | Moreno-Romero et al. 2016 |
| H3K27me1 | 20-day-old leaves | SRR1848395 | |
| H3K27me3 | 20-day-old leaves | SRR1848399<br>SRR1848402 | |
| H3K9me2 | 20-day-old leaves | SRR1848404<br>SRR1848405 | |
| H3K4me3 | 29-day-old leaves | SRR1964977 | Brusslan et al. 2015 |
| H3K9ac | 30-day-old leaves | SRR1964985 | |
| H3K4me2 | Three-week-old rosette leaves | SRR2637923 | Chen, Luo, et al. 2017 |
| H3K4me3 | Three-week-old rosette leaves | SRR2637924 | |
| H3K36ac | 35-day-old leaves | SRR2932297<br>SRR2932298<br>SRR2932293 | Mahrez et al. 2016 |
| H2A | Three-week-old leaves | SRR988543 | Yelagandula et al. 2014 |
| H2A.W | Three-week-old leaves | SRR988544 | |
| H2A.X | Three-week-old leaves | SRR988545 | |
| H2A.Z | Three-week-old leaves | SRR988546 | |
| H3K27me3 | 12-day-old seedlings | SRR3286785 | Chen, Li, et al. 2017 |
| H3K14ac | 12-day-old seedlings | SRR3286787<br>SRR5224476 | |
| H3K18ac | 12-day-old seedlings | SRR3286788<br>SRR5224477 | |
| H4K5ac | 12-day-old seedlings | SRR3286789<br>SRR5224478 | |
| H4K8ac | 12-day-old seedlings | SRR3286790<br>SRR5224479 | |

| | | | |
|---|---|---|---|
| H4K12ac | 12-day-old seedlings | SRR3286791<br>SRR5224480 | |
| H4K16ac | 12-day-old seedlings | SRR5224481<br>SRR3286792 | |
| H3.1 | Three-week-old leaves | SRR6207345 | Lu et al. 2018 |
| H3.3 | Three-week-old leaves | SRR6207346 | |
| PolIISer2P | 10-day-old seedlings | SRR5313784 | Liu et al. 2018 |
| PolIISer2P input | 10-day-old seedlings | SRR5313785 | |
| PolIISer5P | 10-day-old seedlings | SRR5313788 | |
| PolIISer5P input | 10-day-old seedlings | SRR5313789 | |
| PolII | 10-day-old seedlings | SRR5313790<br>SRR5313791 | |
| H3K36me3 | 21-day-old leaves | SRR5113999 | Sanders et al. 2017 |
| H4K16ac | Two-week-old leaves | SRR2046531 | Lu et al. 2015 |
| H3K23ac | Two-week-old leaves | SRR2046530 | |
| ***Oryza sativa japonica*** | | | |
| Hi-C (MobI) | 21-day-old leaves | SRR6470741<br>SRR6470742<br>SRR6470743<br>SRR6470744<br>SRR6470745 | Dong et al. 2018 |
| ***Arabidopsis lyrata*** | | | |
| Hi-C (DpnII) | 30-day-old seedlings | SRR5155580<br>SRR5155617 | Zhu et al. 2017 |

### Supplementary figures

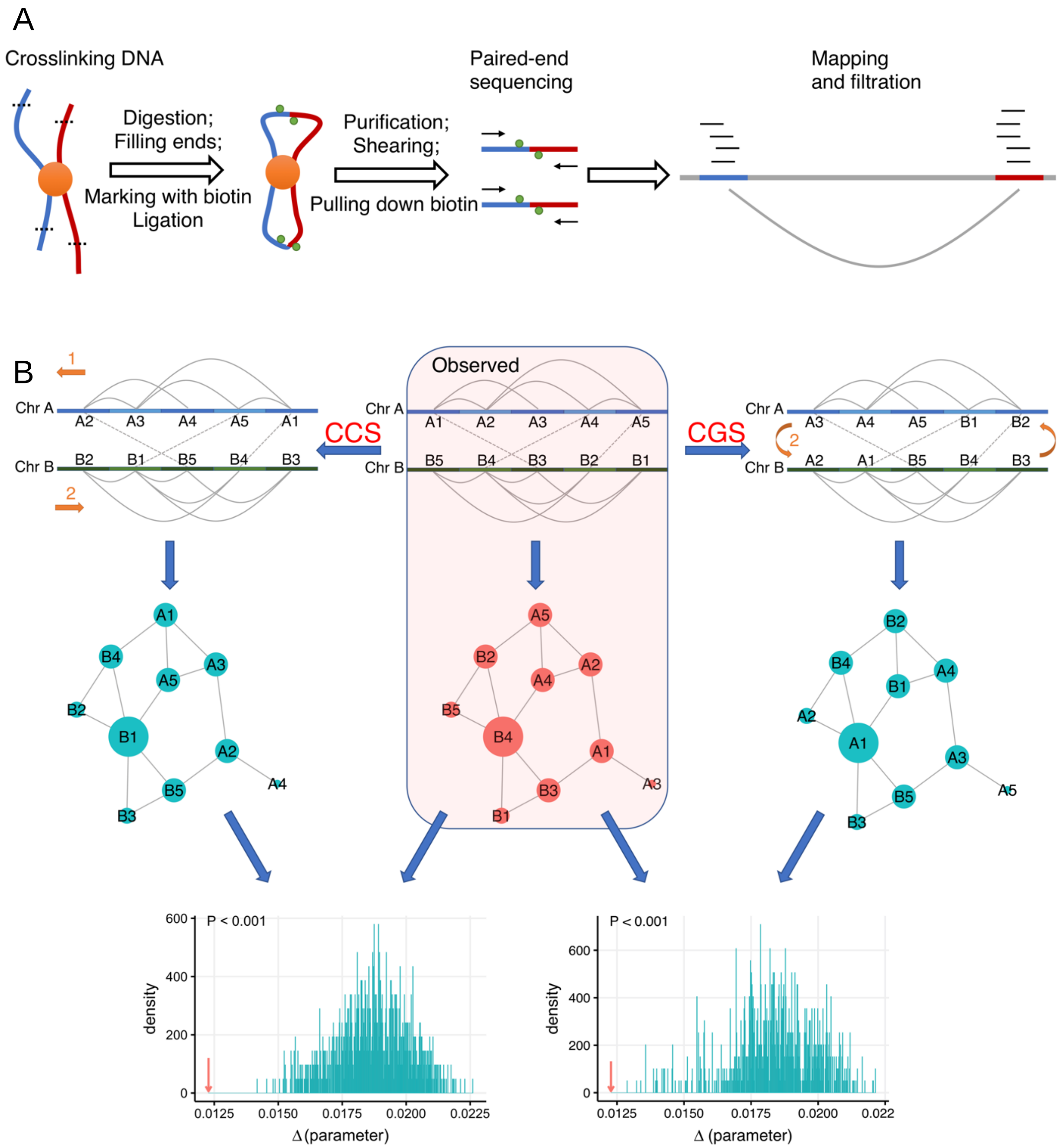

**Figure S1.** Overview of Hi-C and schematic illustration of CIN and simulation models. (A) Overview of Hi-C technology. In a typical Hi-C experiment, spatially adjacent chromatin segments (blue and red lines) mediated by protein complex (orange circle) are first cross-linked with formaldehyde and digested with restriction enzyme (black dash lines represent cutting sites), then the sticky ends are filled in with nucleotides (one of which is marked with biotin; green dots), after that ligation is performed to create chimeric molecules that include interacting DNA pairs, lastly DNA is purified and sheared and those chemeric DNA labeled with biotin is isolated and sequened. By mapping reads to the reference genome (gray horizontal line) and applying filtration criteria, significant interactions between fragments are identified (gray curve). (B) Schematic illustration of CIN and simulation models. In the hypothetical chromosomal conformation identified by Hi-C (top middle), intra- and inter-chromosomal interactions between fragements are represented with solid and dash lines. Note that for the sake of simiplicity, there are interactions between neighbors at the linear scale in the illustration, but in the real data, connections between linear neighbors are removed form CIN construction. The spatial relationship between fragments can be concepualized as a network (the central shaded part), in which a node denotes a fragment while an edge represents the spatial proximity between two fragments. The node degree denotes the number of interactions of a fragment. The sizes of the circles are proportion to the node degree in the figure. For example, fragment B4 has a high degree of five, while A3 has a low degree of one. The two simulation null models are shown at the left and right sides. In the CCS model (top left), fragments are shifted along their respective chromosome by a random positive integer between one and the coreesponding total number of fragments in that chromosome. For example, all fragments on chromosome A are shifted one position along its “circular” chromosome, while all fragments on chromosome B are shifted two positions. In the resulting CIN (middle left), the linear adjacency between fragments along the “circular” chromosomes and the topology of the network remain unchanged, but the contact relationships between fragments in the CIN are perturbed. The CGS model (top right) is similar to the CCS model, but allows fragments to move to other chromosomes. In this example, all fragments are shifted two positions along the “concatenated circular” chromosome (A1-A5+B1-B5). The statistical significance of a parameter is assessed by comparing the actural value and the simulated values (lower part).

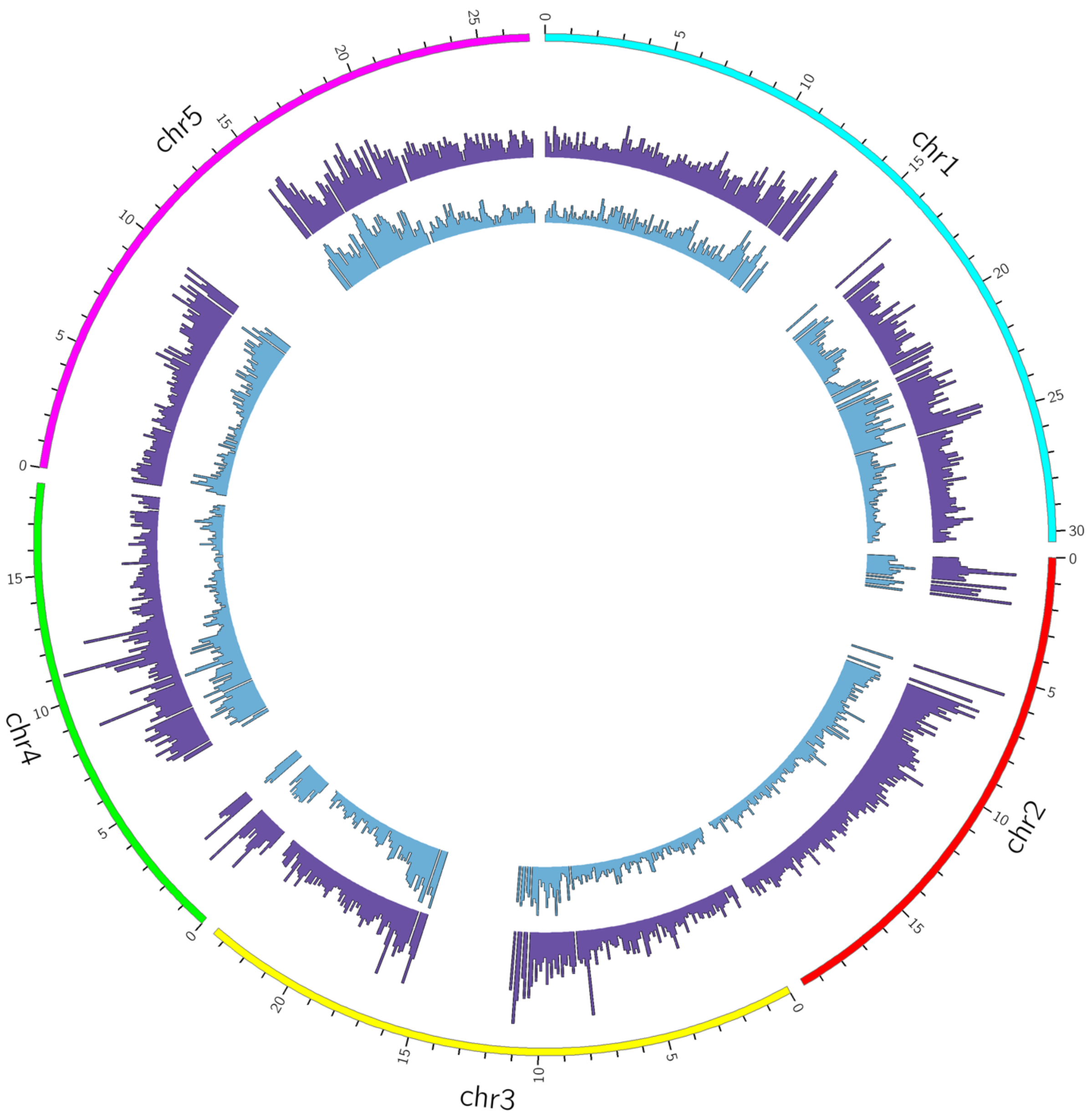

**Figure S2.** Average nucleotide diversity and evolutionary rates across the Arabidopsis genome at the 100 kb scale. Blue bars: nucleotide diversity; purple bars: evolutionary rates. Note that we filter out windows with no more than 15 genes and less than 20% mappable sites.

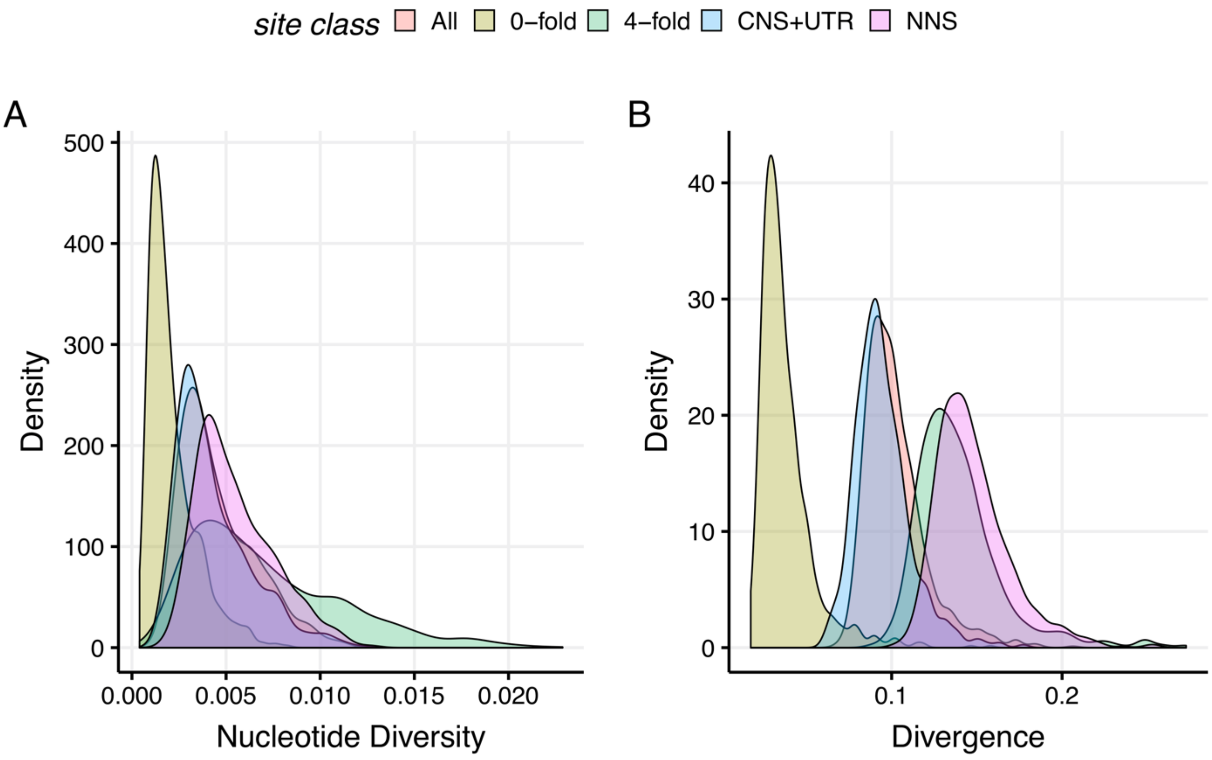

**Figure S3.** Distribution of the average within-species diversity and between-species divergence for the overall and four classes of sites at the 100 kb resolution. (A) Nucleotide diversity; (B) Divergence between *A. thaliana* and *A. lyrata*.

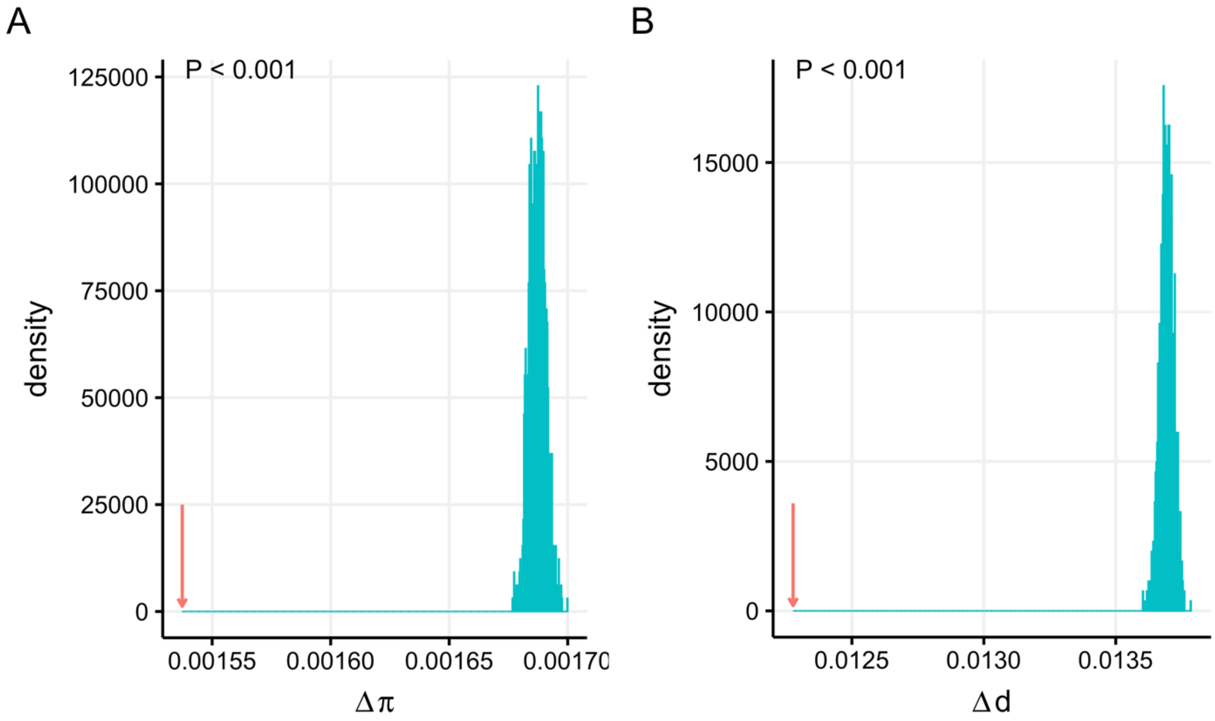

**Figure S4.** Distribution of difference in evolutionary parameters between connected fragments in the degree-preserving simulations. Red arrows show the observed differences between contacting fragments in the real data. (A) Nucleotide diversity; (B) Divergence between *A. thaliana* and *A. lyrata*.

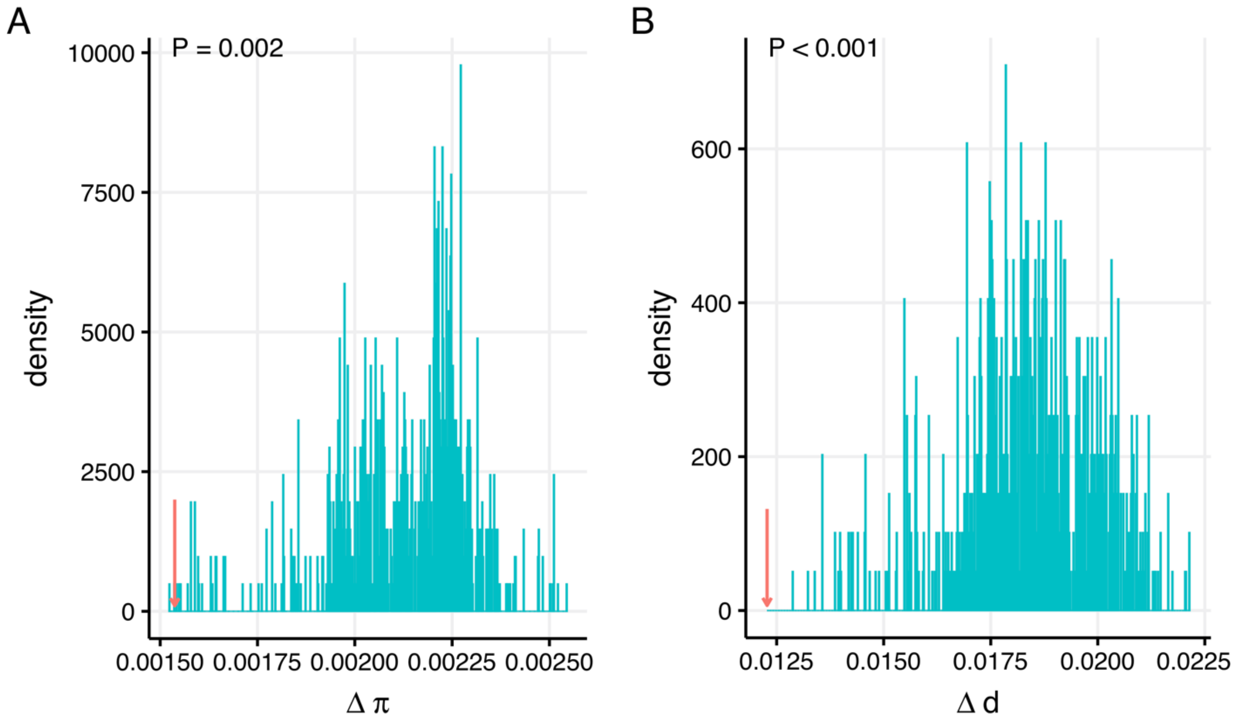

**Figure S5.** Distribution of difference in evolutionary parameters between interacting fragments using the CGS null model. Red arrows show the observed differences between contacting fragments in the real data. (A) Nucleotide diversity; (B) Divergence between *A. thaliana* and *A. lyrata*.

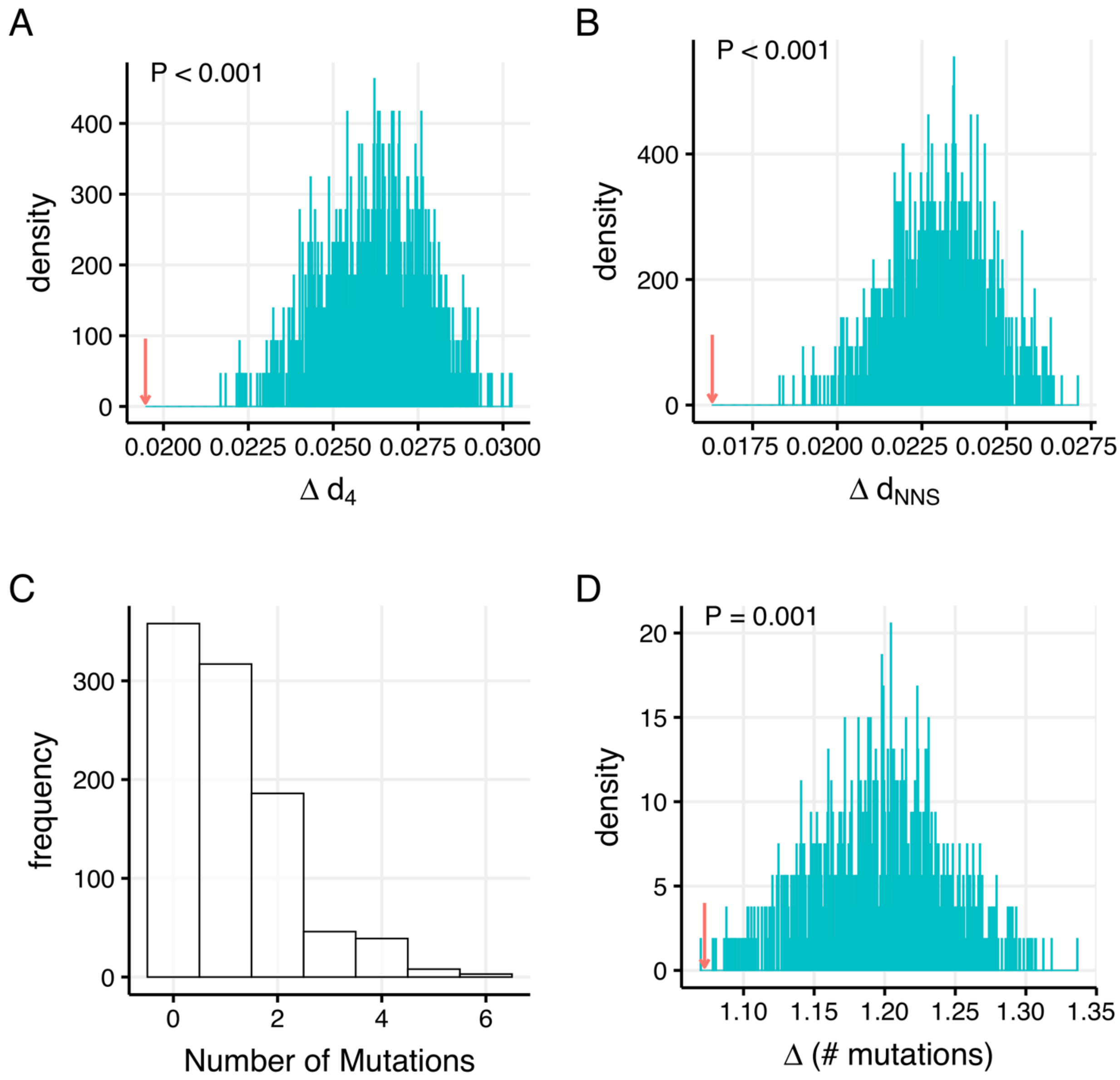

**Figure S6.** Distribution of difference in local mutation rates between interacting fragments using the CCS null model. Red arrows show the observed differences between contacting fragments in the real data. (A), (B), and (D) show the distributions in local mutation rates measured using divergence at 4-fold degenerate sites, divergence at putatively neutral noncoding sites, and the number of *de novo* mutations, respectively; (C) The distribution of the number of *de novo* mutations at the 100 kb scale.

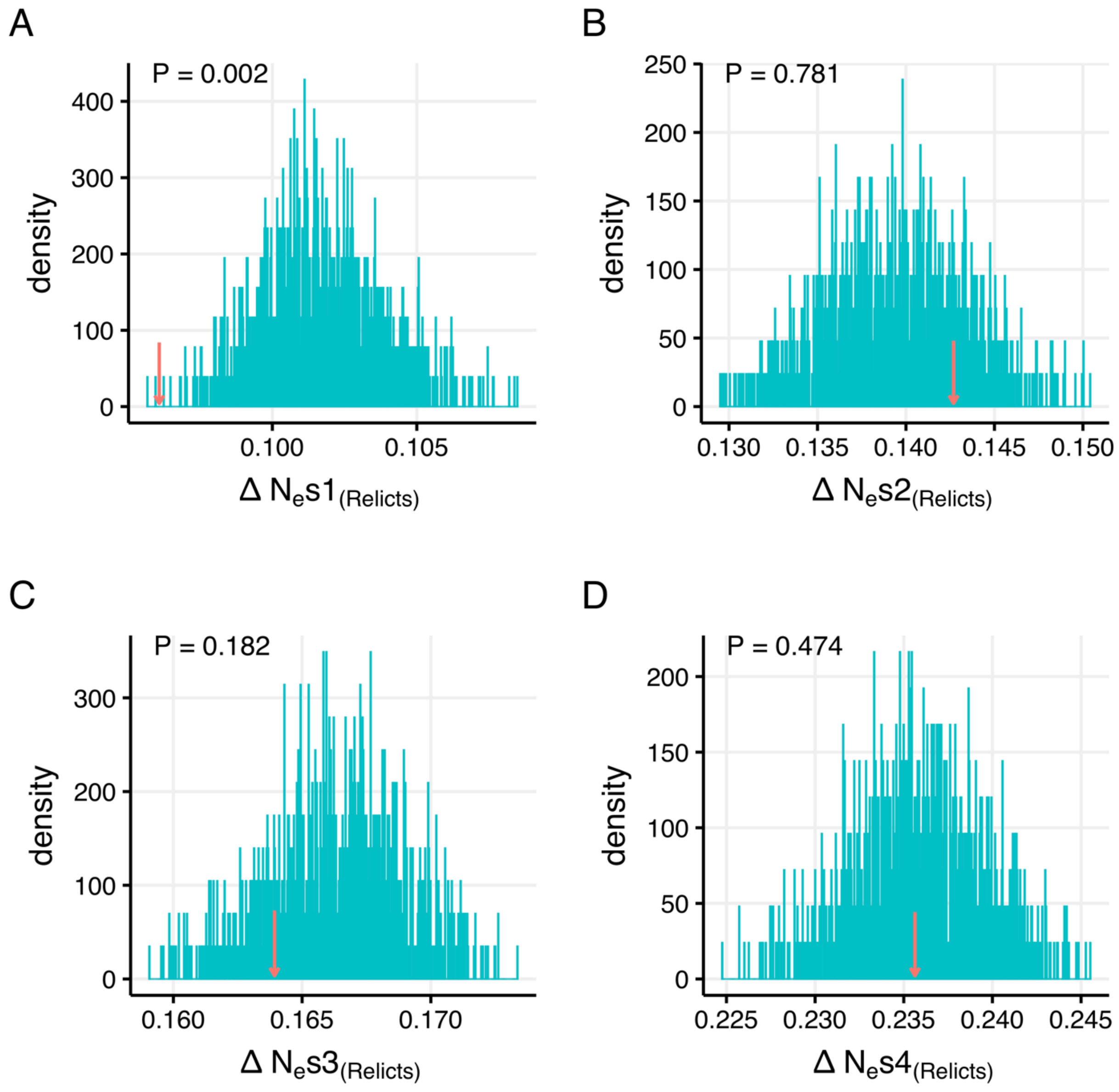

**Figure S7.** Distribution of difference in different categories of $N_e s$ derived from the relict population between interacting fragments using the CCS null model. Red arrows show the observed differences between contacting fragments in the real data. (A) $N_e s1: 0 < N_e s < 1$; (B) $N_e s2: 1 < N_e s < 10$; (C) $N_e s3: 10 < N_e s < 100$; (D) $N_e s4: N_e s > 100$.

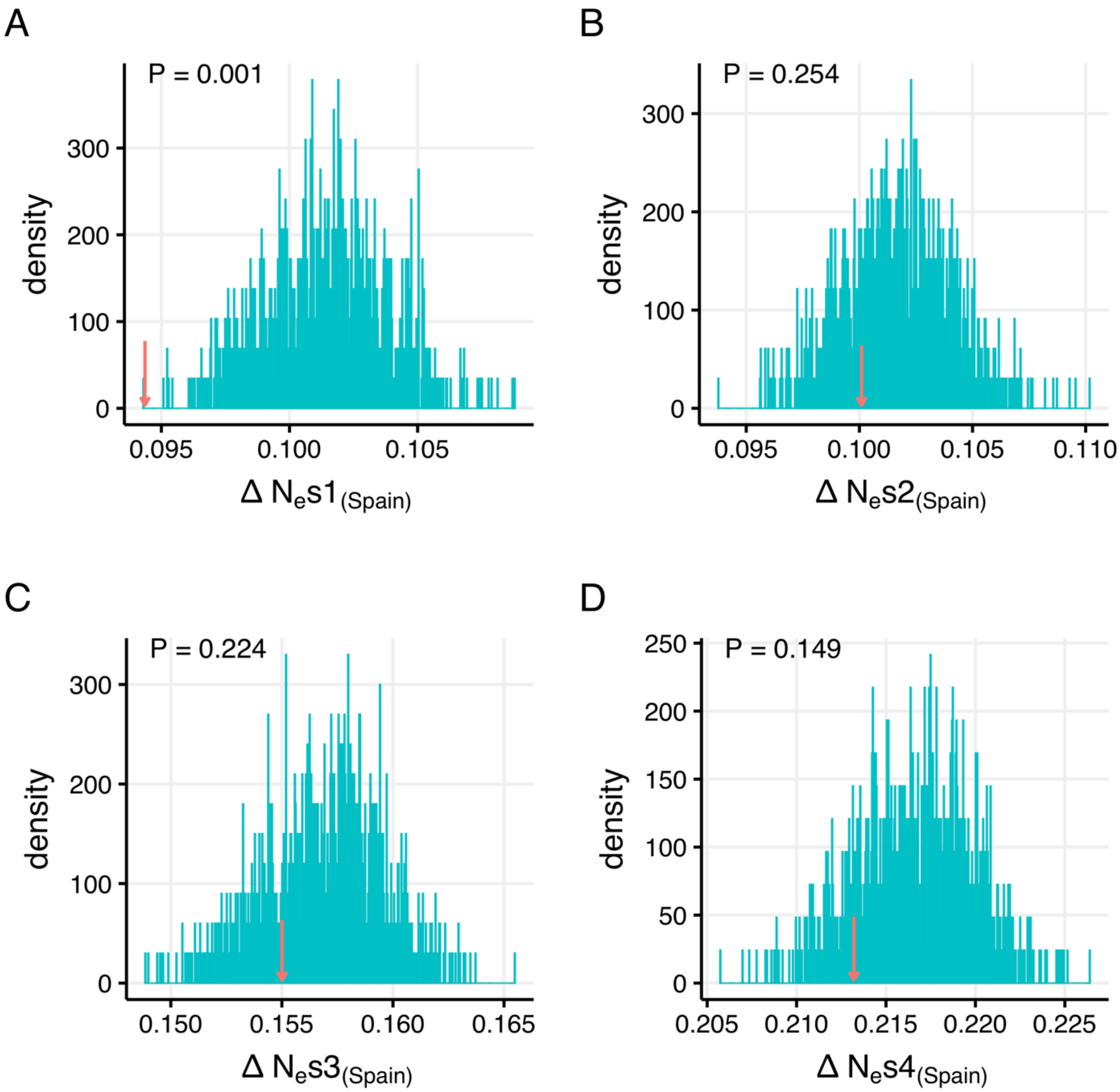

**Figure S8.** Distribution of difference in different categories of $N_e s$ derived from the Spanish population between interacting fragments using the CCS null model. Red arrows show the observed differences between contacting fragments in the real data. (A) $N_e s1: 0 < N_e s < 1$; (B) $N_e s2: 1 < N_e s < 10$; (C) $N_e s3: 10 < N_e s < 100$; (D) $N_e s4: N_e s > 100$.

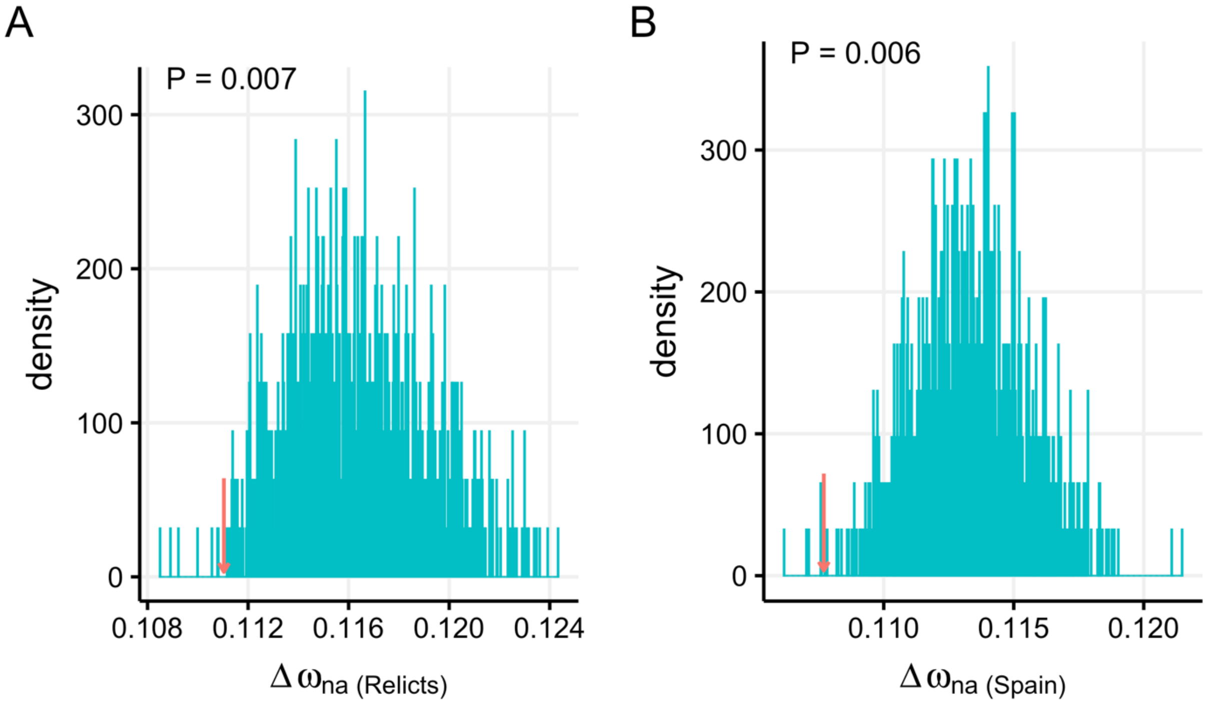

**Figure S9.** Distribution of difference in $\omega_{na}$ between interacting fragments using the CCS null model. Red arrows show the observed differences between contacting fragments in the real data. (A) $\omega_{na}$ derived from polymorphism in the relict population; (B) $\omega_{na}$ derived from polymorphism in the Spanish population.

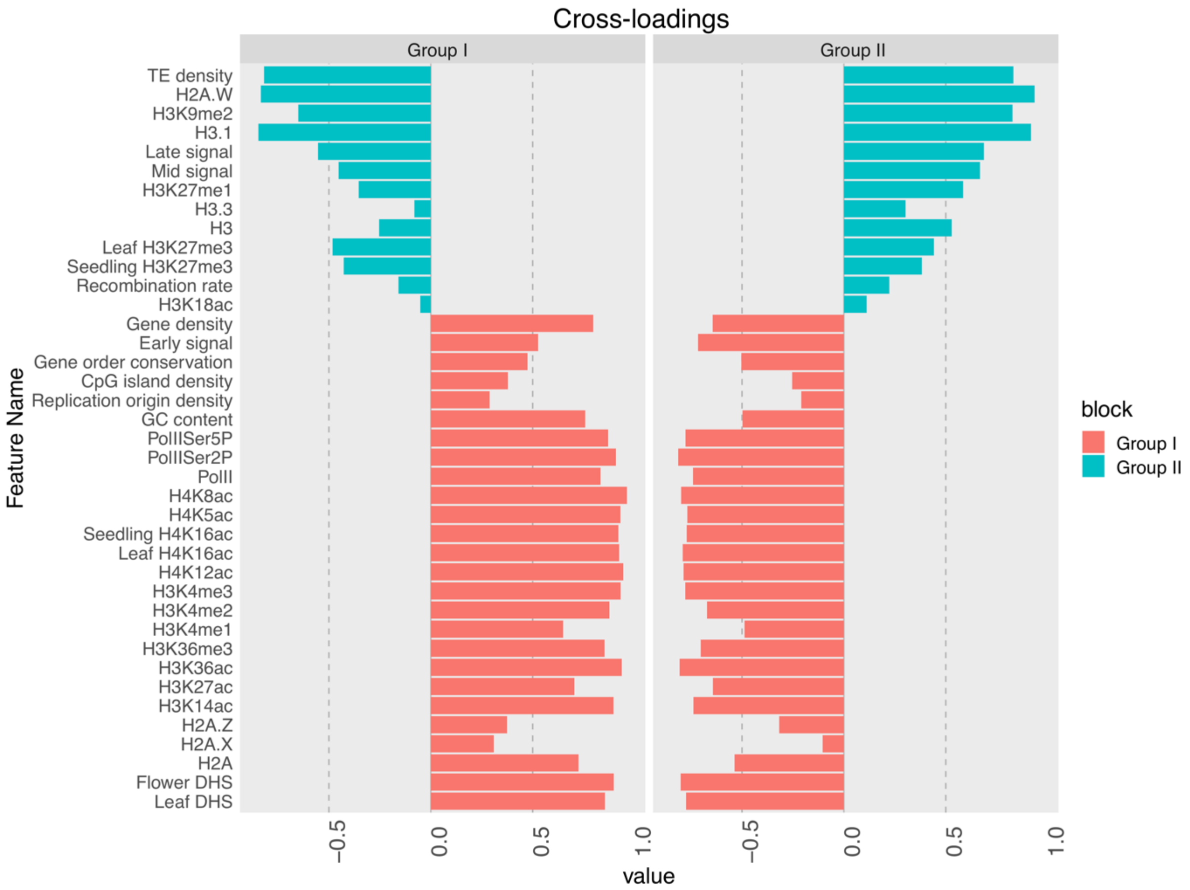

**Figure S10.** The loadings between the indicators and their own latent variables and the cross-loadings between the indicators and other latent variables. Red bars represent features that belong to group I and blue bars are features in group II. The $x$ axis gives the loadings and cross-loadings (i.e., correlations) between the observed indicators and the two latent variables.

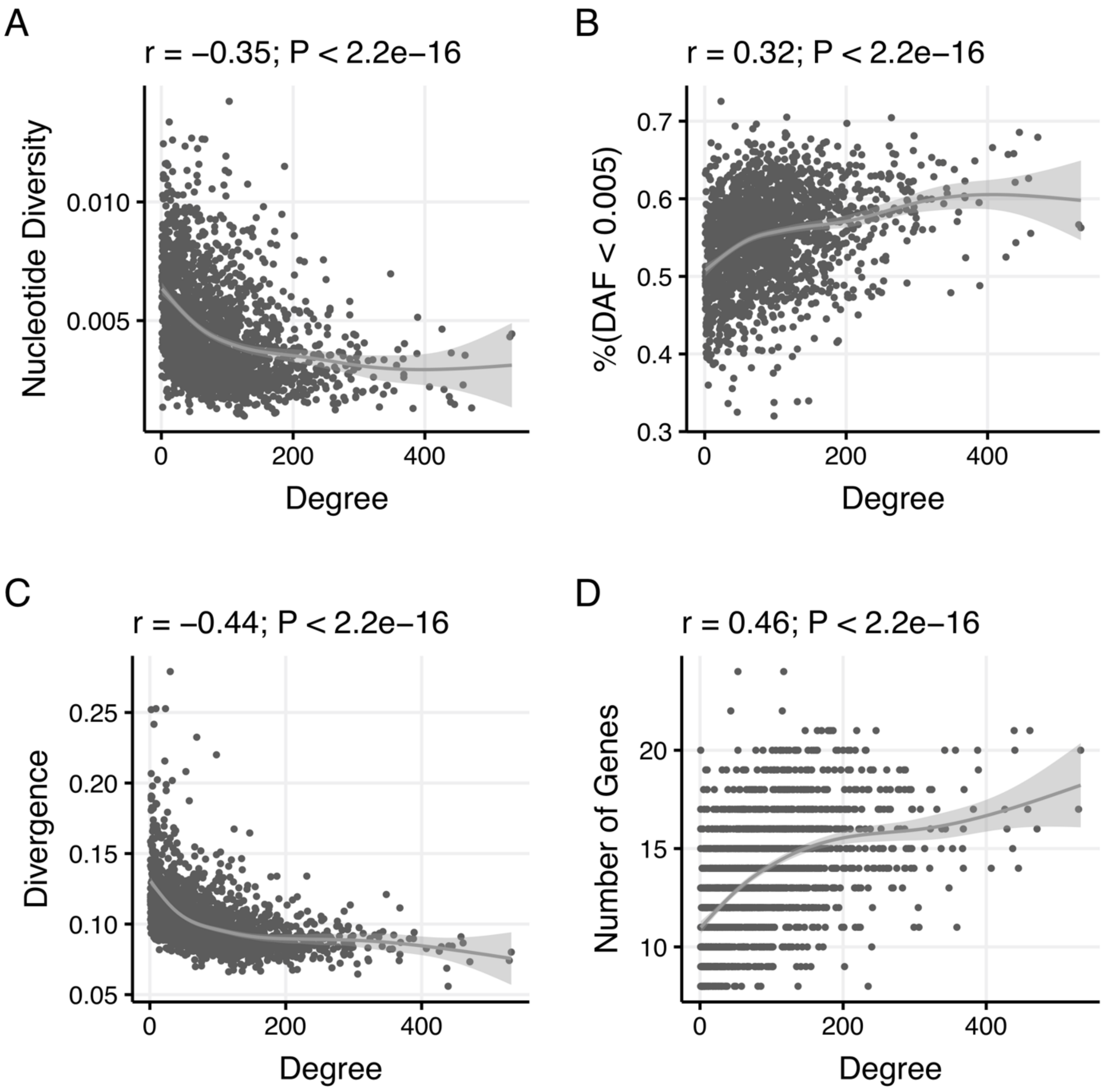

**Figure S11.** Correlations between node degree and evolutionary parameters at the 50 kb scale. (A) nucleotide diversity, (B) %(DAF<0.005), (C) evolutionary rates, and (D) the number of genes. The gray line and shaded area are the loess fitting line and its 95% confidence interval, respectively.

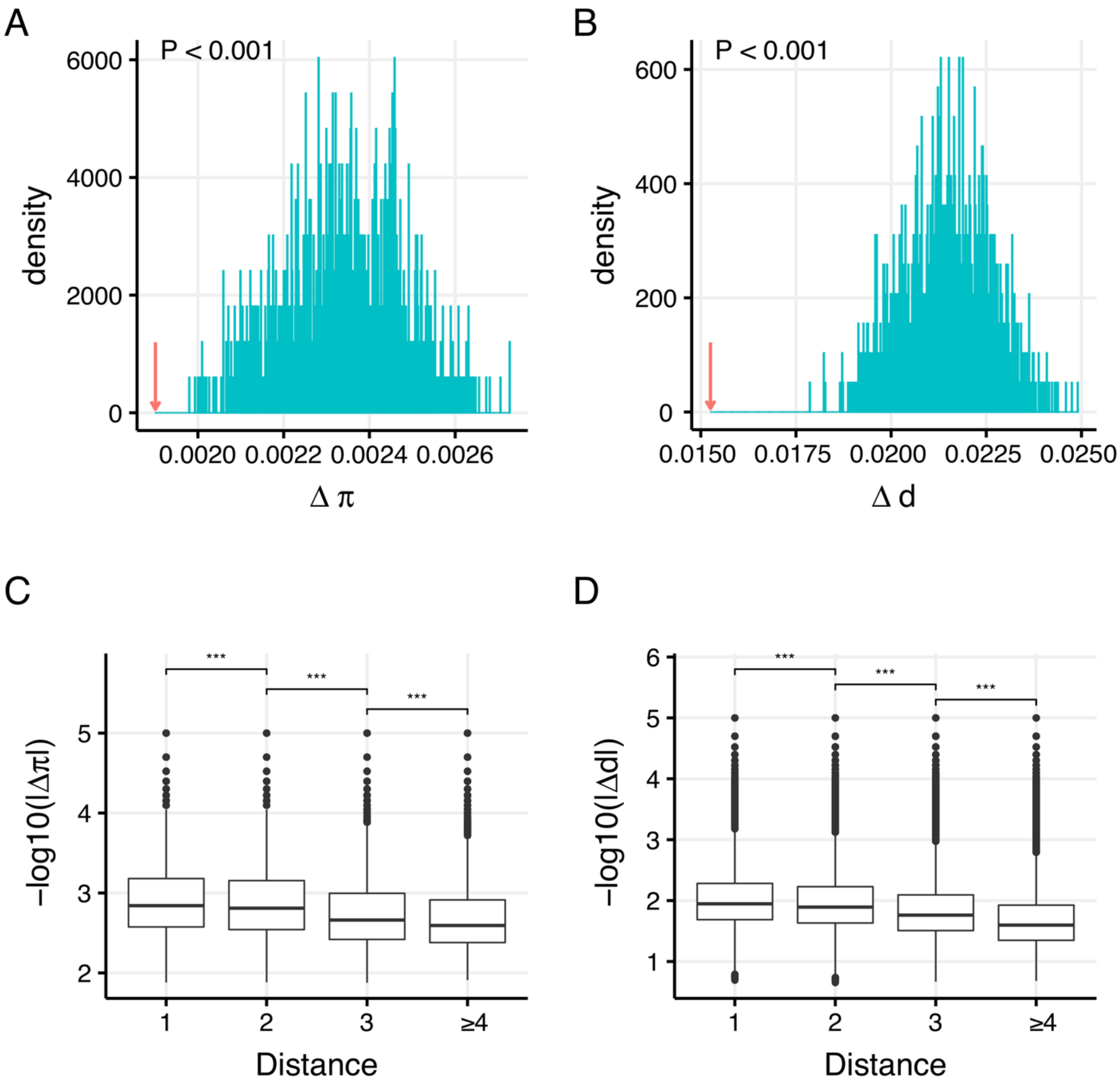

**Figure S12.** At the 50 kb scale, neighboring fragments in the CIN display similar genetic diversity and evolutionary rate. The upper two panels show the distribution of mean difference in genetic diversity (A) and evolutionary rates (B) between adjacent fragments of 1,000 simulated networks using the CCS null model. The blue lines show the simulated distribution of a parameter, while the red arrow represents the observed mean difference of a parameter. The lower two panels show that the similarity in genetic diversity (C) and evolutionary rates (D) between fragment pairs decreases when the 3D distance between them increases. *** $P < 0.001$.

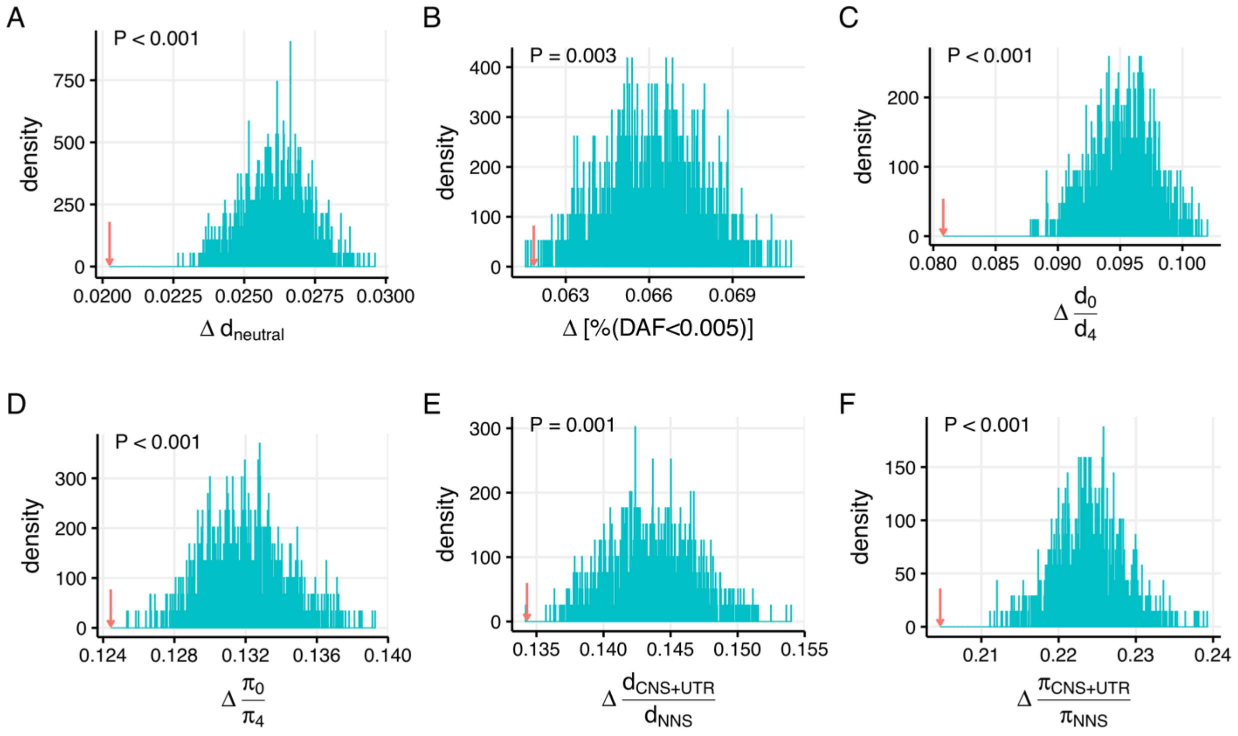

**Figure S13.** At the 50 kb scale, neighboring fragments in the CIN display similar regional mutation rates and evolutionary constraints. The observed average difference and the distribution of the average difference between adjacent nodes in simulated networks are shown for divergence at putatively neutral sites ($d_{neutral}$; A), $\%(DAF < 0.005)$ (B), $d_0/d_4$ (C), $\pi_0/\pi_4$ (D), $d_{CNS+UTR}/d_{NNS}$ (E), and $\pi_{CNS+UTR}/\pi_{NNS}$ (F).

A

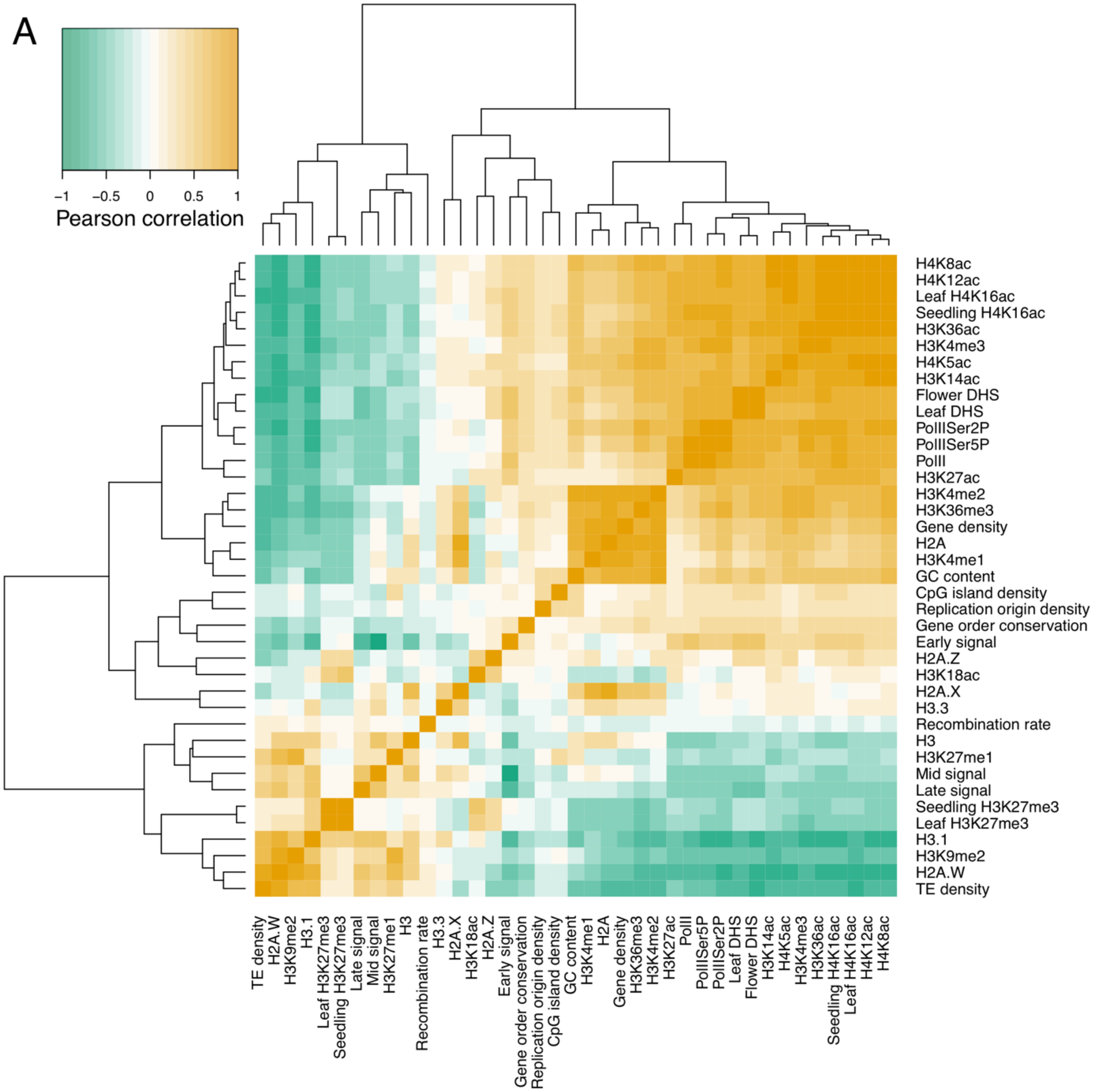

Pearson correlation
−1
−0.5
0
0.5
1
H4K8ac
H4K12ac
Leaf H4K16ac
Seedling H4K16ac
H3K36ac
H3K4me3
H4K5ac
H3K14ac
Flower DHS
Leaf DHS
PolIISer2P
PolIISer5P
PolII
H3K27ac
H3K4me2
H3K36me3
Gene density
H2A
H3K4me1
GC content
CpG island density
Replication origin density
Gene order conservation
Early signal
H2A.Z
H3K18ac
H2A.X
H3.3
Recombination rate
H3
H3K27me1
Mid signal
Late signal
Seedling H3K27me3
Leaf H3K27me3
H3.1
H3K9me2
H2A.W
TE density

B

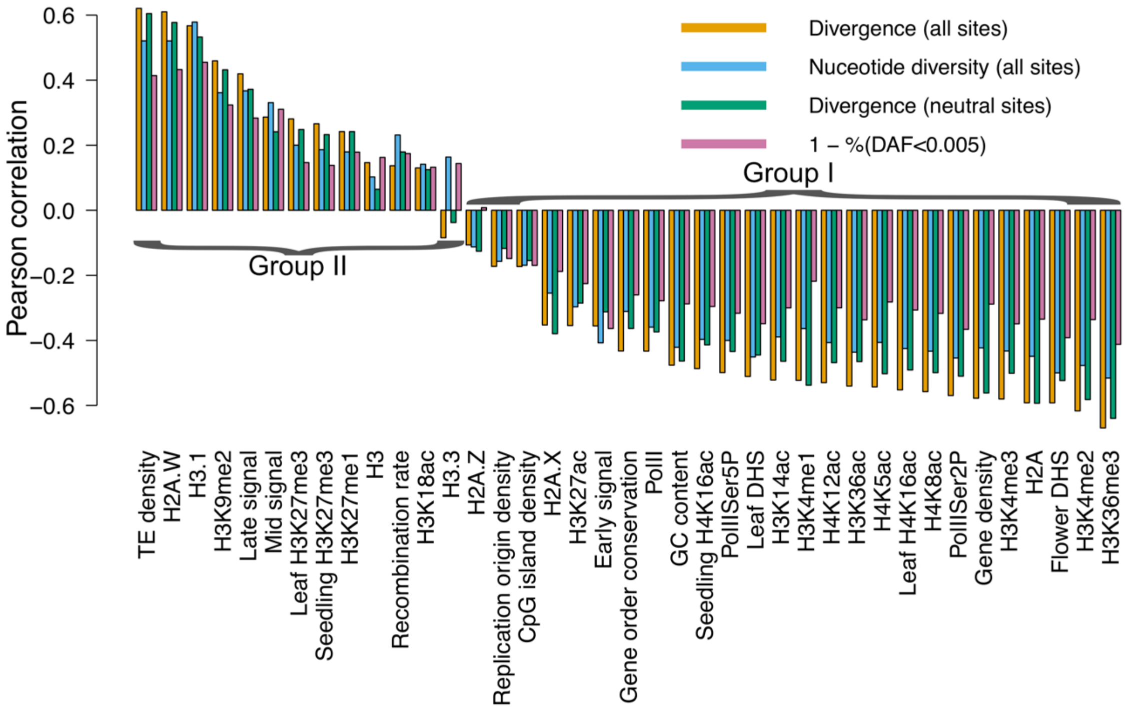
Divergence (all sites)
Nuceotide diversity (all sites)
Divergence (neutral sites)
1 – %(DAF<0.005)
Group I
Group II
Pearson correlation
0.6
0.4
0.2
0.0
−0.2
−0.4
−0.6
TE density
H2A.W
H3.1
H3K9me2
Late signal
Mid signal
Leaf H3K27me3
Seedling H3K27me3
H3K27me1
H3
Recombination rate
H3K18ac
H3.3
H2A.Z
Replication origin density
CpG island density
H2A.X
H3K27ac
Early signal
Gene order conservation
PolII
GC content
Seedling H4K16ac
PolIISer5P
Leaf DHS
H3K14ac
H3K4me1
H4K12ac
H3K36ac
H4K5ac
Leaf H4K16ac
H4K8ac
PolIISer2P
Gene density
H3K4me3
H2A
Flower DHS
H3K4me2
H3K36me3

C

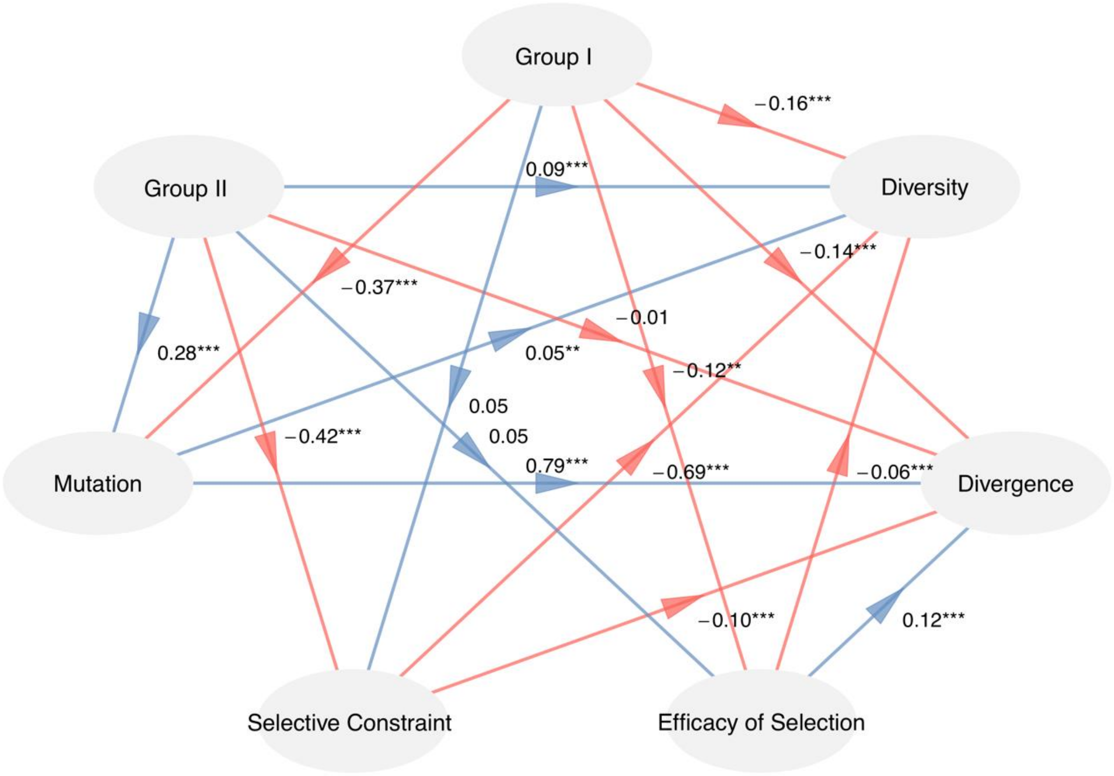
Group I
Group II
Diversity
Mutation
Divergence
Selective Constraint
Efficacy of Selection
−0.16***
0.09***
−0.14***
−0.37***
−0.01
0.28***
0.05**
−0.12**
0.05
−0.42***
0.05
0.79***
−0.69***
−0.06***
−0.10***
0.12***

**Figure S14.** At the 50 kb scale, a set of diverse (epi)genomic features affect local nucleotide diversity and evolutionary rates across the Arabidopsis genome. (A) Correlation heatmap of 39 (epi)genomic features. (B) Pearson's correlation coefficients of divergence, genetic diversity, divergence at putatively neutral sites and $1 - \%(DAF < 0.005)$ with diverse (epi)genomic features in 50 kb nonoverlapping windows. (C) The result of PLS-PM showing the causal relationship between two groups of features and evolutionary parameters. Divergence at putatively neutral sites, $\%(DAF < 0.005)$, and $\pi_0/\pi_4$ are used to represent mutation, selective constraint, and the efficacy of selection, respectively. Arrows represent the predefined causal directions between variables; red indicates a negative impact and blue a positive one. The path coefficients and their significance are labeled beside the arrows. *** P < 0.001, ** P < 0.01, and * P < 0.05.

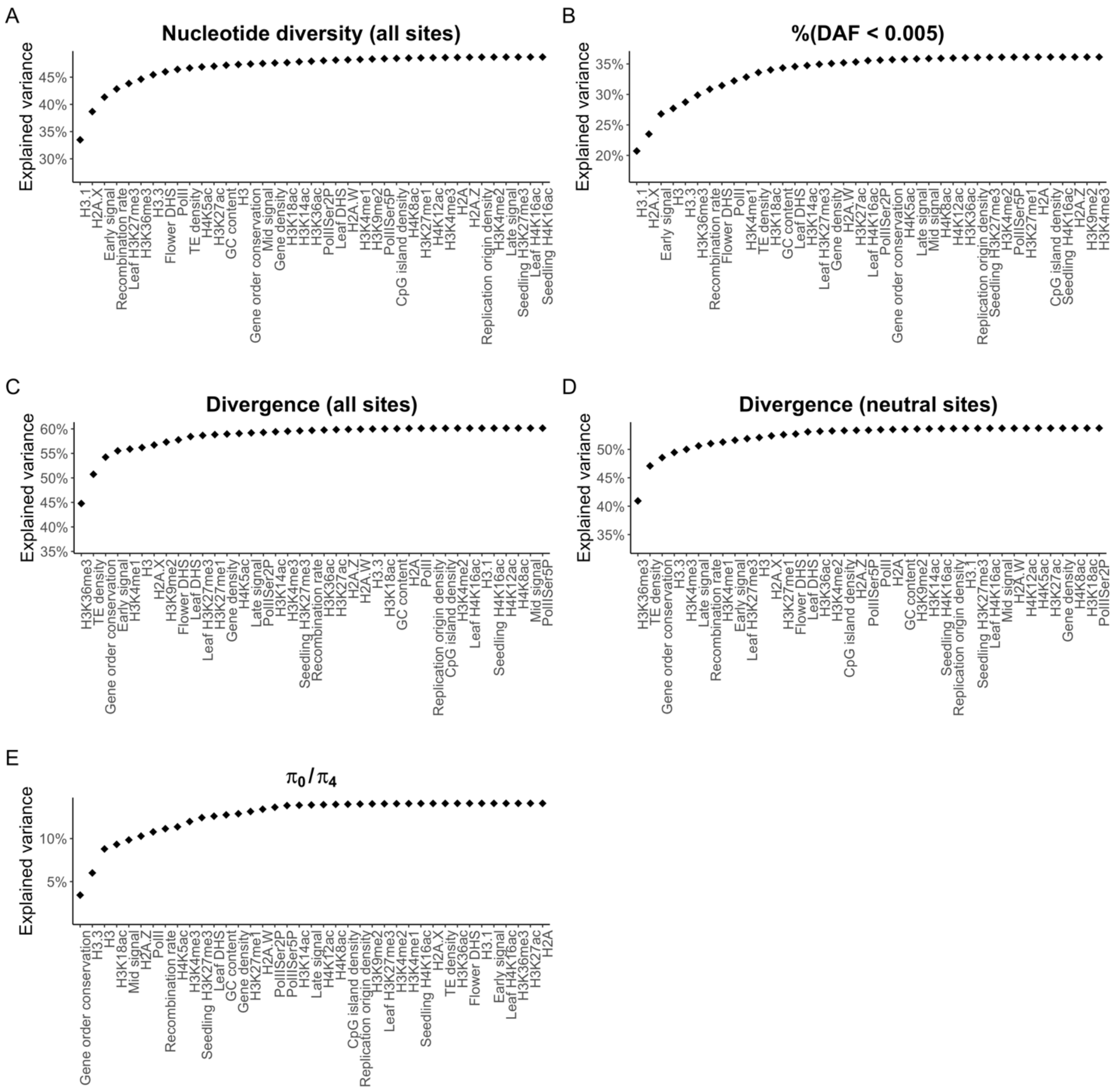

**Figure S15.** At the 50 kb scale, the combination of multiple (epi)genomic features can explain a great proportion of variation of genetic diversity and evolutionary rates across the Arabidopsis genome. The cumulative $R^2$ of linear models (adding the feature on the $x$ axis as a predictor at each step) for nucleotide diversity (A), $\%(DAF < 0.005)$ (B), divergence at all sites (C), divergence at neutral sites (D), and $\pi_0/\pi_4$ (E) are shown.

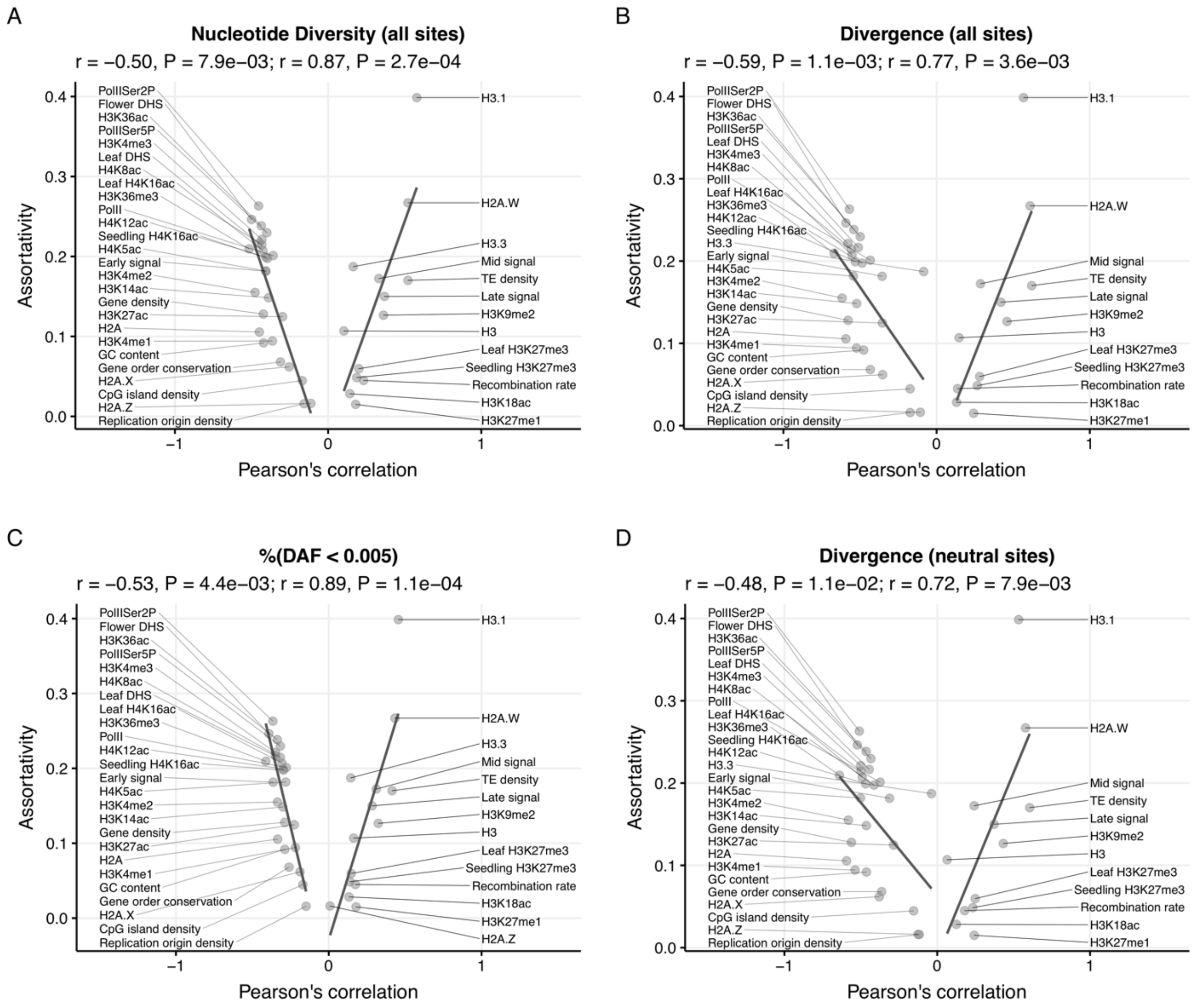

**Figure S16.** At the 50 kb scale, (epi)genomic features that have a higher impact on the evolution of regional sequences exhibit higher levels of assortativity in the CIN. The $x$ axis gives the Pearson's correlation coefficient of each genomic feature with genetic diversity (A), evolutionary rate (B), selective constraint (C), and mutation rate (D). The $y$ axis gives the assortativity of each feature in the CIN, which is the same in all panels. The solid lines represent the linear best fits between assortativity and the Pearson's correlation coefficients, and are obtained for r > 0 and r < 0 separately.

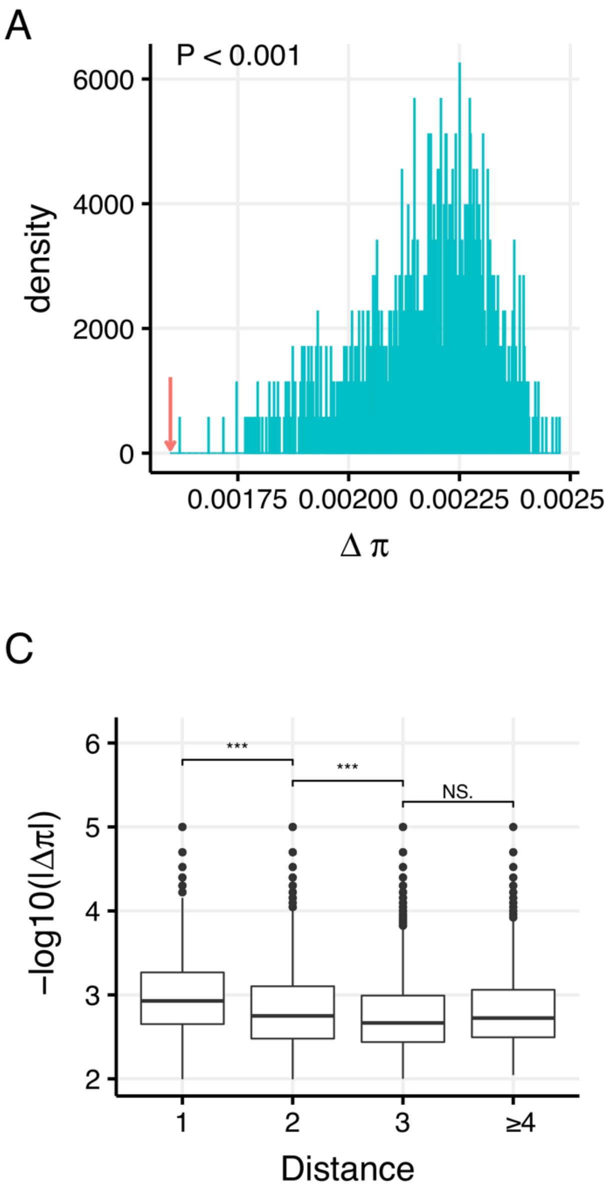

**Figure S17.** The use of different Hi-C dataset does not change the pattern of correlated evolution. In this case, the Hi-C data generated by Feng et al. (2014) was used to construct the CIN. The upper two panels show the distribution of mean difference in genetic diversity (A) and evolutionary rates (B) between adjacent fragments of 1,000 simulated networks using CCS model. The blue lines show the simulated distribution of a parameter, while the red arrow represents the observed mean difference of a parameter. The lower two panels show that the similarity of fragment pairs decreases when the 3D distance between them increases in terms of genetic diversity (C) and evolutionary rates (D). *** $P < 0.001$, NS: nonsignificant.

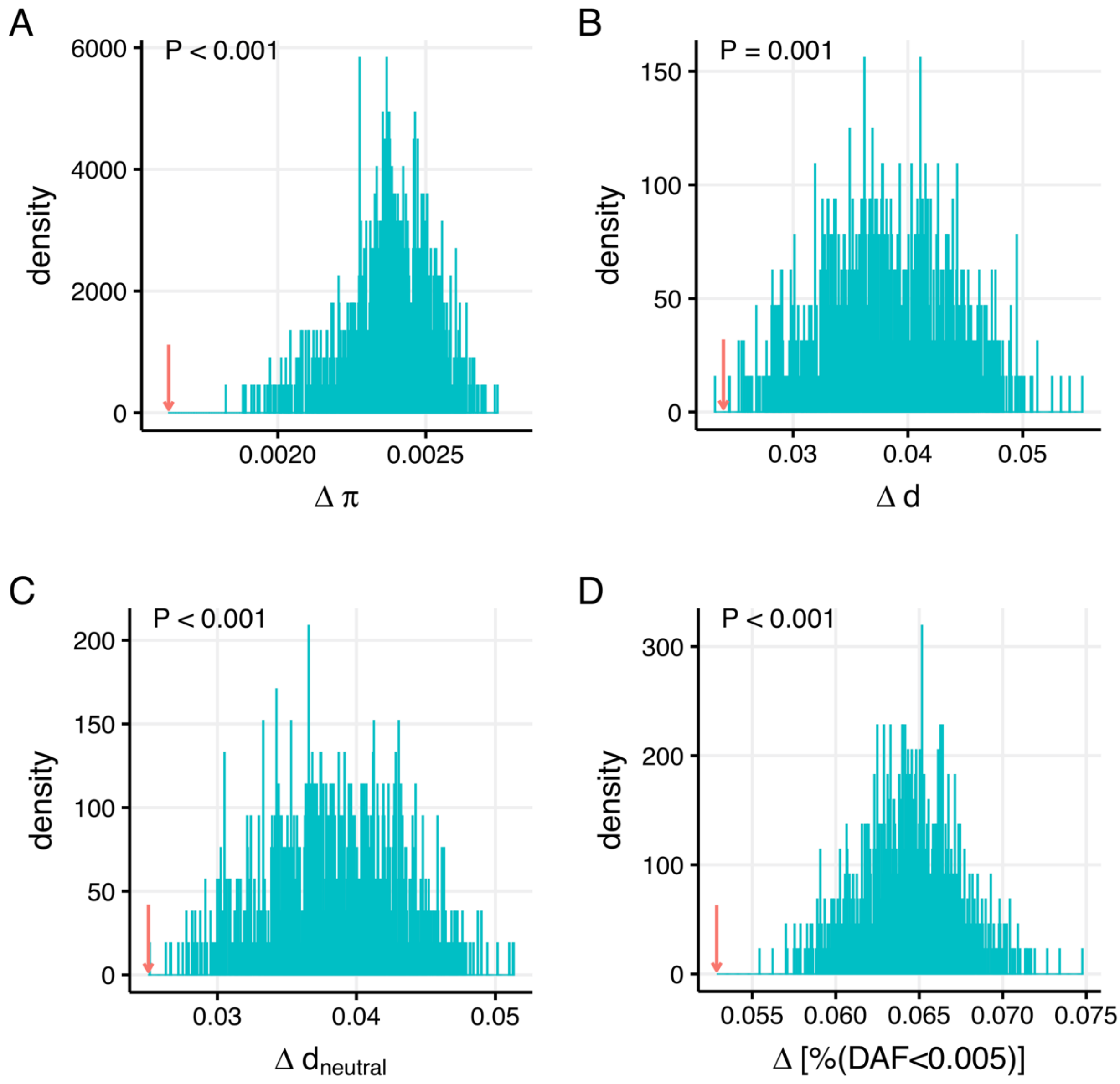

**Figure S18.** Distribution of difference in evolutionary parameters between interacting fragments using the CGS null model when no filtration is applied regarding to gene number and mappability in a fragment. Red arrows show the observed differences between contacting fragments in the real data. (A) Nucleotide diversity; (B) Divergence between *A. thaliana* and *A. lyrata*; (C) Divergence at putatively neutral sites; (D) $\%(DAF < 0.005)$.

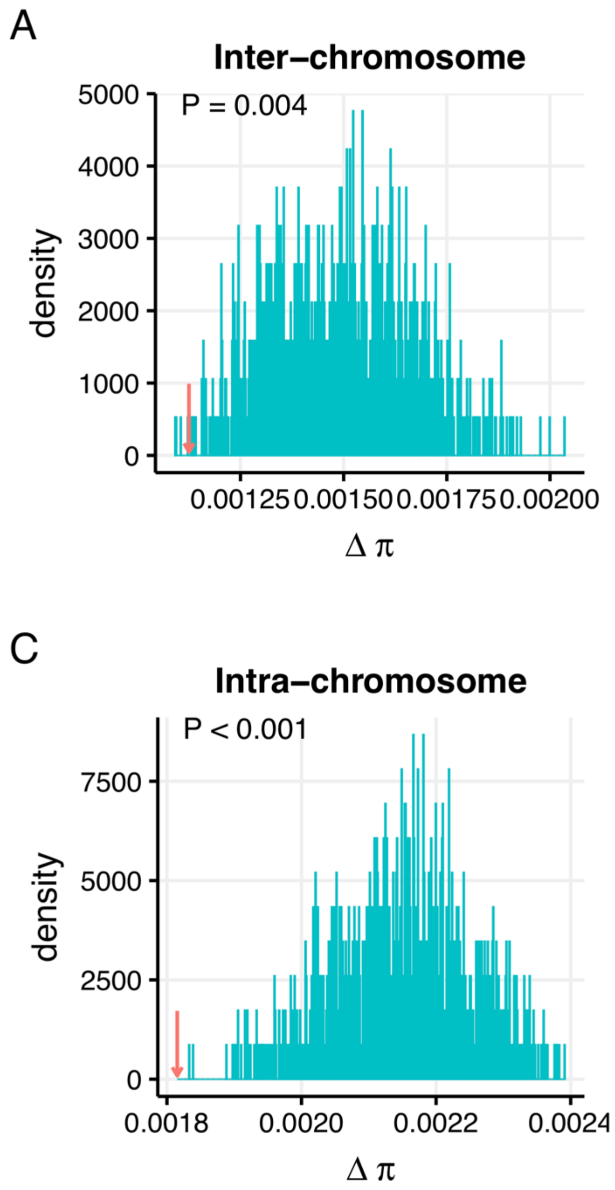

**Figure S19.** Distribution of difference in evolutionary parameters between interacting fragments using the CGS null model when only inter- or intra-chromosomal interactions are considered. Red arrows show the observed differences between contacting fragments in the real data. (A) and (B) show the results of simulations of nucleotide diversity and divergence for inter-chromosomal interactions; (C) and (D) show the results of simulations of nucleotide diversity and divergence for intra-chromosomal interactions.

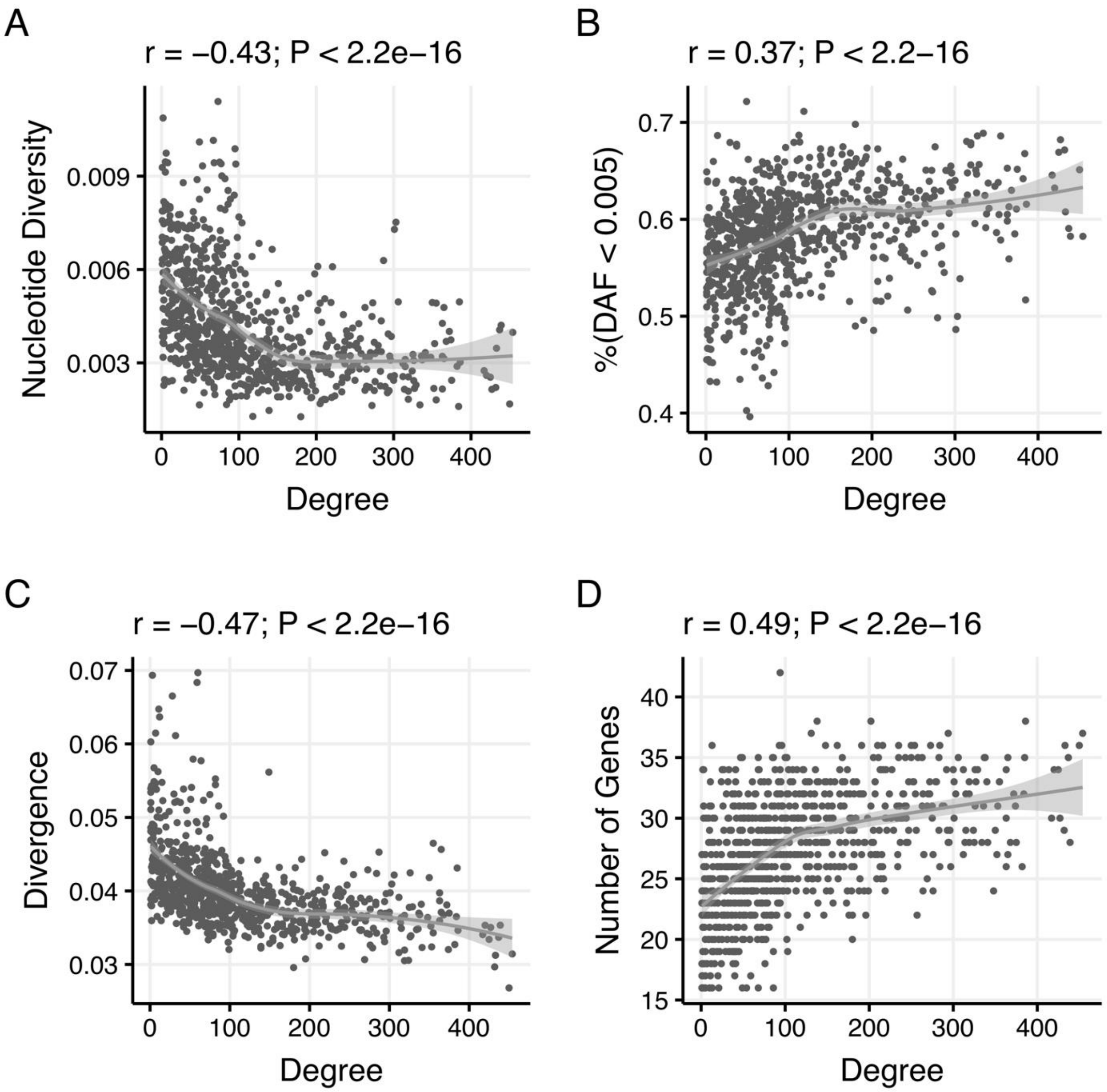

**Figure S20.** Correlations between node degree and evolutionary parameters when using *A. thaliana* lineage specific divergence as a measure of evolutionary rate. (A) nucleotide diversity, (B) %(DAF<0.005), (C) evolutionary rates, and (D) the number of genes. The gray line and shaded area are the loess fitting line and its 95% confidence interval, respectively.

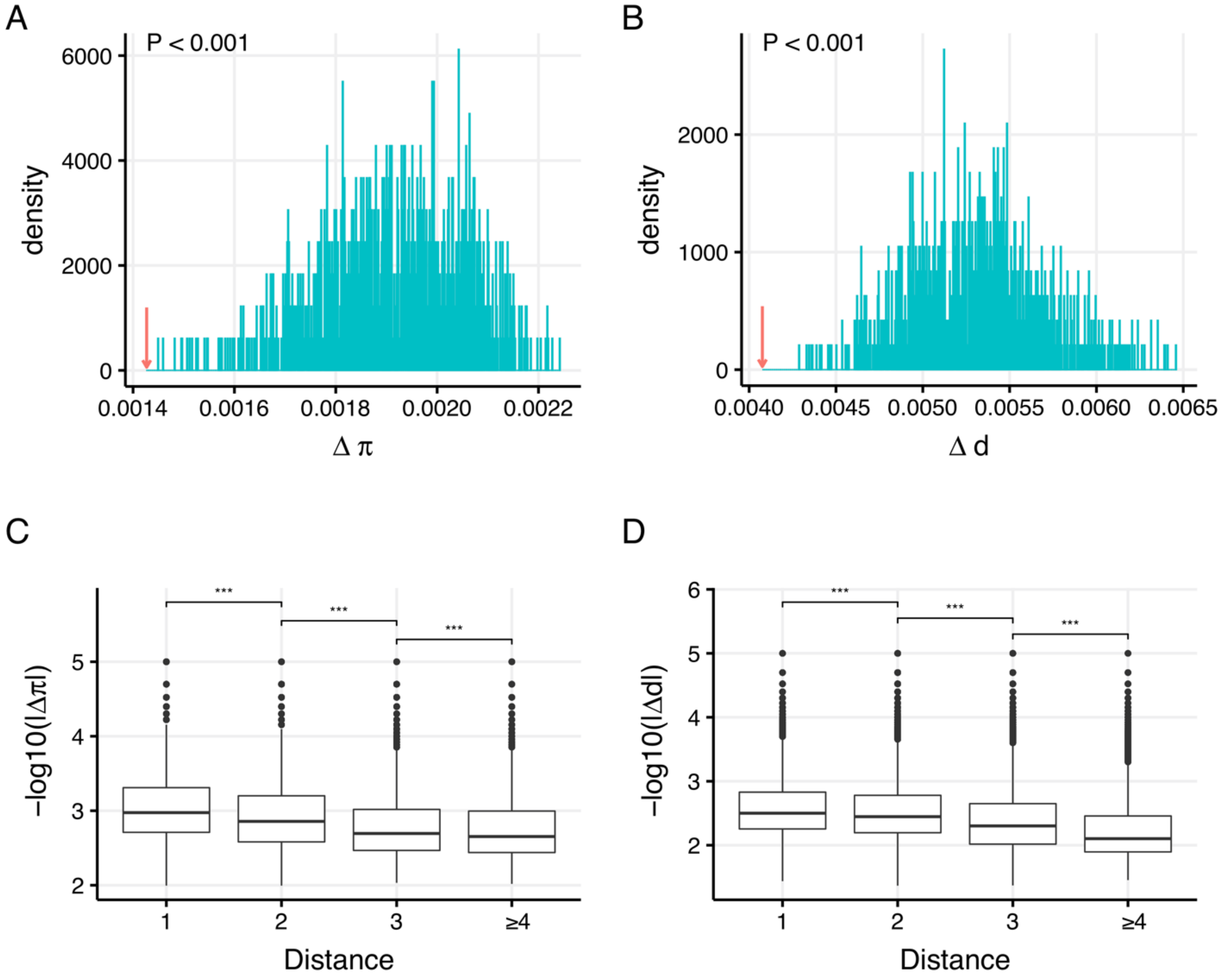

**Figure S21.** When using *A. thaliana* lineage specific divergence as a measure of evolutionary rate, neighboring fragments in the CIN display similar genetic diversity and evolutionary rate. The upper two panels show the distribution of mean difference in genetic diversity (A) and evolutionary rates (B) between adjacent fragments of 1,000 simulated networks using the CCS null model. The blue lines show the simulated distribution of a parameter, while the red arrow represents the observed mean difference of a parameter. The lower two panels show that the similarity in genetic diversity (C) and evolutionary rates (D) between fragment pairs decreases when the 3D distance between them increases. *** $P < 0.001$.

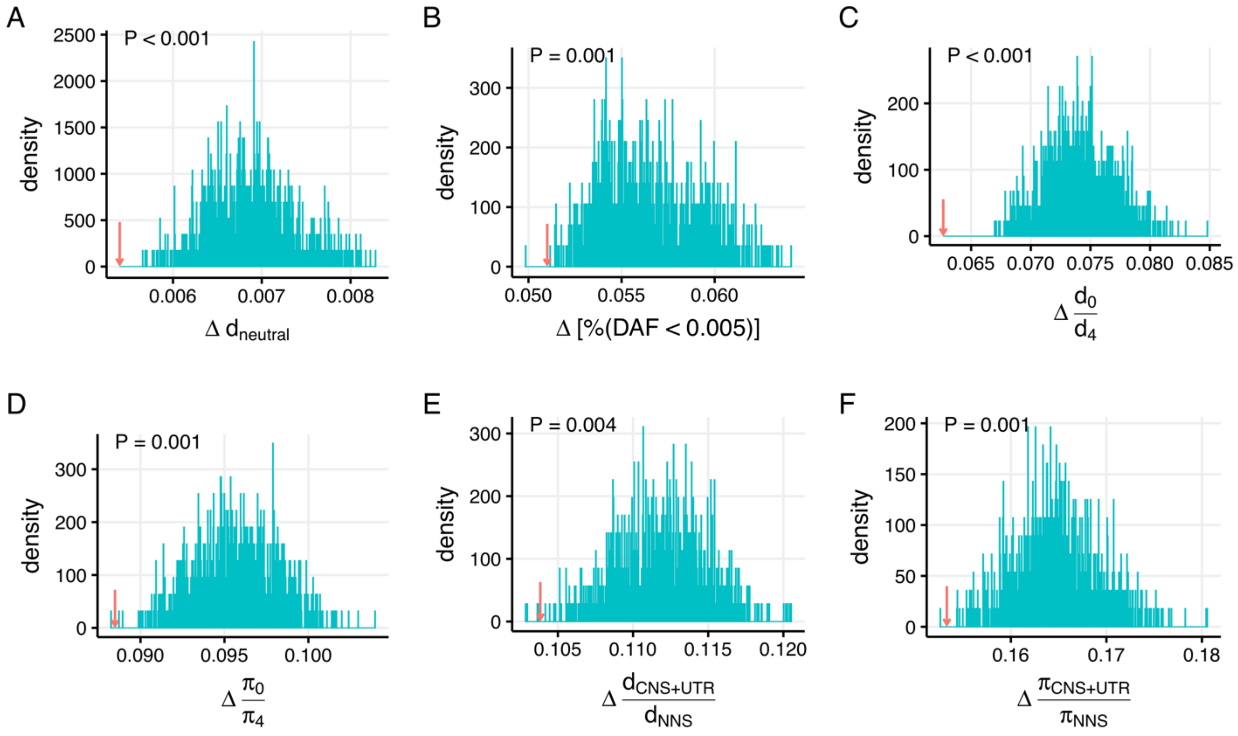

**Figure S22.** When using *A. thaliana* lineage specific divergence as a measure of evolutionary rate, neighboring fragments in the CIN exhibit similar regional mutation rates and evolutionary constraints. The observed average difference and the distribution of the average difference between adjacent nodes in simulated networks are shown for divergence at putatively neutral sites ($d_{neutral}$; A), $\%(DAF < 0.005)$ (B), $d_0/d_4$ (C), $\pi_0/\pi_4$ (D), $d_{CNS+UTR}/d_{NNS}$ (E), and $\pi_{CNS+UTR}/\pi_{NNS}$ (F).

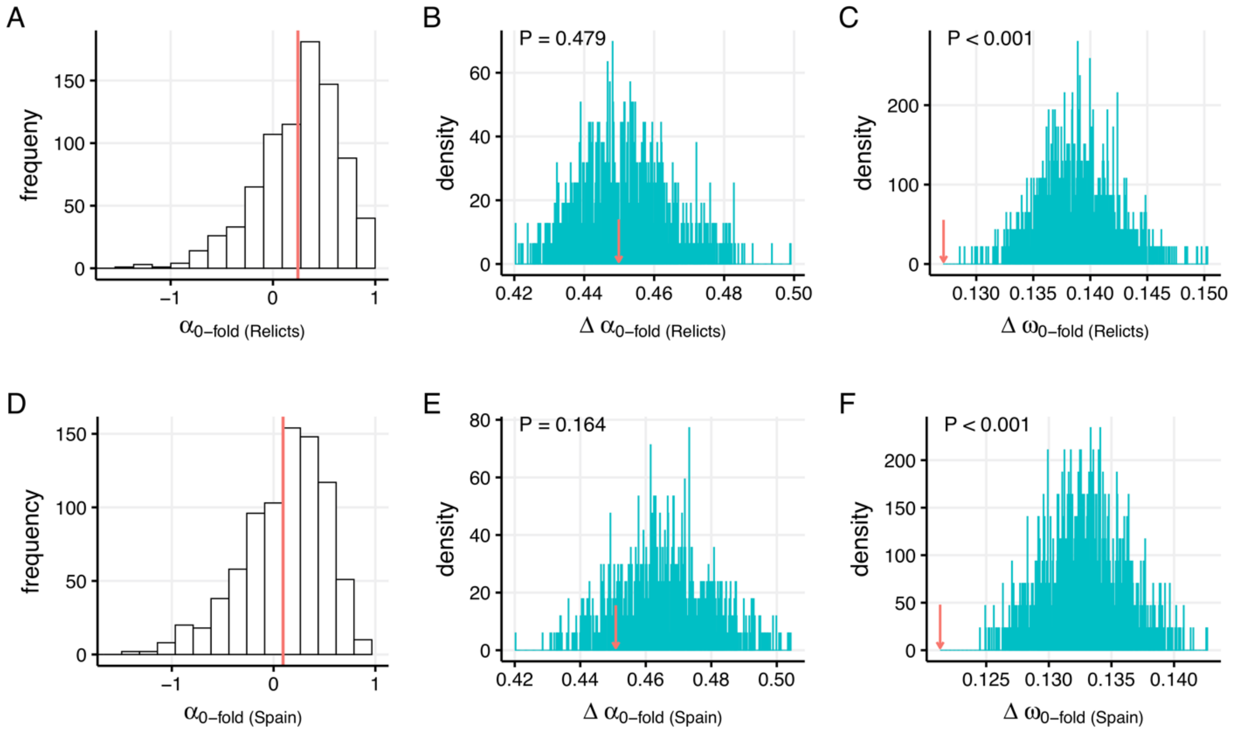

**Figure S23.** The rate of adaptive evolution and similarity of positive selection between 3D neighboring fragments when using *A. thaliana* lineage specific divergence as a measure of evolutionary rate. The upper three panels show the distribution of $\alpha$ (A) in 100 kb fragments and the results of CCS simulations for $\alpha$ (B) and $\omega_a$ (C) using polymorphism data from the relict population. The lower three panels show the distribution of $\alpha$ (A) in 100 kb fragments and the results of CCS simulations for $\alpha$ (B) and $\omega_a$ (C) using polymorphism data from the Spanish population.

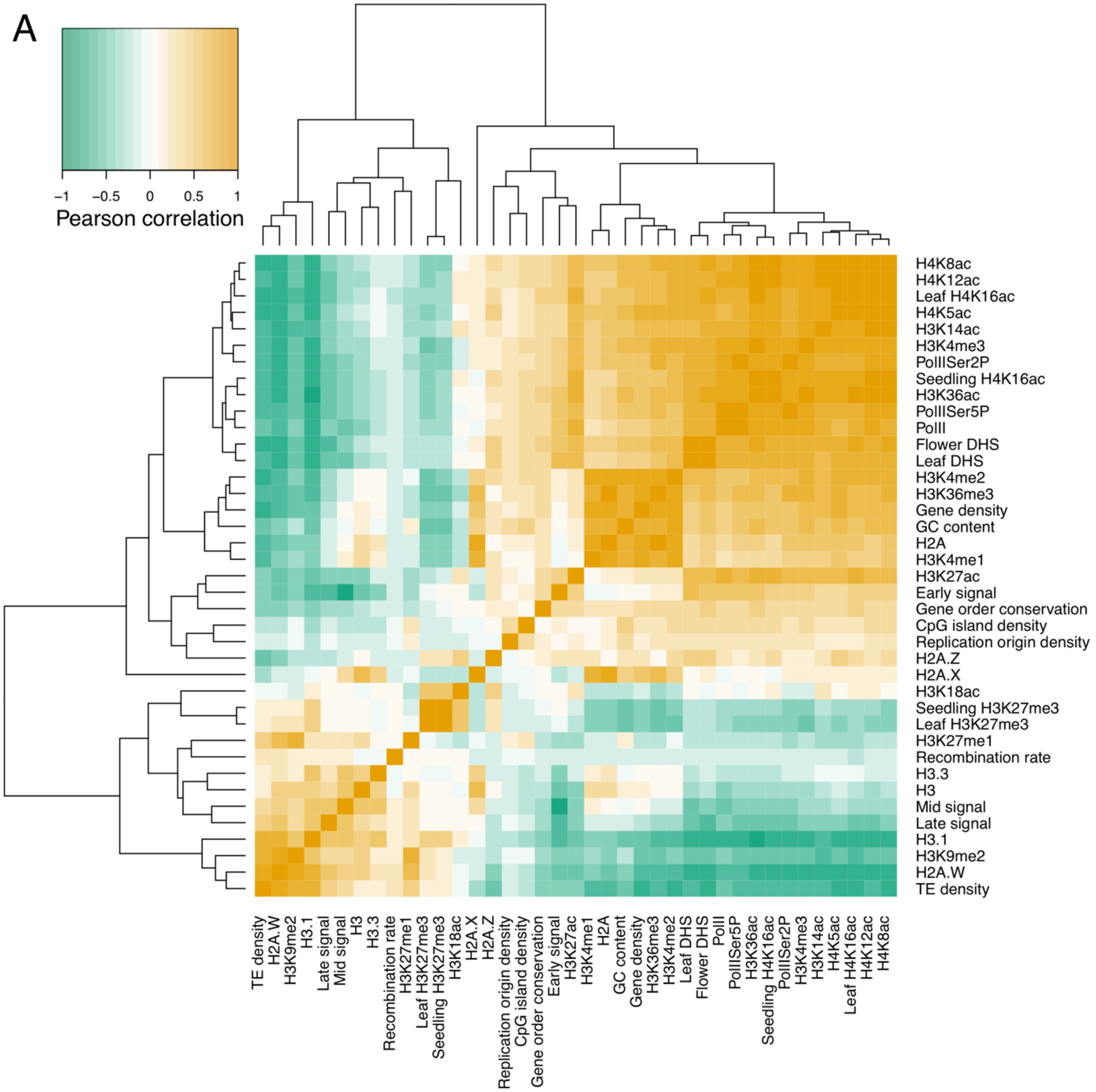
A
Pearson correlation
−1
−0.5
0
0.5
1
H4K8ac
H4K12ac
Leaf H4K16ac
H4K5ac
H3K14ac
H3K4me3
PolIISer2P
Seedling H4K16ac
H3K36ac
PolIISer5P
PolII
Flower DHS
Leaf DHS
H3K4me2
H3K36me3
Gene density
GC content
H2A
H3K4me1
H3K27ac
Early signal
Gene order conservation
CpG island density
Replication origin density
H2A.Z
H2A.X
H3K18ac
Seedling H3K27me3
Leaf H3K27me3
H3K27me1
Recombination rate
H3.3
H3
Mid signal
Late signal
H3.1
H3K9me2
H2A.W
TE density

B

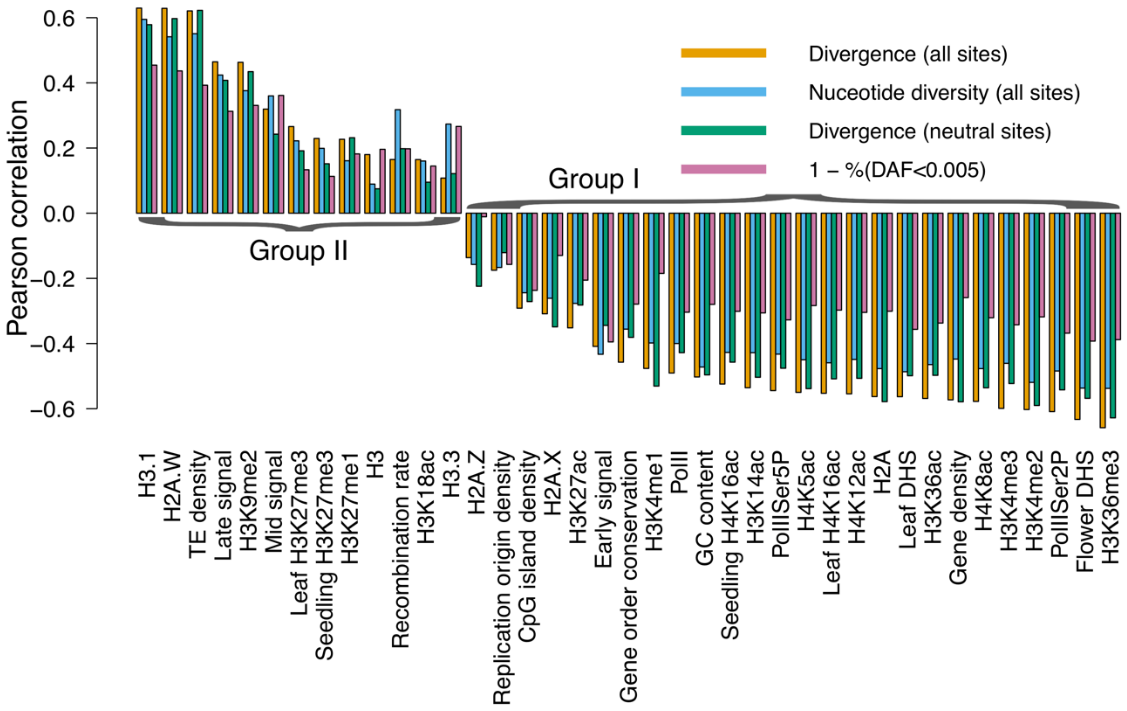
Pearson correlation
0.6
0.4
0.2
0.0
−0.2
−0.4
−0.6
Divergence (all sites)
Nuceotide diversity (all sites)
Divergence (neutral sites)
1 − %(DAF<0.005)
Group I
Group II
H3.1
H2A.W
TE density
Late signal
H3K9me2
Mid signal
Leaf H3K27me3
Seedling H3K27me3
H3K27me1
H3
Recombination rate
H3K18ac
H3.3
H2A.Z
Replication origin density
CpG island density
H2A.X
H3K27ac
Early signal
Gene order conservation
H3K4me1
PolII
GC content
Seedling H4K16ac
H3K14ac
PolIISer5P
H4K5ac
Leaf H4K16ac
H4K12ac
H2A
Leaf DHS
H3K36ac
Gene density
H4K8ac
H3K4me3
H3K4me2
PolIISer2P
Flower DHS
H3K36me3

C

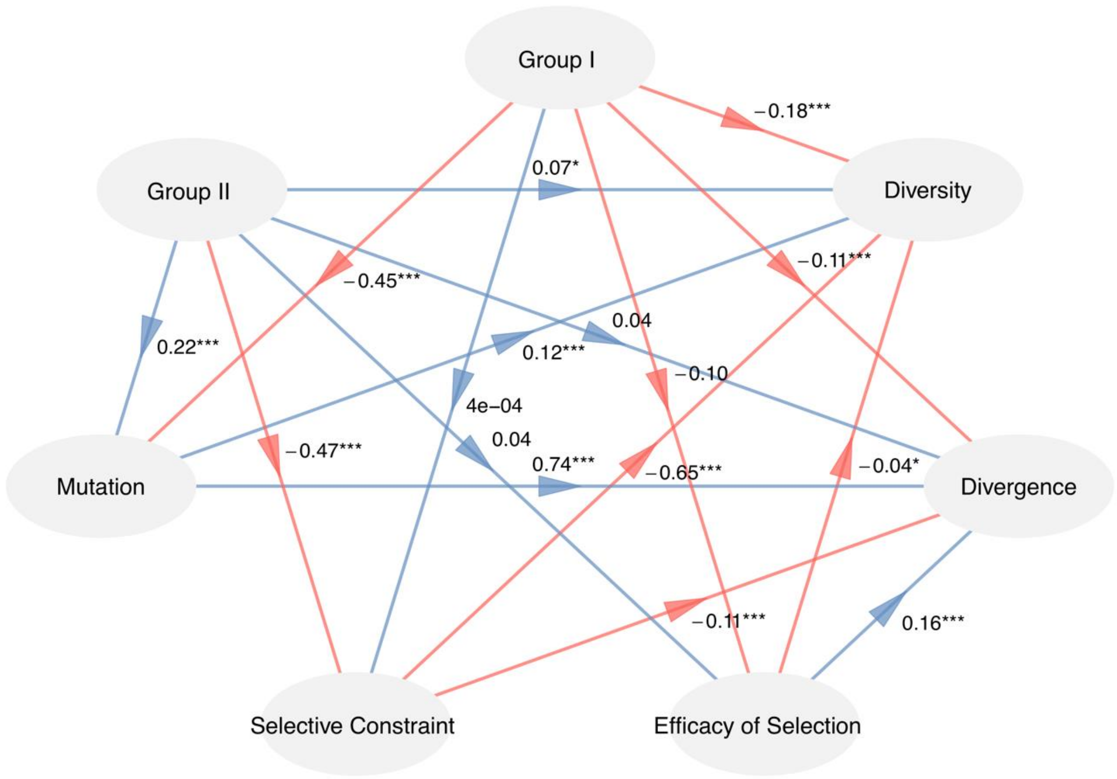
Group I
Group II
Diversity
Mutation
Divergence
Selective Constraint
Efficacy of Selection
−0.18***
0.07*
−0.11***
−0.45***
0.22***
0.04
0.12***
−0.10
4e−04
0.04
−0.47***
0.74***
−0.65***
−0.04*
−0.11***
0.16***

**Figure S24.** When using *A. thaliana* lineage specific divergence as a measure of evolutionary rate, a set of diverse (epi)genomic features affect local nucleotide diversity and evolutionary rates across the genome. (A) Correlation heatmap of 39 (epi)genomic features. (B) Pearson's correlation coefficients of divergence, genetic diversity, divergence at putatively neutral sites and $1 - \%(DAF < 0.005)$ with diverse (epi)genomic features in 100 kb nonoverlapping windows. (C) The result of PLS-PM showing the causal relationship between two groups of features and evolutionary parameters. Divergence at putatively neutral sites, $\%(DAF < 0.005)$, and $\pi_0/\pi_4$ are used to represent mutation, selective constraint, and the efficacy of selection, respectively. Arrows represent the predefined causal directions between variables; red indicates a negative impact and blue a positive one. The path coefficients and their significance are labeled beside the arrows. *** P < 0.001, ** P < 0.01, and * P < 0.05.

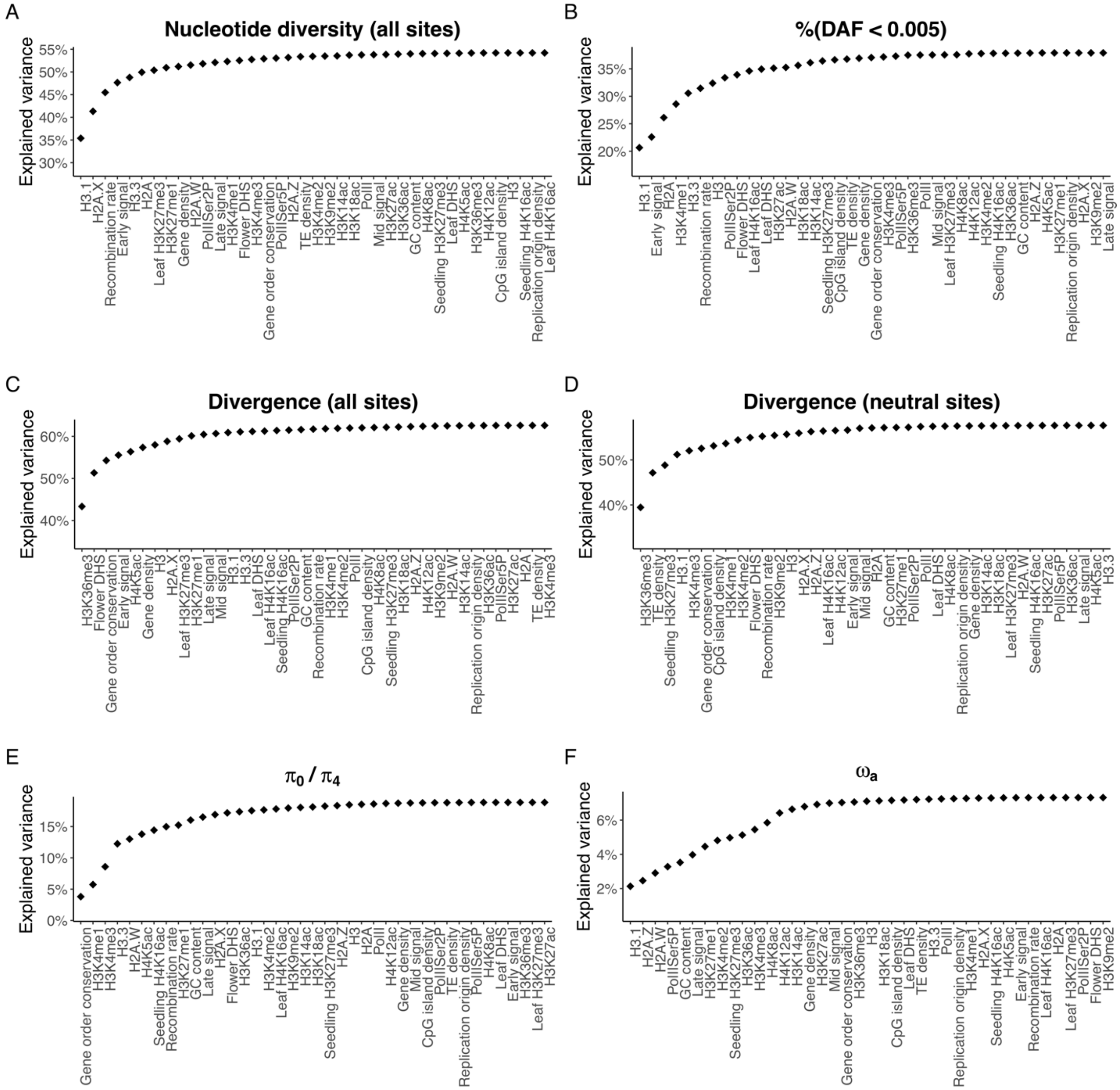

**Figure S25.** When using *A. thaliana* lineage specific divergence as a measure of evolutionary rate, the combination of multiple (epi)genomic features can explain a great proportion of variation of genetic diversity and evolutionary rates across the Arabidopsis genome. The cumulative $R^2$ of linear models (adding the feature on the $x$ axis as a predictor at each step) for nucleotide diversity (A), $\%(DAF < 0.005)$ (B), divergence at all sites (C), divergence at neutral sites (D), $\pi_0/\pi_4$ (E), and $\omega_a$ (F) are shown.

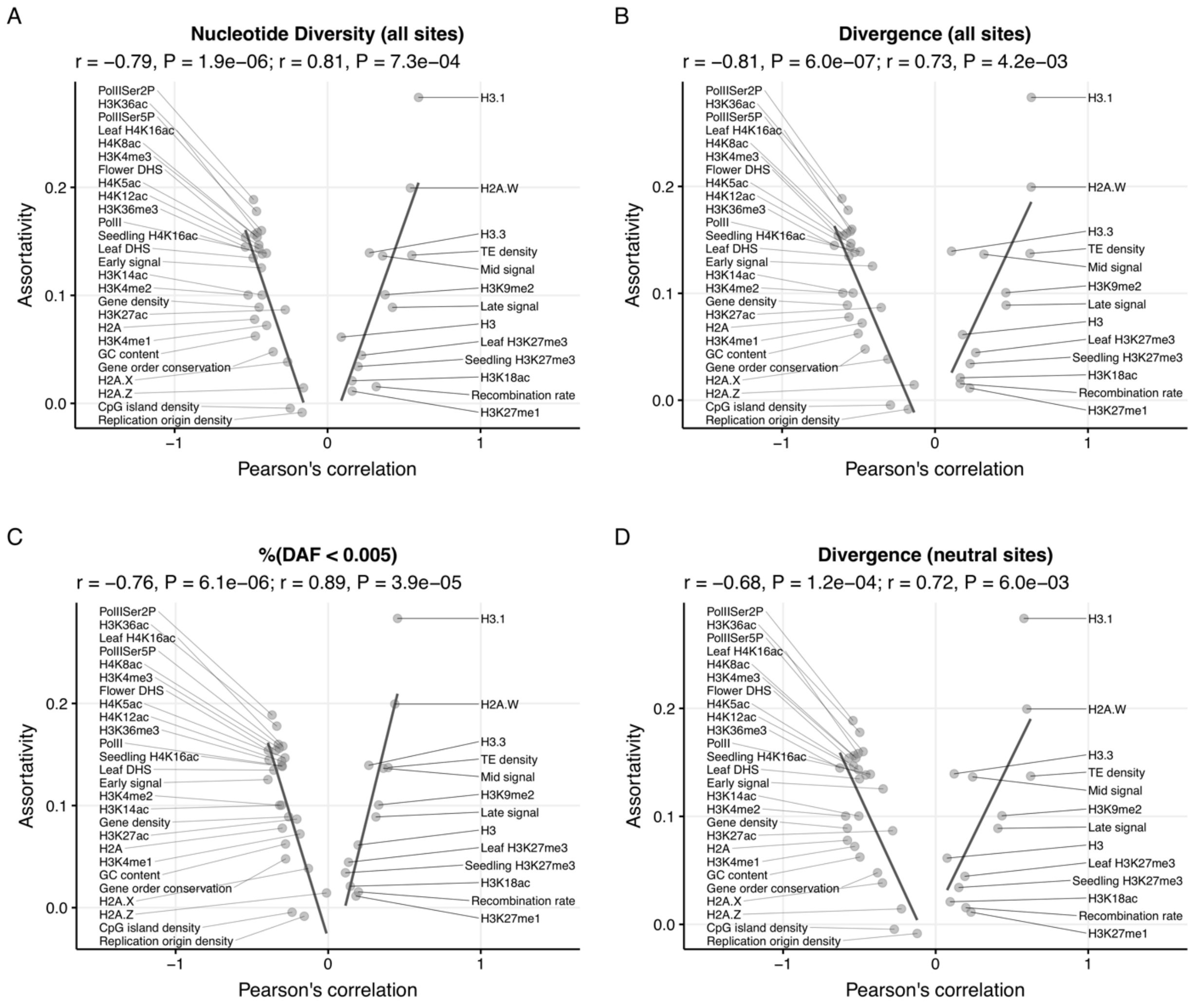

**Figure S26.** When using *A. thaliana* lineage specific divergence as a measure of evolutionary rate, (epi)genomic features that have a higher impact on the evolution of regional sequences exhibit higher levels of assortativity in the CIN. The $x$ axis gives the Pearson's correlation coefficient of each genomic feature with genetic diversity (A), evolutionary rate (B), selective constraint (C), and mutation rate (D). The $y$ axis gives the assortativity of each feature in the CIN, which is the same in all panels. The solid lines represent the linear best fits between assortativity and the Pearson's correlation coefficients, and are obtained for r > 0 and r < 0 separately.

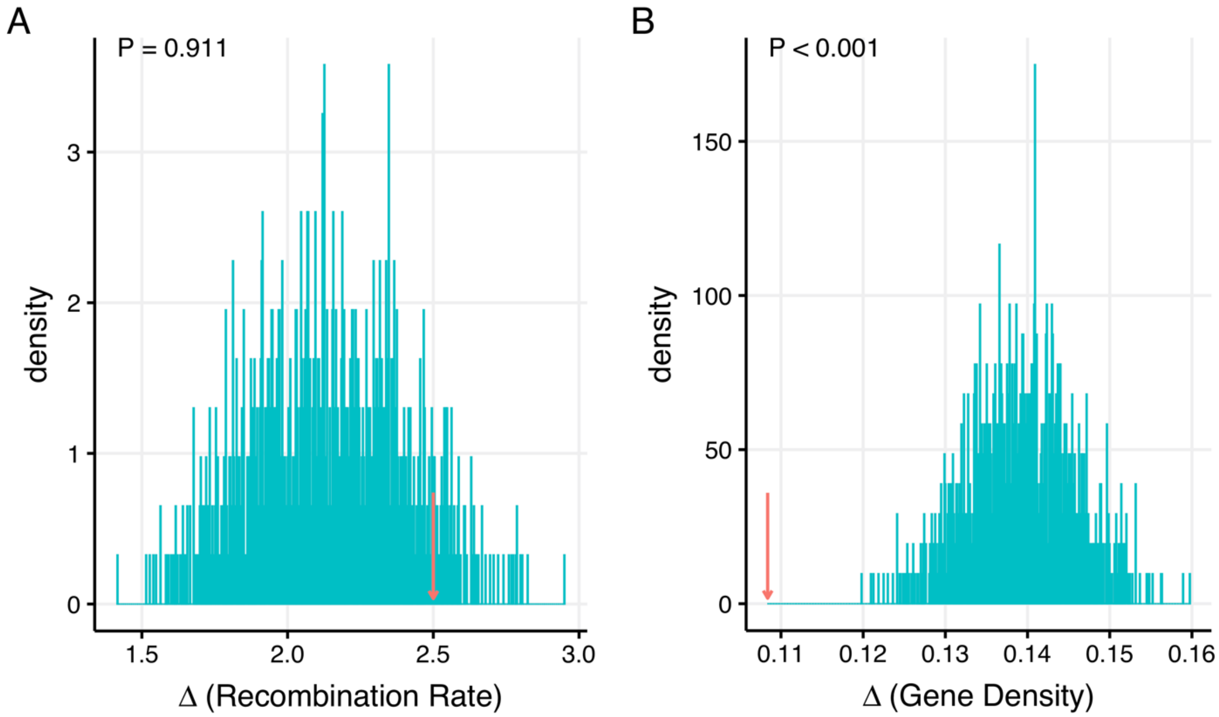

**Figure S27.** Distribution of difference in local recombination rates and gene density between interacting fragments using the CCS null model. The blue lines show the simulated distribution of a parameter, while the red arrow represents the observed mean difference of a parameter. (A) Recombination rate; (B) Gene density.

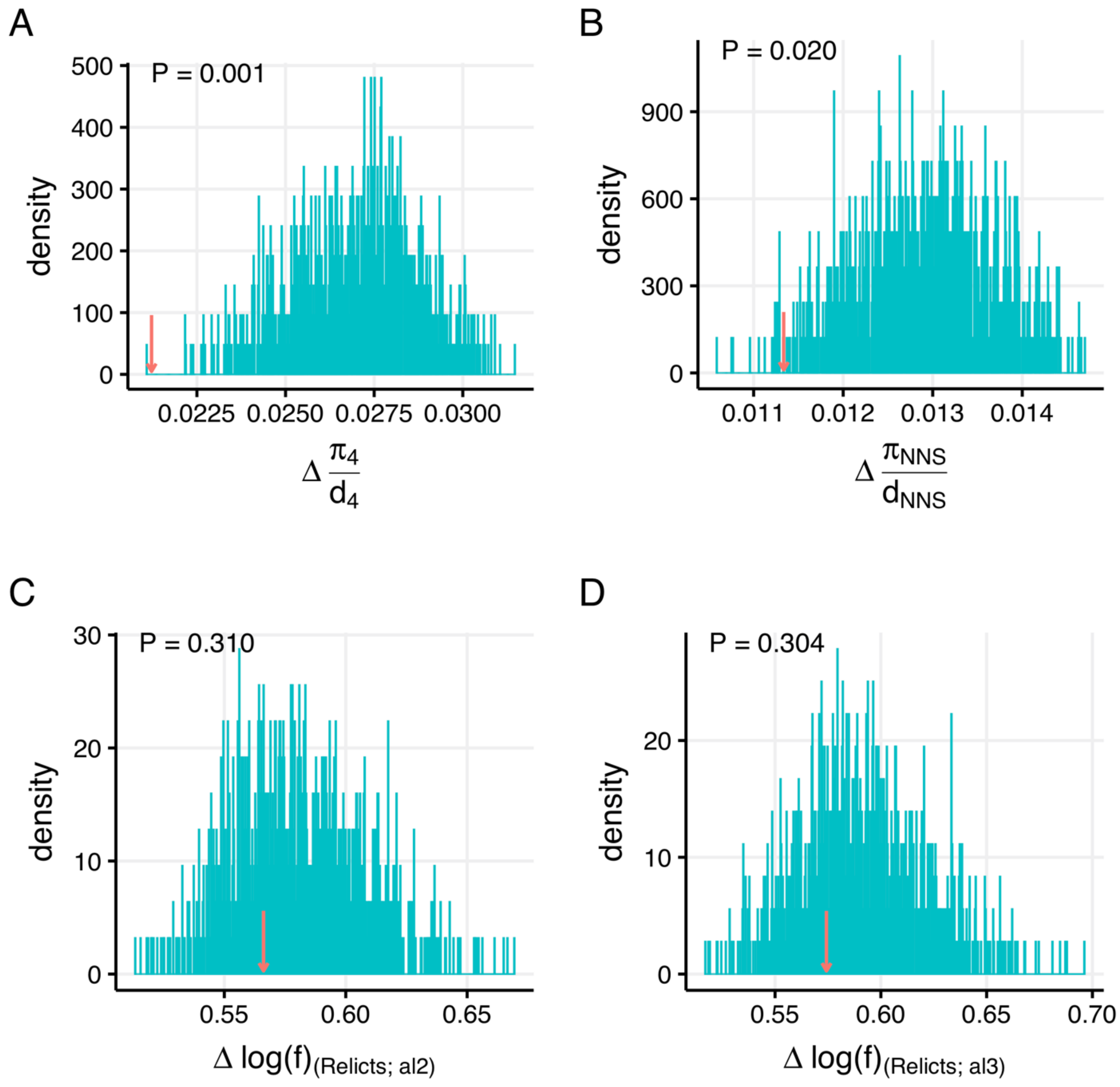

**Figure S28.** Distribution of difference in $N_e$ between interacting fragments using the CCS null model. The blue lines show the simulated distribution of a parameter, while the red arrow represents the observed mean difference of a parameter. (A) $N_e$ measured as $\pi_4/d_4$; (B) $N_e$ measured as $\pi_{NNS}/d_{NNS}$; (C) Relative $N_e$ [denoted as $\log(f)$] estimated using varne with the LBFGS algorithm; (D) Relative $N_e$ estimated using varne with the TNEWTON algorithm.

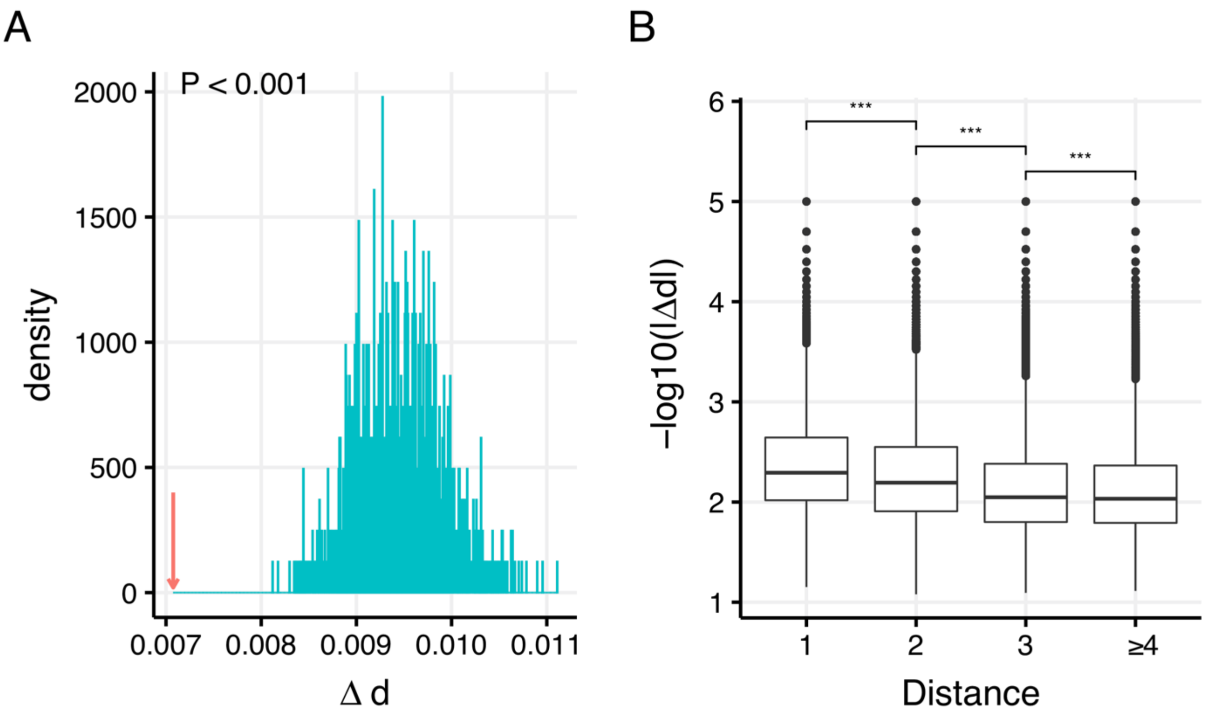

**Figure S29.** Correlated evolution between 3D interacting fragments in the *A. lyrata* genome at the 100 kb scale. (A) The distribution of mean difference in evolutionary rates between adjacent fragments of 1,000 simulated networks using CCS model. The blue lines show the simulated distribution of a parameter, while the red arrow represents the observed mean difference of a parameter. (B) shows that the similarity in evolutionary rates between fragments decreases when the 3D distance between them increases. *** P < 0.001.

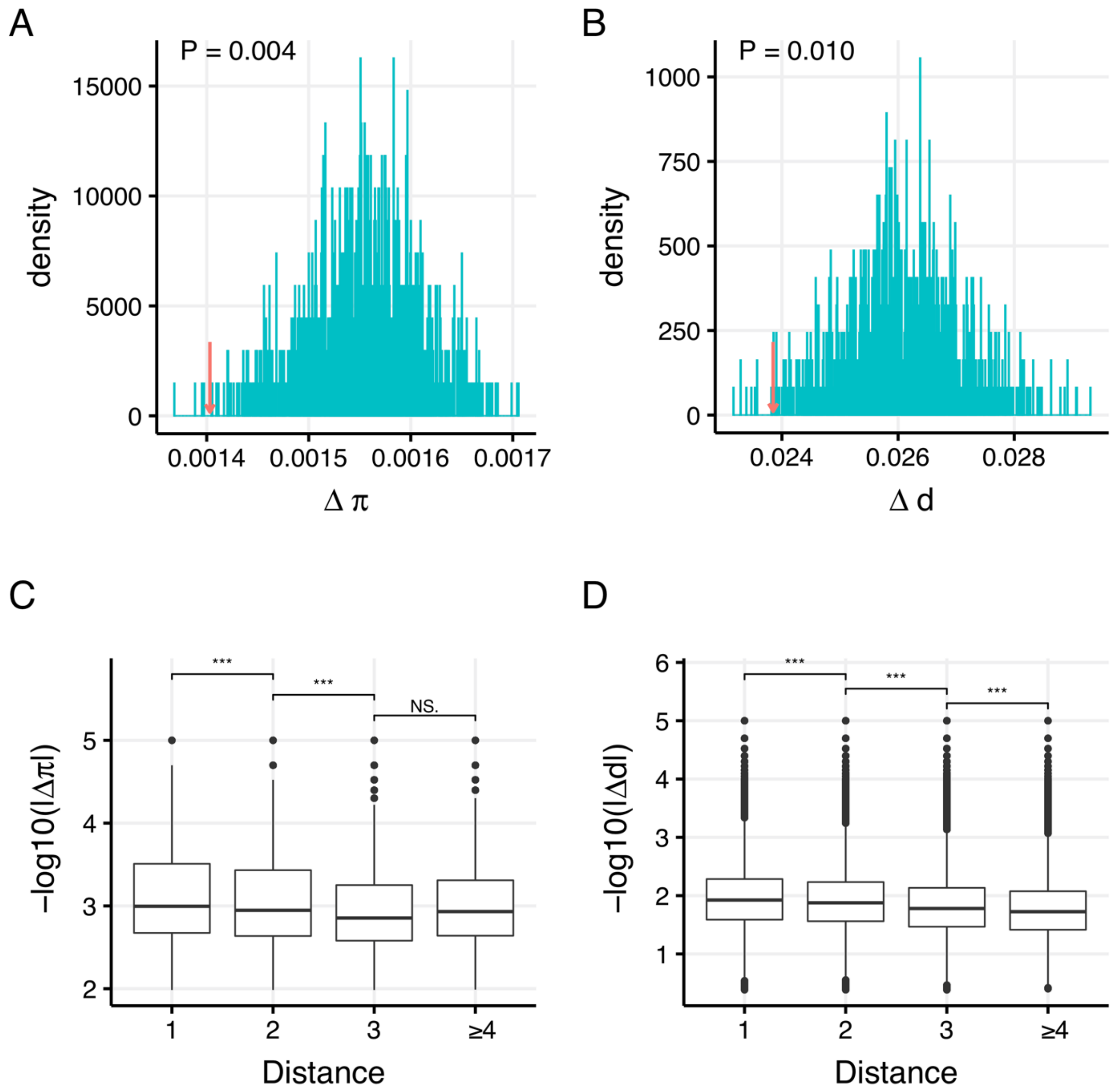

**Figure S30.** Correlated evolution between 3D interacting fragments in rice at the 100 kb scale. The upper two panels show the distribution of mean difference in genetic diversity (A) and evolutionary rates (B) between adjacent fragments of 1,000 simulated networks using CCS model. The blue lines show the simulated distribution of a parameter, while the red arrow represents the observed mean difference of a parameter. The lower two panels show that the similarity of fragment pairs decreases when the 3D distance between them increases in terms of genetic diversity (C) and evolutionary rates (D). *** $P < 0.001$, NS: nonsignificant.
